# Oxytocin receptor function regulates neural signatures of pair bonding and fidelity in the nucleus accumbens

**DOI:** 10.1101/2024.06.23.599940

**Authors:** Kimberly L. P. Long, Nerissa E. G. Hoglen, Alex J. Keip, Robert M. Klinkel, DéJenaé L. See, Joseph Maa, Jenna C. Wong, Michael Sherman, Devanand S. Manoli

## Abstract

The formation of enduring relationships dramatically influences future behavior, promoting affiliation between familiar individuals. How such attachments are encoded to elicit and reinforce specific social behaviors in distinct ethological contexts remains unknown. Signaling via the oxytocin receptor (Oxtr) in the nucleus accumbens (NAc) facilitates social reward as well as pair bond formation between mates in socially monogamous prairie voles^1–9^. How Oxtr function influences activity in the NAc during pair bonding to promote affiliative behavior with partners and rejection of other potential mates has not been determined. Using longitudinal *in vivo* fiber photometry in wild-type prairie voles and those lacking Oxtr, we demonstrate that Oxtr function sex-specifically regulates pair bonding behaviors and associated activity in the NAc. Oxtr function influences prosocial behavior in females in a state-dependent manner. Females lacking Oxtr demonstrate reduced prosocial behaviors and lower activity in the NAc during initial chemosensory investigation of novel males. Upon pair bonding, affiliative behavior with partners and neural activity in the NAc during these interactions increase, but these changes do not require Oxtr function. Conversely, males lacking Oxtr display increased prosocial investigation of novel females. Using the altered patterns of behavior and activity in the NAc of males lacking Oxtr during their first interactions with a female, we can predict their future preference for a partner or stranger days later. These results demonstrate that Oxtr function sex-specifically influences the early development of pair bonds by modulating prosociality and the neural processing of sensory cues and social interactions with novel individuals, unmasking underlying sex differences in the neural pathways regulating the formation of long-term relationships.

## Introduction

Long-term attachments between individuals are one of the most intriguing forms of social behavior and are central to human interactions, from parent-child bonds to enduring relationships between mates^10–12^. Despite the importance of attachment for the organization of complex social structures across species^13,14^, little is known about the neural mechanisms mediating these behaviors. Seminal work in socially monogamous prairie voles revealed that oxytocin and the oxytocin receptor (Oxtr) are key modulators of pair bonding, *i.e.*, the formation of selective and enduring attachments between mates. Pair bonded animals demonstrate both a preference for a bonded partner (partner preference) and active rejection of novel potential mates (strangers)^15^. Exogenous oxytocin in the brain facilitates the formation of partner preference in both male and female prairie voles^16,17^, while pharmacological inhibition of Oxtr disrupts the display of partner preference after mating^18^. Defining the neural circuits that govern pair bonding and determining how they are regulated by Oxtr are key to understanding social attachment behaviors. We recently demonstrated that, strikingly, prairie voles lacking Oxtr display partner preference^19^. However, Oxtr mutants display delayed development of partner preference and increased prosocial behavior towards strangers, suggesting that Oxtr influences the patterns of social interactions that facilitate pair bonding and controls the rejection of strangers^20^.

A key site of oxytocin action in the brain is the nucleus accumbens (NAc). The NAc has long been implicated in the reinforcement of behaviors ranging from addiction to innate displays associated with social interactions, including mating, aggression, and reciprocal interactions that mediate enduring attachments between mates^1,2,21–25^. The NAc integrates input from regions including the prefrontal cortex, thalamus, and amygdala as well as dopaminergic input from the ventral tegmental area to regulate diverse functions associated with reward- and survival-related behaviors^21,26–29^. Neuromodulatory signals -- including oxytocin from the paraventricular nucleus of the hypothalamus and serotonin from the dorsal raphe -- are also integrated within the NAc to influence prosocial behaviors and social reward^3,4,30–33^. In parallel to the reinforcement of rewarding stimuli, subregions of the NAc also appear to mediate responses to aversive stimuli, suggesting that components of the mesocorticolimbic system may control prosocial as well as agonistic interactions between individuals^34,35^. Compared to closely related but promiscuous vole species, prairie voles exhibit dramatically enriched oxytocin binding in the NAc^5,36,37^, and knockdown of Oxtr expression specifically within the NAc disrupts pair bonding^6^. Neuronal activity in the NAc in prairie voles evolves as a pair bond develops, such that ensembles responding to partner approach expand in size over time^38^. However, it remains unclear how the NAc responds to complex social interactions in the context of attachment and how such activity is modulated by Oxtr signaling.

Here, we utilize *in vivo*, longitudinal fiber photometry to examine NAc calcium activity across pair bond development in male and female, wild-type (WT) and Oxtr null (Oxtr^1-/-^) prairie voles. We demonstrate that the NAc responds to various types of social interaction and that Oxtr signaling regulates both pair bonding behaviors and behavior-related NAc activity in a sex-specific manner. Together, these results demonstrate that Oxtr regulates the development of pair bonds by modulating prosociality and neural processing of sensory and social interactions with novel individuals.

## Results

### Oxtr regulates NAc neural responses of naïve females to novel males

Our recent findings open questions about the precise role of Oxtr function for the formation of a pair bond^19^. To test the effects of Oxtr function on neural activity in the NAc during pair bonding and attachment behaviors, we implemented fiber photometry in WT and Oxtr^1-/-^ voles of both sexes. We examined NAc activity associated with social interactions during and after the course of pair bond formation (Fig. 1a). Specifically, these assays included introduction to a WT, opposite sex mate (partner); a partner preference assay; mating following estrus induction; acute separation from and reunification with the partner; and exposure to a novel, WT, sexually naïve, opposite sex animal (stranger). Importantly, this sequence allows us to compare dyadic social interactions with novel and familiar animals before and after bond formation^20^.

**Figure 1:**
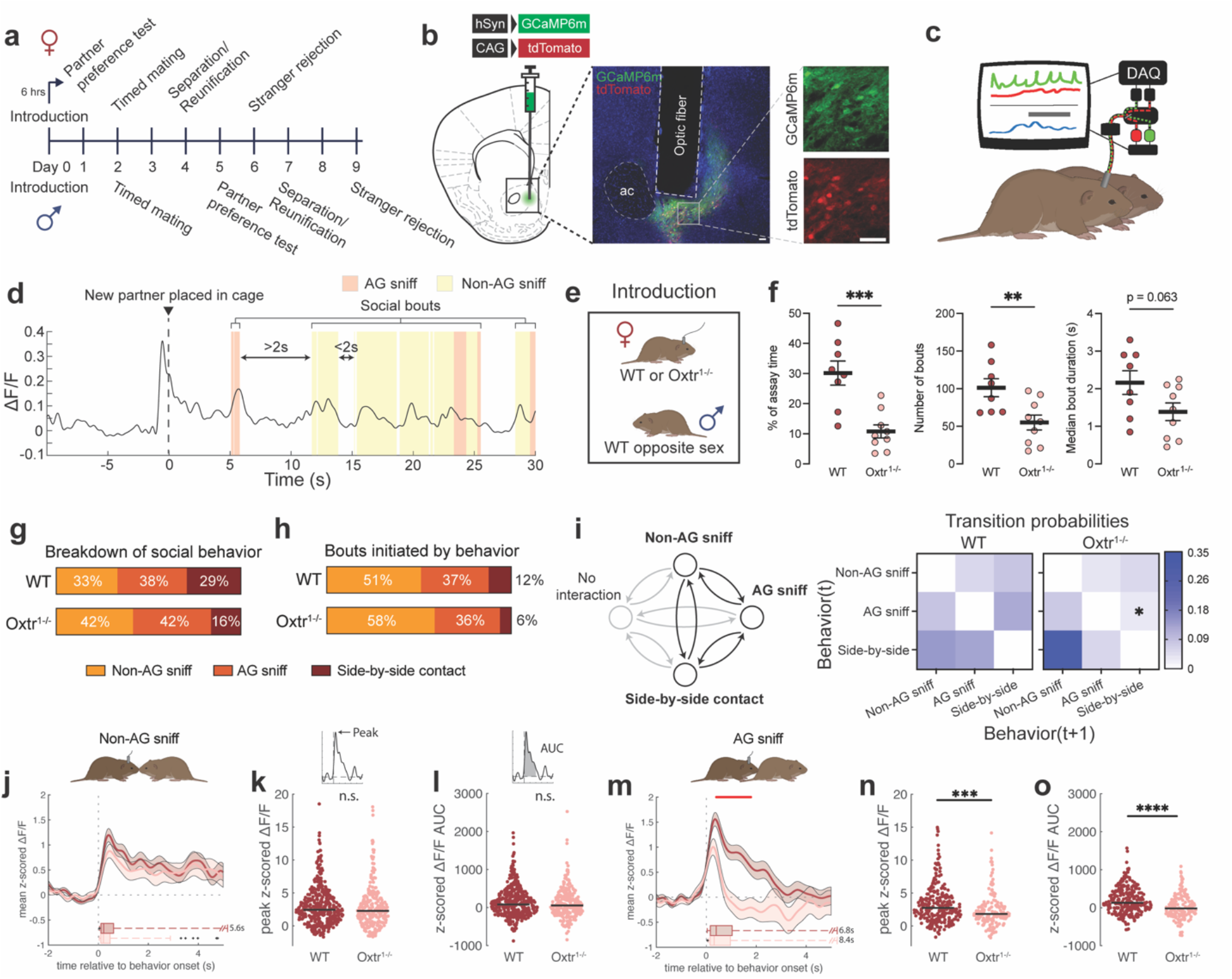
Loss of Oxtr in naive females disrupts NAc neural responses to novel males. **a**, Timeline of assays. **b**, Injection and fiber targeting in the medial NAc (scale bar, 50um). **c**, Photometry setup to image GCaMP6m during dyadic interactions in freely-moving prairie voles. **d**, Example GCaMP6m trace at the start of the introduction, with behavior events overlaid (anogenital [AG] sniff, orange; non-AG sniff, yellow). Behavior events separated by less than 2 seconds were considered part of a single social bout. **e**, Introduction procedure. A wild-type (WT) or Oxtr^1-/-^ female was placed into a clean cage, and a WT male was then introduced. **f**, Total percent of assay time engaged in (left), number of (middle), and median duration of (right) social bouts exhibited by females (for all plots, WT n=8, Oxtr^1-/-^ n=9 voles). **g**, Mean breakdown of social contact by the percentage of contact time engaged in AG sniffing, non-AG sniffing, and side-by-side contact. **h**, Percentages of social bouts initiated with non-AG sniff, AG sniff, or side-by-side contact. **i**, Left, schematic of Markov modeling of behavior, focusing on transitions from one social behavior to another. Right, heat maps of transition probabilities. **j**, Mean peri-event time histogram (PETH) of z-scored GCaMP6m ΔF/F by genotype aligned to the onset of non-AG sniffs (WT n=391 traces; Oxtr^1-/-^ n=272 traces). At the base of the plot is an adjusted boxplot of the durations of the initiating non-AG sniff. **k**, Swarm plot of peak z-scored ΔF/F values following non-AG sniffs. **l**, Area under the curve (AUC) values from z-scored ΔF/F traces following non-AG sniffs. **m**, Mean PETH by genotype aligned to AG sniffs. The red line indicates time points at which mean z-scored ΔF/F differ between WT and Oxtr^1-/-^ females (WT n=268 traces; Oxtr^1-/-^ n=161 traces). **n**, Peak z-scored ΔF/F values. **o**, AUC values from z-scored ΔF/F traces. Detailed statistics are presented in Extended Data File 1. *p<0.05, **p<0.01, ***p<0.001, ****p<0.0001. ac, anterior commissure; AG, anogenital; WT, wild-type; AUC, area under the curve.

We virally expressed the fluorescent calcium indicator GCaMP6m under the synapsin promoter in the medial NAc core and shell and implanted an optic fiber over the site of injection (Fig. 1b, Extended Data Fig. 1). We examined calcium activity within the NAc as WT or Oxtr^1-/-^ voles of either sex freely interacted with a stimulus animal and engaged in specific social interactions, such as chemosensory investigation, affiliation, mating, aggression, and defensive behaviors (Fig. 1c; Table 1). We then extracted z-scored GCaMP6m fluorescence traces surrounding individual social bouts (social touch preceded by at least 2 seconds of no interaction, Fig. 1d) and generated peri-event time histograms.

**Table 1:** All behaviors scored during dyadic interactions and their definitions.

| Behavior | Description/Criteria |
| --- | --- |
| Non-anogenital (non-AG) sniff | Social touch in which the snout of the implanted animal makes contact with face or flank, but not the anogenital zone (see below) of the stimulus animal. |
| Anogenital (AG) sniff | Social touch in which the snout of the implanted animal makes contact with the anogenital region (closest to the tail and the surrounding hindquarters, excluding the spine and back). |
| Side-by-side contact | Social touch in which more than 50% of the implanted animal's flank is in contact with the stimulus animal. |
| Social bout | A sequence of non-AG sniffing, AG sniffing, and/or side-by-side contact separated by less than 2 seconds of no interaction. |
| Mounting | A male vole approaches from behind and places its forequarters on the hindquarters of a female vole. The start of a mount was defined as the moment the male grasps the female's flank with its forepaws. A mount was scored for a female vole when she received mounting from a male. (Female mounting was extremely rare; we observed one instance out of all females in this study.) |
| Intromission | Pelvic thrusting of a male while in the mounting position. Intromission was scored for a female when a female received intromission. |
| Strike | A lunge, bite, or kick by the experimental vole directed towards the stimulus animal. |
| Defensive strike | A lunge or strike by the experimental vole following the receipt of aggression or social touch from a stimulus animal. |
| Tussle | A highly aggressive rolling fight in which both voles are biting and/or wrestling. |
| Aggression receipt | The receipt of a strike or defensive strike by the experimental vole from a stimulus vole. |
| Defensive rear | A defensive posture in which both forepaws are raised off the ground while oriented towards the stimulus vole. |

In mice and other rodents, including prairie voles, Oxtr signaling modulates prosocial behavior^4,7,17,32^. To interrogate the role of Oxtr during female prairie voles’ initial interactions with a mate, we examined patterns of behavior and activity in the NAc during the introduction of a naïve female to a naïve WT male partner (introduction, Fig. 1e). Compared to WT females, Oxtr^1-/-^ females exhibited less investigative and affiliative social interaction with males, reflected in both the total amount of time spent in social interactions and the number of social bouts initiated (Fig. 1f, Extended Data Fig. 2a). Chemosensory investigation (*i.e*., sniffing), including anogenital sniffs and sniffs directed to other parts of the body, comprised a large proportion of these social interactions and frequently preceded other behaviors (Fig. 1g,h). Markov chain modeling of female behavior revealed that Oxtr^1-/-^ females were also less likely than WT females to transition from anogenital investigation to side-by-side contact, a highly prosocial and affiliative behavior that increases with pair bonding (Fig. 1i, Extended Data Fig. 2b-d). Thus, loss of Oxtr decreases social and affiliative displays by females to a novel partner.

We next tested whether these changes in behavior in Oxtr^1-/-^ females were accompanied by changes in neural activity within the NAc. Activity during initial social interactions with males was decreased in Oxtr^1-/-^ females compared to WT females in a behavior-specific pattern. Oxtr^1-/-^ females exhibited decreased peak fluorescence and area under the curve (AUC) at the onset of anogenital investigations of males (Fig. 1j-o). In contrast, activity associated with non-anogenital sniffs or bouts initiated by side-by-side contact did not differ between WT and Oxtr^1-/-^ females (Fig 1j-l, Extended Data Fig. 2e-l). The specificity of this difference to chemosensory investigation suggests that Oxtr regulates neural processing of male chemosensory cues in naïve female prairie voles^39–42^ and that disruptions of these responses may contribute to changes in prosocial behavior towards novel males.

### Oxtr modulates neural and behavioral responses to novel males in a state-dependent manner in female prairie voles

After establishing a role for Oxtr in regulating naïve female responses to novel males, we examined the effects of Oxtr on the response of bonded females to familiar (partner) and novel (stranger) males. Pair bonding results in increased prosocial and affiliative (huddling) behavior with partners and a dramatic switch to agonistic (rejection) behavior directed towards strangers^14,20^. We tested how changes in bonding state affect behavioral and neural responses to a male partner by comparing female responses at an early stage of bonding to responses to the same male after pair bond formation (Extended Data Fig. 3a, Day 0: Introduction vs. Day 4: Reunion). Consistent with previous studies, WT females exhibited reduced anogenital investigation of a partner during reunion compared to their first encounter (Extended Data Fig. 3b,c). Oxtr^1-/-^ females showed low levels of anogenital investigation regardless of bonding status, and WT and Oxtr^1-/-^ females did not differ in levels of investigation of a partner following bonding. Similarly, both WT and Oxtr^1-/-^ females spent more time in affiliative side-by-side contact with their familiar partner during reunion compared to the same male during the introduction (Extended Data Fig. 3d). Broadly, WT and Oxtr^1-/-^ females exhibited similar patterns of behavior and activity in the NAc after bonding. Activity in the NAc of both WT and Oxtr^1-/-^ females was higher during non-anogenital investigations of familiar partners at the time of reunion than of novel partners during introduction (Extended Data Fig. 3e-g), suggesting that interactions with familiar partners elicit greater activity in the NAc after bonding. Furthermore, WT and Oxtr^1-/-^ females did not differ in levels of activity in the NAc during anogenital investigations of partners following bonding (Extended Data 3h-q). This indicates that Oxtr is not required for the changes in activity in the NAc during the investigation of partners that result from pair bond formation.

We next examined the responses of females to a novel male before and after bonding (Extended Data Fig 3a, Day 0: Introduction vs. Day 6: Stranger rejection). WT females exhibited reduced anogenital investigation of a novel male following bonding (stranger), compared to when they were first paired with a novel male (partner). In contrast, Oxtr^1-/-^ females showed consistently low levels of chemosensory investigation over the course of pair bonding. Furthermore, WT and Oxtr^1-/-^ females did not differ in their neural responses to stranger males after bonding. Thus, loss of Oxtr specifically affects activity in the NAc during females’ interactions with novel males early in bonding and is not required for bonding-associated changes in NAc activity during anogenital investigation of a partner or a novel stranger.

Mating accelerates pair bond formation in prairie voles^43,44^. To understand the effects of Oxtr on behavior and activity in the NAc during the early stages of bonding, we examined activity in females during mating. Loss of Oxtr did not affect females’ mating or social behaviors with the partner 48 hours after animals were first introduced to each other and after estrus was induced and synchronized across pairs (Extended Data Fig. 4a-e). During mounting attempts, activity within the NAc gradually decreased below baseline, and this decrease was greater during successful mounting attempts in which the male was able to proceed to intromission compared to attempts after which intromission did not occur (Extended Data Fig. 4f-k). Furthermore, activity within the NAc during most interactions did not differ between WT and Oxtr^1-/-^ females, except for side-by-side contacts, during which Oxtr^1-/-^ females showed higher levels of activity (Extended Data Fig. 4l-w). These results suggest that Oxtr exerts more significant effects on activity in the NAc of female voles at the earliest stages of bonding and that its effects are highly dependent on the bonding state.

The selective preference for a partner over a stranger is a hallmark of pair bonding^43,45^. A few hours of cohabitation with a novel male are sufficient for female prairie voles to form a pair bond, reflected in the display of partner preference, *i.e*. a preference to engage in social interaction and side-by-side contact with their partner rather than a novel, potential mate^20,45^. We therefore tested whether the loss of Oxtr impacts partner preference at the earliest point at which WT females display such behavior. We placed females in a 3-chamber arena six hours after initial introductions and allowed them to choose between interacting with their partner or a male stranger (Fig. 2a). We found that loss of Oxtr did not disrupt female displays of partner preference. All females, regardless of genotype, interacted significantly more with their partner than with the stranger (Fig. 2b; Extended Data Fig. 5a,b) and displayed a clear preference to engage in affiliative side-by-side contact with their partner (Fig. 2c). Neural responses during interactions with the partner were also similar between WT and Oxtr^1-/-^ females (Fig. 2f-h). This contrasts with the differences observed during females’ first encounter with the same partner male, again suggesting that Oxtr function is not required for bonding-associated changes in neural responses to partners.

**Figure 2:**
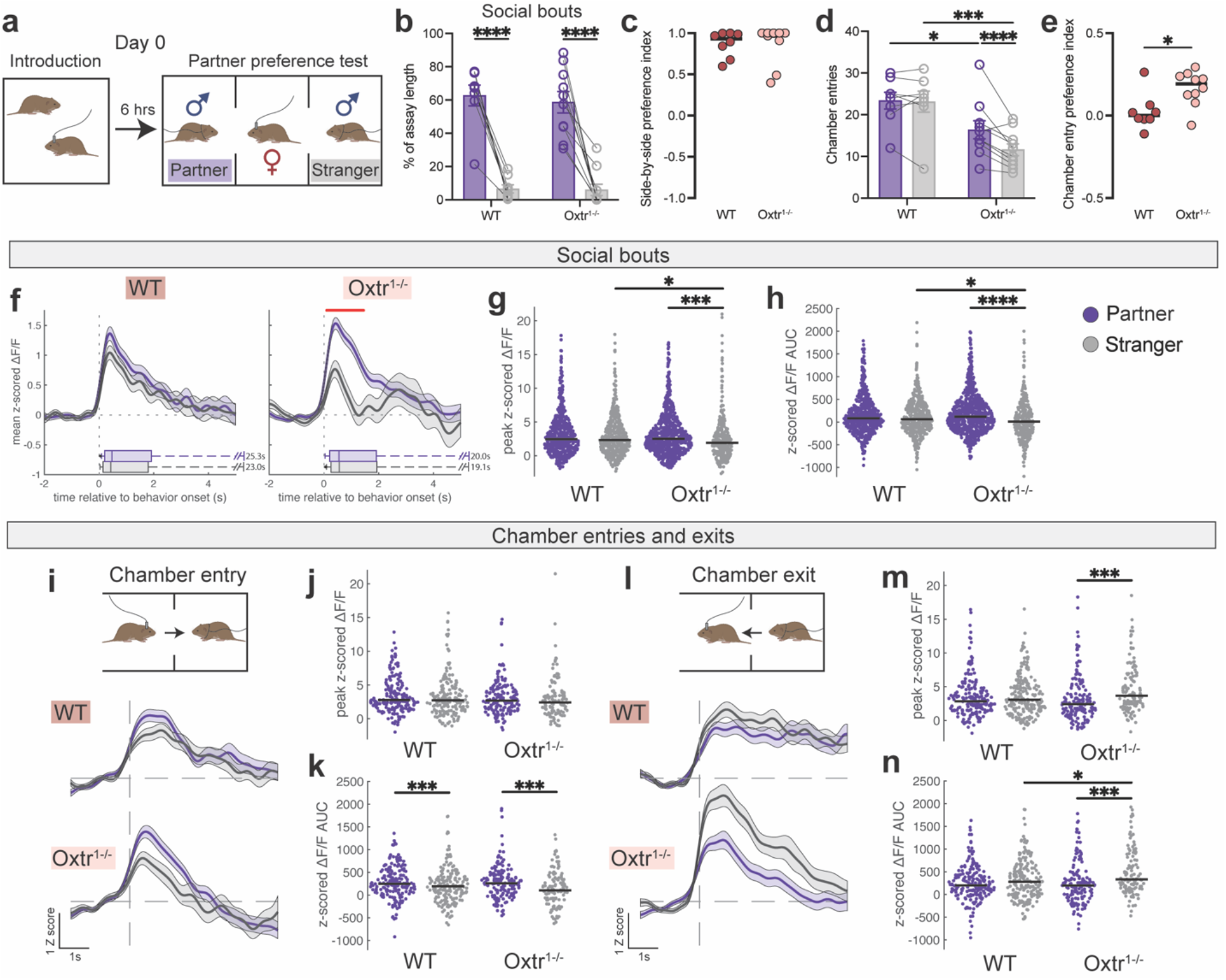
Loss of Oxtr disrupts female neural and behavioral responses to stranger males. **a**, Timeline and schematic of the introduction and partner preference test (PPT) for females. **b**, Percent of assay time spent engaged in social bouts with either the partner (purple) or the stranger (gray) (for all plots, WT n=8, Oxtr^1-/-^ n=10 voles). **c**, Side-by-side contact preference index scores. Preference index scores of 1 indicate exclusive side-by-side contact with the partner, and -1 with the stranger. **d**, Number of entries to the partner or stranger chambers. **e**, Chamber entry preference index scores. **f**, Mean PETH of z-scored GCaMP6m ΔF/F by stimulus animal aligned to the onset of social bouts (WT_Partner_ n=539 traces; WT_Stranger_ n=444 traces; Oxtr^1-/-^_Partner_ n=732 traces; Oxtr^1-/-^_Stranger_ n=304 traces). At the base of the plot is an adjusted boxplot of the durations of the initiating behavior. The red line indicates time points at which mean z-scored ΔF/F differs between partner and stranger-related activity in Oxtr^1-/-^ females. **g**, Swarm plot of peak z-scored ΔF/F values. **h**, AUC values from z-scored ΔF/F traces. **i**, Left, mean ΔF/F PETH aligned to entries to either the partner chamber or stranger chamber (WT_Partner_ n=184 traces; WT_Stranger_ n=182 traces; Oxtr^1-/-^_Partner_ n=160 traces; Oxtr^1-/-^_Stranger_ n=115 traces). **j**, Peak z-scored ΔF/F values. **k**, AUC values from z-scored ΔF/F traces. **l**, Left, mean ΔF/F PETH aligned to exits from either the partner chamber or stranger chamber (WT_Partner_ n=187 traces; WT_Stranger_ n=195 traces; Oxtr^1-/-^_Partner_ n=150 traces; Oxtr^1-/-^_Stranger_ n=121 traces). **m**, Peak z-scored ΔF/F values. **n**, AUC values from z-scored ΔF/F traces. Detailed statistics are presented in Extended Data File 1. *p<0.05, **p<0.01, ***p<0.001, ****p<0.0001. WT, wild-type; PETH, peri-event time histogram; AUC, area under the curve.

While we did not observe a difference in females’ interactions with their partner between WT and Oxtr^1-/-^ animals during tests of partner preference, we observed differences in both behavioral and neural responses to stranger males in Oxtr^1-/-^ females. In contrast to WT females, which all showed at least two bouts of side-by-side contact with strangers, Oxtr^1-/-^ females avoided stranger males (Extended Data Fig. 5c). Furthermore, Oxtr^1-/-^ females were less likely to enter the stranger male’s chamber (Fig. 2d-e). Neural responses to stranger males reflected these different behavioral response patterns in Oxtr^1-/-^ females. Both peak and AUC of calcium activity in the NAc during interactions with the stranger male were significantly decreased in Oxtr^1-/-^ females when compared to both their own interactions with partners as well as WT females’ interactions with strangers (Fig. 2f-h, Extended Data Fig. 5d-l). Entry into the partner chamber elicited greater total activity (AUC) than entry into the stranger chamber, regardless of genotype (Fig. 2i); however, Oxtr^1-/-^ females exhibited greater NAc activity upon leaving the stranger chamber than WT females (Fig. 2j). Our findings suggest that Oxtr controls prosocial behavior and associated neural activity in the NAc in a state-dependent manner in females. In naïve females, Oxtr function promotes prosocial behavior towards novel males and associated increases in neural activity in the NAc. Strikingly, however, increases in prosocial behavior, huddling behavior, and activity in the NAc following pair bonding occur independent of Oxtr function.

### Oxtr regulates NAc neural signatures of partner preference in male prairie voles

Previous studies, including our own recent findings, suggest that Oxtr function differs between the sexes and that loss of Oxtr impacts females and males in different ways^20,46^. We therefore examined NAc neural signatures of pair bonding in male prairie voles to determine if the influence of Oxtr function on pair bonding behavior and associated neural activity in the NAc also differs between sexes. We first tested the effects of Oxtr on NAc activity prior to bond formation during naïve males’ first encounter with a WT female. We found that, while Oxtr^1-/-^ males show increased prosocial behavior during early interactions with a female partner, they show no differences in activity in the NAc associated with these behaviors. During the first introduction to a naïve female partner, Oxtr^1-/-^ males displayed increased social investigation of females when compared to WT males, contrary to patterns we observed in Oxtr^1-/-^ females (Fig. 3a,b, Extended Data Fig. 6a-d). Moreover, Oxtr^1-/-^ males engaged in significantly less agonistic behavior (strikes) towards females than WT males, with no mutant males displaying strikes towards females (Extended Data Fig. 6e-h). In contrast to females, we found no differences in activity in the NAc between WT and Oxtr^1-/-^ males during social bouts or specific social interactions, including both anogenital and non-anogenital investigation (Fig. 3c-e, Extended Data Fig. 6i-q). Similarly, we found no differences between WT and Oxtr^1-/-^ males in behavior or NAc activity during mating (Extended Data Fig. 6r-dd). In naïve male prairie voles, Oxtr function thus appears to reduce prosocial investigation of novel females and facilitates agonistic displays but does not appear to modulate NAc activity under these conditions.

**Figure 3:**
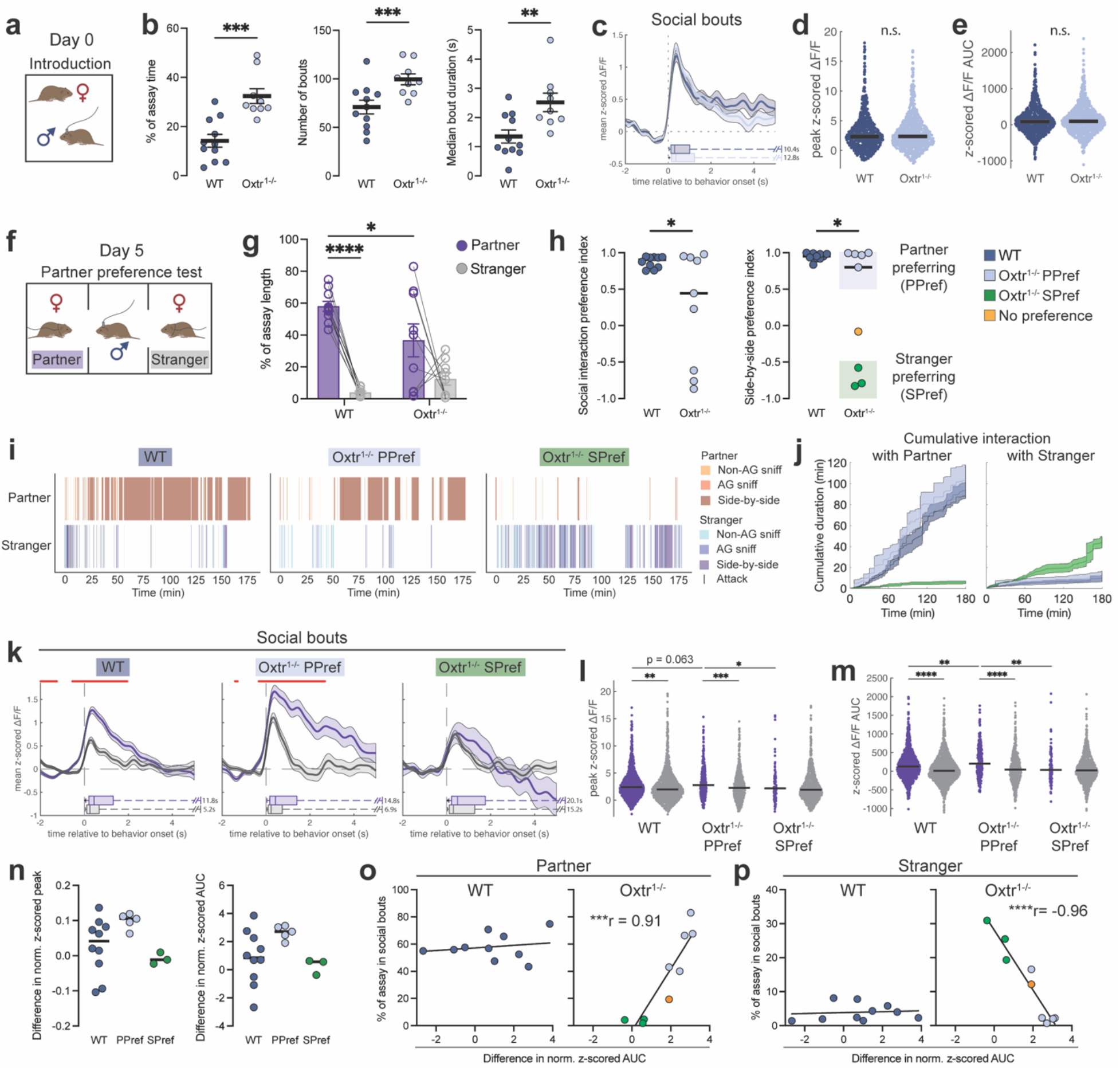
Oxtr regulates NAc neural signatures of partner preference in male prairie voles. **a**, Introduction of a WT or Oxtr^1-/-^ male to a WT female partner. **b**, Total percent of assay time engaged in (left), number of (middle), and median duration of (right) social bouts exhibited by males (WT n=11, Oxtr^1-/-^ n=9). **c**, Mean ΔF/F PETH by genotype aligned to the onset of social bouts (WT n=750 traces from 11 animals; Oxtr^1-/-^ n=865 traces from 9 animals). At the base of the plot is an adjusted boxplot of the durations of the initiating behavior. **d**, Peak z-scored ΔF/F values. **e**, AUC values from z-scored ΔF/F traces. **f**, Schematic of the PPT, conducted on day 5 in males. **g**, Percent of assay time spent engaged in social bouts with either the partner (purple) or stranger (gray) (WT n=10, Oxtr^1-/-^ n=9). **h**, Left, social interaction preference index scores. Right, side-by-side contact preference index scores. Preference for partner or stranger was determined by whether side-by-side preference index scores were greater than 0.5 (partner-preferring, PPref) or less than -0.5 (stranger-preferring, SPref). For all following plots, WT n=10, Oxtr^1-/-^ PPref n=5, Oxtr^1-/-^ SPref n=3 voles. **i**, Example behavior rasters from a WT, Oxtr^1-/-^ PPref, and Oxtr^1-/-^ SPref male. **j**, Mean (+/- s.e.m.) cumulative duration plots of social interaction with either the partner or stranger across the 3-hour assay. **k**, Mean ΔF/F PETH aligned to social bouts with either the partner or stranger. The red line indicates time points at which mean z-scored ΔF/F differ between partner- and stranger-related activity (WT_Partner_ n=932 traces and WT_Stranger_ n=1193 traces; Oxtr^1-/-^ PPref_Partner_ n=296 traces and PPref_Stranger_ n=500 traces; Oxtr^1-/-^ SPref_Partner_ n=111 traces and SPref_Stranger_ n=821 traces). **l**, Peak ΔF/F values. **m**, AUC values. **n**, Per animal difference (partner - stranger) between partner-elicited and stranger-elicited peak ΔF/F (left) and AUC (right). **o**, Correlations of partner-stranger AUC difference and percent of time spent in social interactions with the partner. **p**, Correlations of partner-stranger AUC difference and percent of time spent in social interactions with the stranger. Detailed statistics are present in Extended Data File 1. *p<0.05, **p<0.01, ***p<0.001, ****p<0.0001. PPT, partner preference test; WT, wild-type; PPref, partner-preferring; SPref, stranger-preferring; AUC, area under the curve; norm., Box Cox normalized.

We next examined the effects of Oxtr loss on neural activity in the NAc associated with pair bond formation in males. Compared to females, male prairie voles require longer periods of cohabitation and mating before they display robust preference for partners^20,45^. We therefore examined partner preference in males five days after introduction to WT females, a period of cohabitation that is sufficient for partner preference formation in WT males^20^ (Fig. 3f). In contrast to our observations in females, loss of Oxtr disrupted the display of partner preference in males even after 5 days of cohabitation (Fig. 3g,h). The difference in preference between populations of WT versus Oxtr^1-/-^ males was due to a subgroup of mutant males that strongly preferred interacting with a stranger female over their partner (Fig. 3h-j, Extended Data Fig. 7a,b). Approximately one half (5 out of 9) of Oxtr^1-/-^ males preferred to engage in side-by-side contact with the partner (index score >0.5, “partner-preferring”), while 3 out of 9 spent little time with the partner in favor of the stranger (index score <-0.5, “stranger-preferring,” Fig. 3h). We then analyzed activity in the NAc to determine whether these behaviorally defined subpopulations of Oxtr^1-/-^ males also differed in their neural responses in the NAc during social interactions. All males showed increased activity in the NAc upon entering the partner’s chamber versus that of the stranger and, inversely, increased activity when exiting strangers’ versus partners’ chambers (Extended Data Fig. 7c-f). Thus, approach towards partners (or departure from strangers) increases NAc activity in male prairie voles independent of Oxtr function, consistent with prior work^38^. However, activity differed between subgroups during direct social interactions. Both WT and partner-preferring Oxtr^1-/-^ males showed significantly higher levels of activity during social interactions with partners when compared to interactions with strangers (Fig. 3k-m, Extended Data Fig. 7g-p). Moreover, partner-preferring Oxtr^1-/-^ males showed higher levels of activity in the NAc during social interactions with partners compared to WT males (Fig. 3m). In contrast, stranger-preferring Oxtr^1-/-^ males showed lower levels of activity during interactions with partners when compared to partner-preferring Oxtr^1-/-^ males, and no differences in activity in the NAc when comparing interactions with partners or strangers. Thus, with the formation of partner preference in males, increased activity in the NAc during interactions with partners versus strangers occurs independent of Oxtr function but is further increased in the absence of Oxtr. In contrast, loss of Oxtr unmasks a population of males that fail to display partner preference and show no difference in NAc activity during interactions with partners versus strangers.

Given the intriguing difference in partner preference and activity in the NAc between subpopulations of Oxtr^1-/-^ males, we next tested whether this relationship is evident in WT males. Despite robust preference for partners over strangers and higher levels of activity associated with partner versus stranger interactions as a population, individual WT males showed large variation in the difference between partner- and stranger-associated activity in the NAc, which did not correlate with the amount of social interaction displayed with either female (Fig. 3n-p).

In contrast, Oxtr^1-/-^ males that displayed a preference for their partner tended to have a larger difference in partner-versus stranger-associated activity in the NAc when compared to mutant males that displayed a preference for strangers (p=0.064 for peak and p=0.077 for AUC, Fig. 3n). The difference between levels of activity in the NAc of Oxtr^1-/-^ males in response to partners or strangers strongly predicted the amount of social interaction displayed towards either female. Oxtr^1-/-^ males with less neural difference between partner and stranger interactions displayed a stronger preference for stranger females (Fig. 4o-p). These observations suggest that Oxtr function increases pair bonding-related behaviors in some males, including display of a partner preference as well as associated neural activity in the NAc, but is not necessary for the demonstration of partner preference or associated activity in others. Oxtr may therefore function during pair bond formation in males to reinforce both prosocial behaviors and increases in activity in the NAc with partners or, alternatively, to suppress prosocial behavior during interactions with strangers.

**Figure 4:**
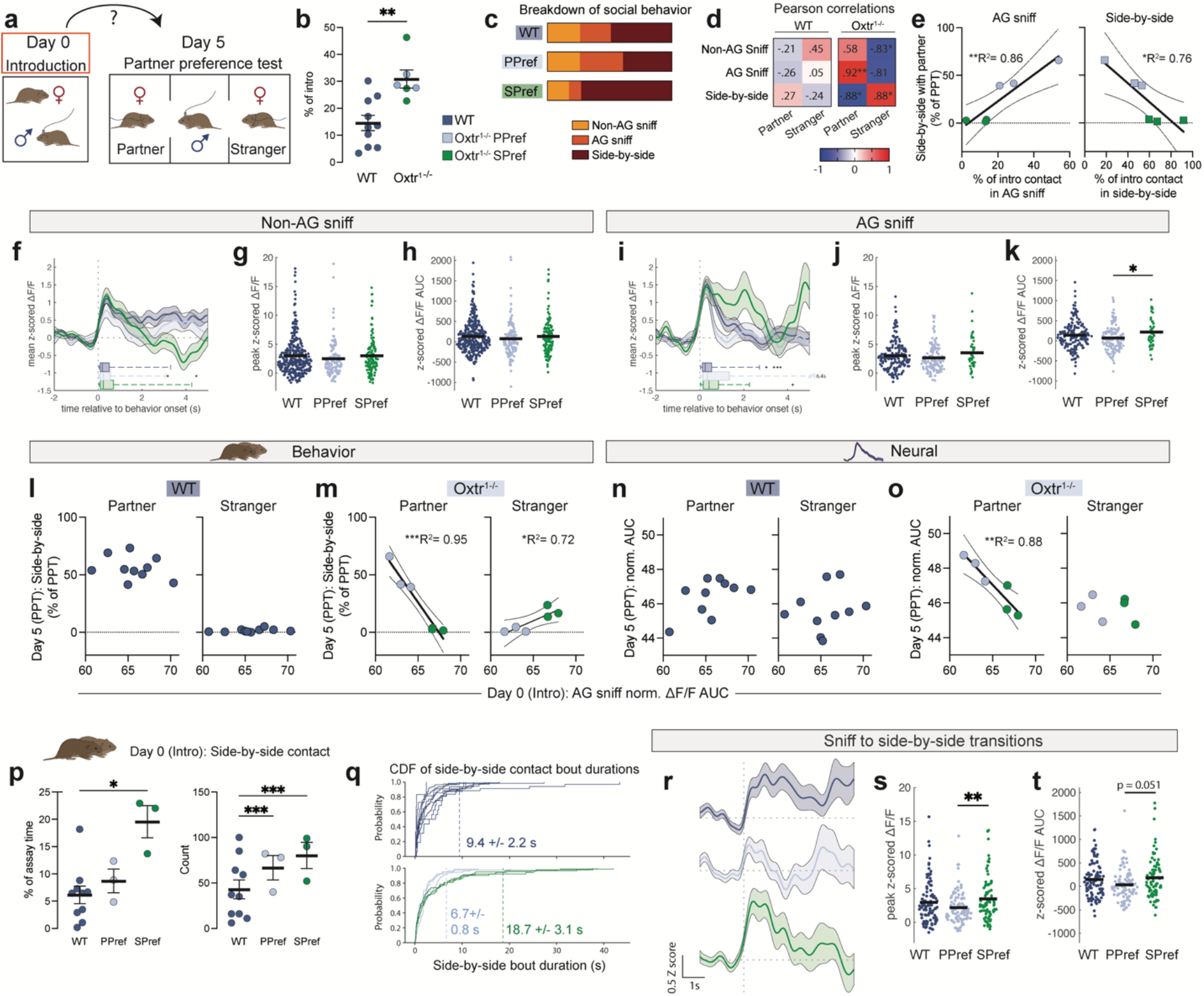
Oxtr regulates behavioral and neural trajectories of pair bonding in male prairie voles. **a**, Examination of behavior and neural activity from Day 0 (introduction) in relation to metrics from Day 5 (PPT). **b**, Percent of time engaged in social interaction with a newly partnered female during the introduction, plotting only animals from which data were successfully collected during both introduction and PPT. Individual animals are colored according to PPT behavior profile (for all plots, WT n=10, Oxtr^1-/-^ PPref n=3, Oxtr^1-/-^ SPref n=3). **c**, Mean breakdown of social interaction during the introduction by type of social touch. **d**, Heat maps of Pearson correlations between introduction behavior (% of contact time) and PPT partner or stranger side-by-side contact (% of assay time). **e**, Linear regression of introduction behavior to PPT behavior. X-axis: Percent of social touch during the introduction spent AG sniffing (left) or in side-by-side contact (right). Y-axis: Percent of PPT time spent in side-by-side contact with the partner. **f**, Mean NAc ΔF/F PETH aligned to non-AG sniffs during the introduction (WT n=338 traces; Oxtr^1-/-^ PPref n=138 traces; Oxtr^1-/-^ SPref n=123 traces). At the base of the plot is an adjusted boxplot of the durations of the initiating behavior. **g**, Peak z-scored ΔF/F values. **h**, AUC values from z-scored ΔF/F traces. **i**, Mean ΔF/F PETH aligned to AG sniffs during the introduction (WT n=187 traces; Oxtr^1-/-^ PPref n=129 traces; Oxtr^1-/-^ SPref n=42 traces). **j**, Peak z-scored ΔF/F values. **k**, AUC values from z-scored ΔF/F traces. **l-o**, Linear regressions comparing AG sniff-related NAc activity and PPT behavior or neural data. X-axis: normalized (norm.) AUC at the onset of AG sniff bouts during the introduction, averaged by animal. Y-axis: PPT side-by-side contact (l-m) or PPT normalized AUC surrounding social bouts, averaged by animal (n-o). **p**, Left, percent of introduction time spent in side-by-side contact. Right, number of side-by-side contact events during the introduction. **q**, Cumulative distribution functions (CDF) for side-by-side contact bout durations per animal. Dotted lines show the mean (+/- s.e.m.) of the 95th percentile values from each group. **r**, Mean PETH aligned to transitions from sniffing to side-by-side contact, centered at the onset of the sniff (no filtering for behavior in the 2 seconds prior to sniff onset; WT n=97 traces; Oxtr^1-/-^ PPref n=98 traces, Oxtr^1-/-^ SPref n=82 traces). **s**, Peak z-scored ΔF/F values. **t**, AUC values from z-scored ΔF/F traces. Detailed statistics are presented in Extended Data File 1. *p<0.05, **p<0.01, ***p<0.001, ****p<0.0001. WT, wild-type; PPref, Oxtr^1-/-^ partner-preferring; SPref, Oxtr^1-/-^ stranger-preferring; AG, anogenital; PPT, partner preference test; AUC, area under the curve; norm., Box Cox normalized; CDF, cumulative distribution function.

### Oxtr regulates behavioral and neural trajectories of pair bonding in male prairie voles

Loss of Oxtr reveals distinct populations of males that strongly prefer either a partner or stranger. We therefore analyzed the behavior of these populations at different stages of pair bonding to determine if the trajectories of social behaviors or patterns of activity in the NAc differed between these groups or between WT and Oxtr^1-/-^ males. We found that changes in behavior and NAc activity during pair bond formation in males can occur independent of Oxtr function. We compared chemosensory investigation (anogenital and non-anogenital) between the first encounter with female partners (introduction) and either reunion with these partners or encounters with novel strangers during stranger rejection and found no differences between WT and Oxtr^1-/-^ males (Extended Data Fig. 8a-d). All males displayed increased side-by-side contact with the familiar partner upon reunion compared to the naïve partner during the introduction. Notably, stranger-preferring Oxtr^1-/-^ males did not display higher levels of agonistic behavior towards their partners (Extended Data Fig. 8e-h). Regardless of Oxtr function, activity in the NAc was greater during non-anogenital sniffs of female partners upon reunion compared to the introduction (Extended Data Fig. 8i-n). This suggests that, as with females, cohabitation enhances specific partner-associated activity in the NAc in males independent of Oxtr function.

Given the emergence of distinct populations of Oxtr^1-/-^ males that strongly prefer partners or strangers, we tested whether these populations might be distinguishable at the initial stages of pair bonding. We examined male behavior and neural activity during the first interactions with a partner to determine if patterns of behavior or activity in the NAc could predict the future preference for partners or strangers (Fig. 4a). Naïve Oxtr^1-/-^ males engage in more prosocial interactions when first introduced to a female (Fig. 4b). We found no difference in the total amount of social behaviors when we compared partner-versus stranger-preferring Oxtr^1-/-^ males (Fig. 4b, Extended Data Fig. 8o-q). However, examining the specific patterns of behavior during social interactions revealed significant differences between these populations. Stranger-preferring Oxtr^1-/-^ males tended to spend more of their interaction time engaged in highly affiliative side-by-side contact compared to partner-preferring Oxtr^1-/-^ males, who spent more time engaged in anogenital investigation (Fig. 4c, Extended Data Fig. 8p). Moreover, the amount of time Oxtr^1-/-^ males engaged in anogenital investigation and side-by-side contact during initial interactions with a partner significantly correlated, in opposing directions, with levels of partner- and stranger-directed side-by-side contact when given a choice between either female (Fig. 4d, Extended Data Fig. 8r,s). Thus, behavior exhibited by Oxtr^1-/-^ males during the first 30 minutes of interaction with a female can predict patterns of partner preference 5 days later (Fig. 4e, Extended Data Fig. 8t-x), indicating that the two subpopulations are immediately identifiable based on their behavior.

Stranger-preferring Oxtr^1-/-^ males differ from partner-preferring Oxtr^1-/-^ and WT males in both their amount of anogenital investigation and side-by-side contact (Fig. 4c). We examined whether these behavioral differences were associated with differences in activity in the NAc during social interactions. During the earliest interactions with a partner, activity in the NAc associated with anogenital investigation was greater in Oxtr^1-/-^ males that went on to prefer strangers compared to those that went on to prefer partners (Fig. 4i-k). In contrast, activity in the NAc associated with non-anogenital investigation was similar across all males. Consistent with our behavioral observations, the mean neural activity following an anogenital sniff during initial interactions with a female strongly correlated with affiliative behavior displayed towards partners *and* strangers five days later in individual Oxtr^1-/-^, but not WT, males. (Fig. 4l-m). Neural activity in the NAc of Oxtr^1-/-^ males was strongly inversely correlated between early and late interactions with the same female. Specifically, partner-preferring Oxtr^1-/-^ males showed lower levels of activity following anogenital investigations during the introduction and higher levels of activity during social interactions with partners 5 days later. In contrast, stranger-preferring Oxtr^1-/-^ males displayed the opposite pattern of activity (Fig. 4n). We observed no correlation between early and future partner-related activity in WT males (Fig. 4n,o). Thus, in the absence of Oxtr function, early anogenital investigation of novel females and associated neural activity in the NAc may rapidly influence how males bond during cohabitation and even predict future preference.

We observed that stranger-preferring Oxtr^1-/-^ males spent significantly more time in side-by-side contact and longer bouts of this contact with partner females during initial interactions than both WT males and partner-preferring Oxtr^1-/-^ males (Fig. 4p,q). We analyzed the dynamics of and relationship between prosocial side-by-side contact, activity within the NAc, and future preference for partners or strangers. In rodents, social interactions are typically initiated by chemosensory investigations that inform animals of each other’s identity (*e.g*., species, sex, health, reproductive status, etc.)^47,48^. Consistent with this observation, the majority (73.8%) of side-by-side events during introductions occurred following sniff investigations. Therefore, we examined NAc calcium activity at the transition from sniffs to bouts of side-by-side contact. As with anogenital investigations alone, the transition from such sniffs to side-by-side contact during initial interactions was associated with higher peak activity in the NAc in future stranger-preferring Oxtr^1-/-^ males (Fig. 4r-t). Similarly, mean NAc neural responses at the onset of such transitions were strongly correlated with patterns of preference for partners or strangers 5 days later in Oxtr^1-/-^, but not WT, males (Extended Data Fig. 8y-bb). The dynamics of chemoinvestigation and prosocial behaviors and the associated neural activity in the NAc during the first interactions of Oxtr^1-/-^ males with a novel female can therefore predict future preference for and neural responses to partners and strangers. In the absence of Oxtr, higher levels of activity in the NAc following anogenital investigations correlate with increased prosocial behavior during males’ initial interactions with a partner female, but a preference for a stranger female in the future. In contrast, lower levels of NAc activity upon introduction correlate with increased chemoinvestigation and fewer initial prosocial behaviors, but a robust later preference for these females. Thus, from the first interaction between a male vole and his female partner, Oxtr function modulates neural responses to chemosensory information towards a common pair bonded outcome. Loss of Oxtr unmasks a population of males that show a robust stranger preference, even after prolonged cohabitation with a partner. These findings suggest that Oxtr function works to stabilize a range of sensory responses and social behaviors towards common and less-varied patterns of interactions to promote social monogamy in male prairie voles.

## Discussion

Adult attachments are comprised of nuanced and finely tuned behaviors under complex neural and hormonal control. To examine neural activity associated with the formation of long-term social attachments, we implemented fiber photometry in the NAc, a key node in the circuitry that mediates pair bonding in socially monogamous prairie voles, and examined activity in this region across pair bond formation and related attachment behaviors. We compared activity between wild-type animals and those lacking Oxtr in both males and females. We found that activity in the NAc during interactions with partners diverges from that with strangers over the course of pair bonding. Loss of Oxtr has opposing effects on specific pair bond-related social behaviors and associated NAc activity between males and females. Strikingly, in Oxtr^1-/-^ males, behavior and activity in the NAc during the first interactions with potential mates strongly predicts future preference for partners or strangers 5 days later. Critically, these interrogations would not have been possible without the integration of *in vivo* calcium recordings, molecular genetics, and ethological approaches^49,50^.

Sex differences within the NAc may arise from intrinsic properties of neurons within this region to influence activity within the NAc in sex-specific ways^51–66^. We recently demonstrated that loss of Oxtr unmasks sex differences in gene expression in the NAc that are not found in wild-type animals^20^. Thus, the absence of Oxtr signaling during development may also contribute to the differences we observed in patterns of activity in the NAc between male and female Oxtr^1-/-^ voles^51–57^. Alternatively, or in combination, such sex differences in the NAc may arise from Oxtr-regulated inputs to this region that differ between males and females^3,4,58–61,67–69^. Consistent with this model, circuits that differ between the sexes influence the function of less dimorphic brain regions to generate sex-typical neural activity and behavior across species^70–72^. Our observations that loss of Oxtr has different, and even opposing, effects on prosocial behaviors and activity in the NAc between males and females may reflect that Oxtr signaling and other neuromodulatory pathways facilitating pair bonding evolved in prairie voles to act on ancestral, sexually dimorphic neural circuits^13,73,74^. Such neuromodulation may thus influence these pathways in prairie voles and other monogamous species to generate more monomorphic and synchronized reciprocal patterns of behavior to promote long-term attachment between mates^20,75–83^.

In the absence of Oxtr function, we find that a population of naïve males already contains two distinct classes. These populations are distinguished by whether a male shows a robust preference for his “partner” or a novel “stranger” following days of cohabitation with the partner. Notably, whether an Oxtr^1-/-^ male will ultimately prefer the partner female can be predicted by social behavior and neural activity during his first interactions with that female “partner.” The underlying biology of these two populations remains unknown; however, stranger-preferring males may constitute a genetically distinct population that is less responsive to Oxtr-independent pathways that influence males’ propensity for pair bond formation^84–86^. Alternatively, the distinct prosocial behavior displayed by stranger-preferring males may impact females’ reciprocal social behavior in such a way that males prefer not to continue interacting when given a choice^20^. The predictive relationship between initial social interactions and the fidelity of future bonds in the absence of Oxtr suggests that Oxtr signaling functions to reinforce attachment between mates or even to override other pathways controlling social behaviors in animals with a lower propensity for pair bond formation^81–83^. By decorrelating variation in behavior and neural activity during initial social interactions between partners from the social and neural mechanisms that mediate pair bonding between them, Oxtr may have evolved to reduce promiscuity amongst male prairie voles, decrease sexual dimorphism in behavior, and reinforce behaviors that facilitate enduring attachments between mates.

By observing a wide range of social interactions across pair bonding, we investigated how modulatory mechanisms that have evolved to facilitate pair bonding influence both the behavior and neural activity that underlie long-term social attachments^19,20,72^. The effects of Oxtr signaling vary across time and sex, supporting a model in which other neuromodulatory pathways intersect with Oxtr function during bonding and attachment to influence prosocial and agonistic behaviors in a state-dependent manner. Extending these circuit and behavioral investigations to other brain regions will enable us to determine how Oxtr and other signaling systems influence the activity of specific neural populations to control distinct modules of attachment, shedding light on the evolutionary processes driving social monogamy and the complex patterns of reciprocal interactions that constitute enduring relationships.

## Methods

### Animals

All animal care and procedures were approved by the University of California, San Francisco, Institutional Animal Care and Use Committee. A total of 76 adult (55-78 days at the start of behavior assays), sexually naive prairie voles (*Microtus ochrogaster*) were used in this study. Of these, 31 animals were rejected for experiments or analyses due to death, poor signal quality, head cap loss, ataxia, excessive aggression towards the partner, mistargeting of the fiber, or incorrect genotyping. We used both male and female voles, with sex determined by the presence or absence of testes at weaning. Voles were bred in our laboratory from a population that originated from systematic outbreeding of a wild-caught stock captured near Champagne, IL, and housed at our facilities at the University of California, San Francisco. Oxtr^1-/-^ voles were derived from a line that we previously generated^19^, and wild-type (WT) voles were obtained from the same outcrossing background line. Breeding voles were maintained in large, plastic cages (10½” W x 19” L x 8” H, Ancare, R20 Rat/Guinea Pig caging) on Paperchip bedding (Shepherd Specialty Papers). Weaned voles were maintained at our breeding facility in clear plastic cages (45 × 25 × 15 cm, Innovive, Innocage IVC Rat Caging) on Paperchip bedding until they were transferred to our lab housing facility, at which point they were transferred to Sani-Chips woodchip bedding (P.J. Murphy - Forest Products Corp.). Voles were weaned at 21-25 days into group-housed cages, with 2-6 total same-sex siblings or similarly aged weanlings in a cage. Group-housed cages were given 2 cotton nestlets and a large PVC elbow tube. Voles had *ad libitum* access to food and water. When animals were paired, voles were housed in 30.80 x 30.80 x 18.72 cm cages (Thoren, Maxi-Miser Model #4) on Sani-Chips bedding and provided with 2 cotton nestlets and 2 small plastic tubes. All animals were kept on a 14:10 light-dark cycle.

### Genotyping

At the time of weaning, we collected a small tail sample from each animal and digested the tissue in lysis buffer with proteinase K (Sigma-Aldrich, St. Louis, MO). We conducted a polymerase chain reaction for the Oxtr gene using the following primers: Forward ACTGGAGCTTCGAGTTGGAC; Reverse ATGCCCACCACTTGCAAGTA. The resulting product was digested using XcmI enzyme (New England Biolabs, Ipswich, MA) and run on a 2% agarose gel. A second tail sample was collected after an animal concluded all assays, and genotyping was repeated for confirmation. Animals whose post-experiment samples did not match the original genotyping were excluded from all analyses.

### Surgery

Voles aged P29-55 were anesthetized with isoflurane and administered bupivicaine subcutaneously at the incision site. Using a stereotaxic frame (Kopf Instruments, Tujunga, CA), a craniotomy was made +1.7 mm anterior and +1.2 mm lateral relative to bregma, and a 33-gauge cannula was lowered to -5.1 mm relative to bregma. We injected a 5:1 mixture of AAV8-Syn-GCaMP6m-WPRE (1×10^13^ GC/mL, Vigene Biosciences, Inc., Rockville, MD) and AAV8-CAG-tdTomato (5×10^12^ GC/mL, Dr. Ed Boyden Lab, UNC Vector Core). We infused 0.7-1 µL of virus at a rate of 0.1 µL/min via an automated injection system (Genie Touch, Kent Scientific Corporation, Torrington, CT). The cannula was left in place for ten minutes to allow for viral diffusion, then was slowly removed to minimize viral infection along the needle tract. Following viral injection, 2 microscrews were implanted into the skull to provide additional stability for the dental cement. A fiber optic cannula (400 μm-diameter silica core, 0.48 NA, 6.5 mm length with flat tip) with metal ferrule (1.25 mm base dimension, Doric Lenses, Inc., Québec, Canada) was implanted 0.05 mm above the injection site and secured to the skull with Metabond acrylic cement. Additionally, we created a well with which to hold a silicone elastomer that would provide extra stability for the fiber optic patch cable: A small hole was drilled into the cap of a microcentrifuge tube. The cap was placed upside down over the implant with the fiber ferrule protruding through the hole. The cap was secured to the Metabond with Ortho-Jet acrylic resin. Animals received postoperative buprenorphine and recovered for two to three weeks prior to testing. After three weeks, we checked each animal’s signal quality; animals with fluctuations in the 488 nm channel <5% of baseline (*i.e.*, the signal output while the animals were still) and/or insufficient signal output in the 581 nm channel (<10% of background fluorescence) were excluded from further study.

### Fiber Photometry

Photometry experiments were conducted with an RZ5P Base Processor and Synapse software (Tucker-Davis Technologies, Alachua, FL). The RZ5P modulated a two-channel LED driver (Doric Lenses) controlling two connectorized LEDs (465 nm and 560 nm, Doric Lenses). The LEDs were coupled to a two-color filter cube system (5 port Fluorescence Mini Cube, Doric Lenses) allowing transmission of 465-480 nm light and 555-570 nm light for excitation of GCaMP and tdTomato, respectively, and capture of 500-540 nm and 580-680 nm light for monitoring brain fluorescence. Light was transmitted to a 1×1 fiber-optic rotary joint (Doric Lenses), which coupled to a patch cable (400 μm-diameter silica core, 0.48 NA, with 1.1 mm hytrel protective jacket, Doric Lenses) connected to the animal’s fiber optic cannula with a bronze or zirconia mating sleeve. Emission was filtered by the mini cube and captured by two 2151 Femtowatt Photoreceivers (Newport Corporation, Irvine, CA). The RZ5P recorded raw broadband photoreceiver traces. A USB camera was mounted above the cage and recorded behavior at 20 frames per second. Both photometry and camera recordings were controlled by the Tucker-Davis Technologies Synapse software, with camera frames and photometry samplings time stamped for later synchronization.

### Behavior

Photometry procedures and habituation: All procedures were carried out during the light cycle. Introductions occurred between 08H00 and 12H00, and all other assays occurred between 12H00 and 20H00. To habituate animals to fiber tethering prior to testing, implanted voles were connected to a patch cable, placed in a clean cage, and given 30 minutes to habituate to fiber tethering at least 24 hours before social behavior tests began. On the day of an assay, voles were transferred from the housing room to the procedure room at least half an hour prior to testing. Voles were briefly restrained by the experimenter, and a patch cable was connected to the fiber optic implant. The cable was additionally secured to the cannula via a silicone elastomer (Ecoflex, Smooth-On, Macungie, PA). During all assays, all tubes, food hoppers, and water bottles were removed from the cage. An experimenter remained in the room away from the cage to monitor signal quality, fiber tangling or damage, and aggression. If tangling of the fiber occurred, or if voles chewed the fiber, the experimenter intervened to untangle the fiber or discourage the mischievous vole with a loud snap of the fingers. We terminated an assay a) if excessive chewing continued, b) if the fiber was irreparably damaged, or c) if 3 bouts of highly aggressive tussling occurred.

Introductions: After attaching the patch cable, the vole was placed in a clean cage and given at least 5 minutes to habituate to fiber placement. An age-matched, opposite-sex prairie vole was then introduced to the cage, and behavior was recorded for 30 minutes.

Mating: 24 hours after the introduction, a clear, plastic barrier with 1 cm-diameter holes was placed into the cage to separate the male and female. Each side was given one plastic tube and half of the nest. The following day, the implanted animal was connected to a patch cable and given at least 5 minutes to habituate to fiber placement. The barrier was then removed, and behavior was recorded for 30 minutes.

Partner preference test (PPT): A PPT was conducted 6 hours after pairing for females, or 3 days after mating for males. For females, animals were disconnected from the photometry system after the conclusion of the 30-minute introduction, and the cage was assembled and placed in the housing room until the 6 hours had elapsed. The PPT consists of a 3 chamber arena with an open top and 10 x 32in walls. Dividers extended 2in into the arena on either side to separate the apparatus into 3 equal-sized chambers. The partner and an age-matched, opposite-sex stranger are tethered on either end of the arena, and the subject is given free access to all chambers for 3 hours. To begin the assay, the implanted animal was connected to a patch cable, placed in the blocked-off center of the apparatus, and given at least 5 minutes to habituate to fiber tethering. The central barriers were then removed, and recordings lasted 3 hours. The placement of the partner and stranger relative to the orientation of the room were varied randomly by vole.

Separation and reunification: The implanted vole was connected to the patch cable and given at least 5 minutes to habituate to fiber placement. The partner vole was then removed from the home cage and placed in a separate, clean cage out of sight of the experimental vole. After one hour of separation, the partner was returned to the home cage, and behavior was recorded for 30 minutes.

Stranger rejection: The implanted vole was connected to a patch cable and given at least 5 minutes to habituate to fiber placement. The partner vole was then removed from the home cage and placed in a separate, clean cage out of sight of the experimental vole. After one hour of separation, a novel, age-matched, opposite-sex vole was placed in the home cage, and behavior was recorded for 20 minutes.

### Behavioral data analysis

Behavior scoring: Behavior was hand-scored frame by frame with the open-source software Boris^87^. Behaviors scored and their criteria are detailed in Table 1. The TDT photometry software recorded time stamps of each acquired behavior video frame as well as time stamps of neural data, allowing for synchronization of the two data streams. Scoring was conducted by independent observers blind to genotype and partner location.

For each animal, we quantified the number, total duration as percentage of the assay duration, and mean or median bout duration in seconds of each behavior and of social bouts. As described in Table 1, a social bout was defined as a sequence of behaviors initiated by non-AG sniffing, AG sniffing, or side-by-side contact, in which each behavior within the sequence was separated by less than 2 seconds of no interaction. For the PPT, chamber entries and exits were scored when the animal placed more than half of its body into an adjacent chamber. The start of an assay was defined as the moment when the stimulus animal was placed in the cage (having all 4 paws on the floor of the cage) or when the barriers dividing animals were fully removed from the cage (reaching the top edge of the cage or arena).

### Photometry Analysis

We illuminated each channel at sinusoidally-varying intensity using different modulation frequencies to improve signal to noise ratio with a lock-in amplifier system offline^1^. This system, coded in custom MATLAB software (version R2022b), involved performing a fast Fourier transform on the data from each channel to analyze the frequency domain between 100 and 500 Hz. The magnitudes (complex modulus) of the frequency-domain data streams were averaged across a band around the modulation frequency corresponding to each channel. We performed this calculation at each time point; this method identified deviations from the modulation frequency that reflect biological signal. We then low-pass IIR filtered this transformed data at 100 Hz to eliminate fluctuations due to technical noise.

The fluorescence data from both streams were trimmed to eliminate technical noise at the beginning of the recording. To estimate fluctuations in the GCaMP channel that are also present in the control channel and therefore due to noise across the system, we estimated a robust linear regression fitting the data from our control channel (tdTomato) to the GCaMP channel and found the value at each time point. We then calculated the difference between the GCaMP data and the fitted control fluorescence at each time point and normalized this difference by the fitted control fluorescence to result in our final ΔF/F value at each time point.

We extracted a photometry ΔF/F trace surrounding each behavioral timestamp of interest. For social bouts or individual behavior timestamps, we extracted photometry data from -2 to 5 seconds relative to each timestamp. Timestamps of behaviors occurring during periods of experimenter intervention were excluded from analyses. For each ΔF/F trace, we Z scored the trace to the mean and standard deviation of the ΔF/F values immediately prior to the timestamp (-2 to 0 seconds for social bouts). This method was chosen, in part, because we observed that NAc activity decreased below baseline during prolonged periods of quiet restfulness (Extended Data Fig. 9a-e). In addition, across all assays, we observed a notable increase in calcium signal that often preceded the introduction of the stimulus animal or removal of barriers, which may reflect a novelty, arousal, or fear response^88,89^ (Extended Data Fig. 9f-m). This signal decayed to pre-assay levels within roughly 100 seconds (Extended Data Fig. 9g). As a result, Z scores could potentially be biased depending on the behavior of the animal across the long recording period (*e.g.*, if the animal huddled with the stimulus animal for long periods of time) or during the pre-assay period (*e.g*. if the animal sat quietly vs. exploring the arena). For social touch behaviors, we excluded bouts in which social behavior (such as a strike) occurred within the 2s prior to the timestamp of interest. For strikes and chamber transitions, we excluded timestamps in which a strike or chamber transition occurred within the 2s baseline period.

To quantify calcium activity traces, we calculated peak ΔF/F and area under the curve (AUC) of each trace from 0 to 2 seconds after each timestamp. AUC was calculated using Matlab’s “trapz” function. To confirm that fluctuations and differences we observed in our ΔF/F traces were not due to motion artifacts, we used the same trace extraction and z scoring method on the tdTomato control fluorescence. This demonstrated there was little effect of motion at the onset of behaviors such as sniffs or even behaviors that entail a great deal of movement, such as attacks (Extended Data Fig. 9n-q).

### Perfusions, Histology, and Verification of Fiber Placement

Animals were deeply anesthetized with ketamine/xylazine and transcardially perfused with 1X phosphate-buffered saline (PBS) followed by cold 4% paraformaldehyde (PFA) in PBS. Brains were extracted, post-fixed in PFA overnight at 4°C, and incubated in 30% sucrose solution at 4°C until sucrose diffused completely through the brain. Brains were then embedded in Tissue-Tek Optimal Cutting Temperature Compound (Sakura Finetek USA, Torrance, CA), frozen solid at - 80°C, cryosectioned at 50 uM coronally, and treated with 300 nM DAPI solution (Life Technologies, Carlsbad, CA) in PBS for 10 minutes to visualize nuclei. Sections were mounted on glass slides and coverslipped with Aqua-Mount Mounting Medium (Thermo Scientific).

Brain sections were imaged with a 4X objective on a Nikon Eclipse 90i motorized upright epifluorescent microscope and digital camera (Nikon, Minato City, Tokyo, Japan) as well as a 10X objective on a Zeiss LSM 700 confocal microscope with Zen 2010 software (Zeiss Microscopy). Fiber tip locations were assessed by comparing anatomical location of the fiber tract to the Paxinos and Franklin mouse brain atlas (4th edition)^90^. Animals were excluded from behavioral and photometry analyses if the fiber tip was outside of the nucleus accumbens or outside the range of 1.10 - 1.78 mm relative to Bregma (of the mouse atlas).

### Statistics

Details of all statistical tests used and their results are reported in Extended Data File 1. Sample sizes were determined by our previous work investigating pair-bonding behaviors in Oxtr^1-/-^ prairie voles^20^ and increased by ∼20% to account for animals lost to surgery failure or photometry exclusion. Alpha was set to 0.05 for all comparisons. Trends were considered when p<0.07.

#### Behavior data

Statistics and plotting of behavioral data were performed in Prism (version 9.4 for MacOS). In each assay, the metric used to determine outlier status was total social interaction. Animals that were 3 scaled median absolute deviations (MAD) away from the median were deemed outliers and were removed before proceeding. Details on outliers are provided in Extended Data File 1. We assessed normality of residuals by the D’Agostino-Pearson omnibus (K2) test. For continuous and normally distributed measures, we conducted a Student’s t-test, ANOVA, two-way ANOVA, or two-way repeated measures ANOVA with Sidak-corrected post hoc comparisons. A Welch’s corrected t test was used when distributions failed an F test for equality of variances. For distributions that were not normally distributed, we first applied a log transform and retested for normality. Parametric tests were then run on the normalized data. For distributions that could not be log transformed, we used the non-parametric Mann Whitney test. We plotted the data in raw form when feasible. For count data, we used a generalized linear model with a generalized Poisson distribution and a Sidak correction for multiple comparisons to compare between groups. Transformations conducted on each data set prior to statistical testing can be found in Extended Data File 1. For proportion data, we used a binomial test to compare genotypes or groups. To characterize the relationship between behavior at 2 time points (*e.g.*, the introduction and the PPT), we used Pearson correlation or linear regression. To compare behavior across male preference categories (*i.e*., wild-type, partner preferring Oxtr^1-/-^, and stranger-preferring Oxtr^1-/-^), we used a permutation test on the F statistic. For the permutation test, group labels were randomly shuffled, and the statistic was calculated. This was repeated 10,000 times to construct a null distribution, and the p value was calculated as the percentage of the null distribution that was equal to or more extreme than the observed statistic. Post hoc pairwise comparisons were conducted with additional permutation tests with a Sidak correction for multiple comparisons.

For the PPT, preference indices were calculated for total percent of PPT time spent in either social interaction (“Social preference index”) or in side-by-side contact (“Side-by-side preference index”). Preference indices were calculated by taking the percent time spent with the partner, subtracting the time spent with the stranger, and normalizing by total time spent with either partner or stranger. Preference index values were compared across genotypes by performing a permutation test, as described above, on the Earth Mover’s Distance between the two distributions.

#### Markov chains

We performed discrete-time Markov chain analyses on behavioral sequences from the introduction assay. We included only “no interaction,” “non-AG sniff,” “AG sniff,” and “side-by-side contact” due to the fact that all other behavioral events were rare (transition probabilities less than 0.005). Markov chain transition probability matrices were calculated for each animal in Matlab, and probabilities for each transition were averaged across sex and genotype. For each transition, we compared transition probabilities across genotypes or groups by conducting permutation tests, as described above, on the Student’s t statistic or F statistic and applying a Sidak correction for multiple comparisons.

#### Adjusted boxplots

When plotting neural data surrounding behavioral events with duration (*e.g.,* AG sniff events), we included boxplots of those initiating behavior durations. Because duration distributions were heavily right skewed, we used an adjusted boxplot for skewed distributions^91^. Briefly, the medcouple (MC, a measure of skewness) is calculated for each distribution of duration values. The MC is then incorporated in the determination of the whisker boundaries, such that when MC >= 0, whisker boundaries are defined as [*Q*_1_ - 1.5e^-4^ ^MC^ IQR; *Q*_3_ + 1.5e^3^ ^MC^ IQR] and when MC < 0, boundaries are [*Q*_1_ - 1.5e^-3^ ^MC^ IQR; *Q*_3_ + 1.5e^4^ ^MC^ IQR], where *Q*_1_ is the first quartile, *Q*_3_ is the third quartile, and IQR is the interquartile range (*Q*_3_ - *Q*_1_). Outlier values are marked with a plus symbol. The adjusted boxplot was implemented with an adapted version of the function “adjusted_boxplot” written by Brian C. Coe (2024, MATLAB Central File Exchange, https://www.mathworks.com/matlabcentral/fileexchange/72110-adjusted_boxplot).

#### Probability PETHs

To examine the probability of the occurrence of different behaviors surrounding timestamps of interest, we constructed peri-event time histograms (PETHs) of probability (*e.g.,* Extended Data Figure 3c,d). For each timestamp and each time point along the PETH, a given behavior was marked as occurring (1) or not occurring (0). We then averaged by group across each time point.

#### Photometry data

All calculations and statistics for photometry data were performed in Matlab. For z-scored ΔF/F trace statistics (peak and AUC), we first removed outlier values. The criterion for removal was when both peak and AUC values for a given trace were 3 scaled MAD away from the median of the pooled data from all comparison groups. The majority of peak ΔF/F and AUC distributions were non-normally distributed as determined by the Jarque-Bera test; thus, for all comparisons, we pooled data from all groups and applied a Box-Cox transformation prior to testing. We then tested for main effects and interactions between independent variables (*e.g.*, genotype, stimulus animal, etc.) via a linear mixed effects (LME) model in which vole ID was included as a random effect to account for clustering within animals. The resulting model was run through an ANOVA, and we report the F statistics and p values. Multiple comparisons were conducted by pairwise LME models with a Sidak correction applied to the resulting p values. In addition, we compared z-scored ΔF/F values at each time point along the PETH. At each point, we Box-Cox transformed the pooled z-scored ΔF/F values. We then conducted an LME with vole ID included as a random effect, as described above. This was repeated for every time point and every pairwise comparison of, for example, genotype by stimulus animal. We then applied a Benjamini-Yekutieli correction on the resulting p values for *all* pairwise comparisons between groups.

To obtain mean neural responses by individual animal, we calculated the mean of the Box-Cox transformed values of the neural data. We used the transformed values to relate neural data to behavior or other neural metrics by individual animal via Pearson correlation or linear regression.

To test whether NAc calcium activity changed as an animal engaged in prolonged rest (Extended Data Figure 1a-e), we calculated median raw ΔF/F and AUC for each trace from -30s to 0s (pre) and 0s to 30s (post). We Box-Cox transformed all values and conducted an LME with time (pre vs. post) as the independent variable and both vole ID and trace number as random effects. We next calculated the change in ΔF/F (post – pre) for each trace and averaged by individual animal. We then used a two-tailed, one sample t test to determine whether the change in ΔF/F was significantly different from 0 ΔF/F. To test how activity changed at the start of an assay, we constructed PETHs for every animal and every assay from -10s to 180s surrounding assay start (Extended Data Figure 1f,g). We calculated the mean raw ΔF/F value from -10s to 0s to be 0.0237. At each time point along the PETH, we Box-Cox transformed all values including the constant of 0.0237 and conducted an LME with vole ID as a random effect to compare whether ΔF/F was significantly different from the transformed constant. We applied a Benjamini-Yekutieili correction to the resulting set of p values.

#### Plotting

Bar plots show mean +/- s.e.m. with individual animals overlaid. Neural data is plotted as mean +/- s.e.m. Swarm plots of behavior data show individual animals with median overlaid. Swarm plots of neural data show individual PETH values (peak or AUC) with median overlaid. Dotted lines on plots of linear regressions show the 95% confidence interval.

## Supporting information

Statistics

## Acknowledgements

We would like to thank Charles Frye for assistance with statistics. We also thank Kevin Bender and the members of the Manoli Lab for thoughtful comments on the manuscript.

## Author contributions

D.M., K.L., and N.H. devised experiments. K.L., N.H., R.K., D.S., J.W., and J.M. performed experiments. K.L., N.H., and A.K. performed analyses. D.M. directed the project. K.L., D.M., and N.H. wrote the manuscript.

## Data and materials availability

All data associated with this study are available upon request.

## Extended Data Materials

Extended Data Fig. 1: Fiber locations in the prairie vole nucleus accumbens.

Extended Data Fig. 2: Additional data from female introductions.

Extended Data Fig. 3: Cross-assay analyses in females.

Extended Data Fig. 4: Additional data from female timed mating assays.

Extended Data Fig. 5: Additional data from female partner preference tests.

Extended Data Fig. 6: Additional data from male introduction and timed mating assays.

Extended Data Fig. 7: Additional data from male PPT.

Extended Data Fig. 8: Additional data related to Figure 4.

Extended Data Fig. 9: Dynamics of NAc calcium activity during periods of rest and at assay start.

**Extended Data File 1:** Statistics

