## Supplementary material for "Oxytocin receptor function regulates neural signatures of pair bonding and fidelity in the nucleus accumbens": Statistics

### Table 1: Statistics

| Figure | Panel | Dependent Variable | Transformation applied | Test | Independent variable(s) | Random effect(s) | Test Statistic | p-value | Notes | Multiple Comparisons | | | | | | | |
|---|---|---|---|---|---|---|---|---|---|---|---|---|---|---|---|---|---|
| | | | | | | | | | | Comparison group 1 | | Comparison group 2 | | Unadjusted p value | Sidak corrected p | Median WT | Median OxtR-/- |
| 1 | f | % of time engaged in social bouts | None | T Test | Genotype | None | t(15) = 4.40 | 5.00E-04 | | | | | | | | 30.42% | 9.32% |
| | | Number of social bouts | None | GLM with a two-parameter generalized poisson distribution | Genotype | None | Z(15) = 3.441 | 5.80E-04 | | | | | | | | 97 | 56 |
| | | Median duration of social bouts | None | T Test | Genotype | None | t(15) = 2.00 | 6.31E-02 | | | | | | | | 2.25 | 1.45 |
| | g | % of social contact spent in non-AG sniff, AG sniff, or side-by-side contact | None | 2 way RM ANOVA | Genotype x Behavior | None | F (2, 30) = 1.631 | P=0.2126 | | | | | | | | | |
| | | | | | Genotype | | F (1, 15) = 1.812 | P=0.1983 | | | | | | | | | |
| | | | | | Behavior | | F (1.690, 25.35) = 4.277 | P=0.0306 | | | | | | | | | |
| | | | | | Subject | | F (15, 30) = 1.036e-018 | P>0.9999 | | | | | | | | | |
| | h | % of social bouts initiated by behavior | None | 2 way RM ANOVA | Genotype x Behavior | None | F (2, 30) = 0.6865 | P=0.5111 | | | | | | | | | |
| | | | | | Genotype | | F (1, 15) = 1.282 | P=0.2753 | | | | | | | | | |
| | | | | | Behavior | | F (1.275, 19.13) = 27.03 | P<0.0001 | | | | | | | | | |
| | | | | | Subject | | F (15, 30) = 3.402e-014 | P>0.9999 | | | | | | | | | |
| | i | Transition probability | None | Permutation test | Genotype | None | Non-AG sniff --> AG sniff: t(15) = 1.6304 | 1.12E-01 | | | | | | | 0.7589 | | |
| | | | | | Genotype | None | Non-AG sniff --> Side-by-side: t(15) = 1.1749 | 0.2643 | | | | | | | 0.9749 | | |
| | | | | | Genotype | None | Non-AG sniff --> No interaction: t(15) = -1.7038 | 0.1094 | Not shown in figure. See Fig. S3b | | | | | | 0.751 | | |
| | | | | | Genotype | None | AG sniff --> Non-AG sniff: t(15) = 0.3131 | 7.64E-01 | | | | | | | 1 | | |
| | | | | | Genotype | None | AG sniff --> Side-by-side: t(15) = 4.0370 | 1.70E-03 | | | | | | | 0.0202 | | |
| | | | | | Genotype | None | AG sniff --> No interaction: t(15) = -2.6878 | 2.04E-02 | Not shown in figure. See Fig. S3b | | | | | | 0.2191 | | |
| | | | | | Genotype | None | Side-by-side --> Non-AG sniff: t(15) = -1.3749 | 2.01E-01 | | | | | | | 0.9321 | | |
| | | | | | Genotype | None | Side-by-side --> AG sniff: t(15) = 2.0022 | 6.84E-02 | | | | | | | 0.5727 | | |
| | | | | | Genotype | None | Side-by-side --> No interaction: t(15) = 0.6808 | 5.53E-01 | Not shown in figure. See Fig. S3b | | | | | | 0.9999 | | |
| | | | | | Genotype | None | No interaction --> Non-AG sniff: t(15) = -1.1777 | 2.47E-01 | Not shown in figure. See Fig. S3b | | | | | | 0.9668 | | |
| | | | | | Genotype | None | No interaction --> AG sniff: t(15) = 0.6502 | 0.5259 | Not shown in figure. See Fig. S3b | | | | | | 0.999 | | |
| | | | | | Genotype | None | No interaction --> Side-by-side: t(15) = 1.4082 | 0.1809 | Not shown in figure. See Fig. S3b | | | | | | 0.9088 | | |
| | k | Peak Z scored dF/F | Box Cox | LME | Genotype | Vole | F(1,661) = 0.926 | 3.36E-01 | | | | | | | | 2.44 | 2.27 |
| | l | AUC of Z scored dF/F | Box Cox | LME | Genotype | Vole | F(1,661) = 1.755 | 1.86E-01 | | | | | | | | 80.66 | 53.02 |
| | n | Peak Z scored dF/F | Box Cox | LME | Genotype | Vole | F(1,427) = 12.964 | 3.55E-04 | | | | | | | | 2.76 | 1.83 |
| | o | AUC of Z scored dF/F | Box Cox | LME | Genotype | Vole | F(1,427) = 20.137 | 9.28E-06 | | | | | | | | 128.67 | -19.3 |

| Figure | Panel | Dependent Variable | Transformation applied | Test | Independent variable(s) | Random effect(s) | Test Statistic | p-value | Notes | Multiple Comparisons | | | | | | | | | |
|---|---|---|---|---|---|---|---|---|---|---|---|---|---|---|---|---|---|---|---|
| | | | | | | | | | | Comparison group 1 | | Comparison group 2 | | Unadjusted p value | Sidak corrected p | Median WT Partner | Median WT Stranger | Median OxtR-/- Partner | Median OxtR-/- Stranger |
| 2 | b | % of assay length spent in social bouts | None | 2 way RM ANOVA | Genotype x Stim | None | F (1, 16) = 0.1130 | P=0.7411 | | | | | | | | 67.57727466 | 4.192341535 | 60.04168692 | 0.598643056 |
| | | | | | Genotype | | F (1, 16) = 0.2171 | P=0.6476 | | | | | | | | | | | |
| | | | | | Stim | | F (1, 16) = 109.4 | P<0.0001 | | | | | | | | | | | |
| | | | | | Subject | | F (16, 16) = 0.9037 | P=0.5790 | | | | | | | | | | | |
| | c | Partner preference index | None | Permutation test on Earth-Movers Distance | Genotype | None | Observed EM Distance = 0.0791788831 | 0.5227 | 10000 iterations | | | | | | | WT: 0.93097401 | Oxtr-/-: 0.997973439 | | |
| | d | Difference in number of chamber entries between partner and stranger (partner - stranger) | Constant added (abs(min(difference))=5) | GLM with a two parameter Poisson distribution | Genotype | None | Z value (1,16) = -2.675 | 0.00747 | For multiple comparisons, we performed an Exact Poisson test for each genotype to test whether the difference between partner and stranger chamber entries was greater or less than 5 (the constant added). We then conducted GLMs with a two parameter Poisson distribution to test whether partner chamber entries or stranger chamber entries were different across genotypes. Resulting p values were Sidak corrected. | WT | Partner | WT | Stranger | 0.7513 | 0.9962 | 24.5 | 24.5 | 15 | 11 |
| | | | | | | | | | | MU | Partner | MU | Stranger | 1.80E-09 | 0.00001 | | | | |
| | | | | | | | | | | WT | Partner | MU | Partner | 0.0119 | 0.0468 | | | | |
| | | | | | | | | | | WT | Stranger | MU | Stranger | 2.19E-05 | 0.0001 | | | | |
| | e | Chamber preference index | None | Permutation test on Earth-Movers Distance | Genotype | None | Observed EM Distance = 0.143044993 | 0.0175 | 10000 iterations | | | | | | | WT: 0.000575209 | Oxtr-/-: 0.192028986 | | |
| | g | Peak Z scored dF/F | Box Cox | LME | Genotype | Vole | F(1,2015) = 4.8323 | 0.028044 | | WT | Partner | WT | Stranger | 0.30009 | 0.76002 | 2.45 | 2.32 | 2.52 | 1.92 |
| | | | | | Stim | | F(1,2015) = 13.952 | 0.00019276 | | WT | Partner | MU | Partner | 0.89749 | 0.99989 | | | | |
| | | | | | Genotype x Stim | | F(1,2015) = 5.5349 | 0.018736 | | WT | Stranger | MU | Stranger | 0.0051586 | 0.020475 | | | | |
| | | | | | | | | | | MU | Partner | MU | Stranger | 4.90E-05 | 0.00019593 | | | | |
| | h | AUC of Z scored dF/F | Box Cox | LME | Genotype | Vole | F(1,2015) = 1.9267 | 0.16527 | | WT | Partner | WT | Stranger | 0.22306 | 0.63562 | 82.4 | 60.43 | 117.95 | 9.85 |
| | | | | | Stim | | F(1,2015) = 22.994 | 1.74E-06 | | WT | Partner | MU | Partner | 0.27777 | 0.72792 | | | | |
| | | | | | Genotype x Stim | | F(1,2015) = 7.222 | 0.0072607 | | WT | Stranger | MU | Stranger | 0.012379 | 0.048602 | | | | |
| | | | | | | | | | | MU | Partner | MU | Stranger | 6.77E-07 | 2.71E-06 | | | | |
| | j | Peak Z scored dF/F | Box Cox | LME | Genotype | Vole | F(1,637) = 3.2608 | 0.071428 | | | | | | | | 2.76 | 2.72 | 2.69 | 2.36 |
| | | | | | Stim | | F(1,637) = 2.0377 | 0.15393 | | | | | | | | | | | |
| | | | | | Genotype x Stim | | F(1,637) = 0.094397 | 0.75876 | | | | | | | | | | | |
| | k | AUC of Z scored dF/F | Box Cox | LME | Genotype | Vole | F(1,637) = 1.7892 | 0.1815 | | | | | | | | 247.79 | 192.3 | 259.08 | 97.73 |
| | | | | | Stim | | F(1,637) = 12.126 | 0.00053139 | | | | | | | | | | | |
| | | | | | Genotype x Stim | | F(1,637) = 1.6044 | 0.20575 | | | | | | | | | | | |
| | m | Peak Z scored dF/F | Box Cox | LME | Genotype | Vole | F(1,649) = 0.1777 | 0.67349 | | WT | Partner | WT | Stranger | 3.12E-01 | 0.77536 | 2.84 | 3.07 | 2.43 | 3.67 |
| | | | | | Stim | | F(1,649) = 16.443 | 5.62E-05 | | WT | Partner | MU | Partner | 0.091745 | 0.31949 | | | | |
| | | | | | Genotype x Stim | | F(1,649) = 7.8655 | 0.0051894 | | WT | Stranger | MU | Stranger | 0.023089 | 0.089207 | | | | |

| | | | | | | | | | | | | | | | | | | | |
|---|---|---|---|---|---|---|---|---|---|---|---|---|---|---|---|---|---|---|---|
| | | | | | | | | | | MU | Partner | MU | Stranger | 2.95E-05 | 0.00011789 | | | | |
| | n | AUC of Z scored dF/F | Box Cox | LME | Genotype | Vole | F(1,649) = 5.7815 | 0.016475 | | WT | Partner | WT | Stranger | 0.1685 | 5.22E-01 | 200.2 | 279.53 | 197.67 | 330.87 |
| | | | | | Stim | | F(1,649) = 15.647 | 8.48E-05 | | WT | Partner | MU | Partner | 0.86796 | 0.9997 | | | | |
| | | | | | Genotype x Stim | | F(1,649) = 4.8049 | 0.028734 | | WT | Stranger | MU | Stranger | 0.0033573 | 0.013362 | | | | |

| Figure | Panel | Dependent Variable | Transformation applied | Test | Independent variable(s) | Random effect(s) | Test Statistic | p-value | Notes | Multiple Comparisons | | | | | | | | | | |
|---|---|---|---|---|---|---|---|---|---|---|---|---|---|---|---|---|---|---|---|---|
| | | | | | | | | | | Comparison group 1 | | Comparison group 2 | | Unadjusted p value | Sidak corrected p | Median category (if applicable) | Median WT | Median OxtR-/- | Median Ppref OxtR-/- | Median Spref OxtR-/- |
| 3 | b | % of assay engaged in social bouts | None | t test | Genotype | None | t(18) = 4.60 | 0.0002 | | | | | | | | | 13.00% | 28.60% | | |
| | | Number of social bouts | None | GLM with a two-parameter generalized poisson distribution | Genotype | None | Z(18) = -3.39 | 6.98E-04 | | | | | | | | | 71 | 97 | | |
| | | Median duration of social bouts | None | T Test | Genotype | None | t(18) = 3.09 | 0.0064 | | | | | | | | | 1.05 | 2.23 | | |
| | d | Peak Z scored dF/F | Box Cox | LME | Genotype | Vole | F(1,1613) = 0.01962 | 8.89E-01 | | | | | | | | | 2.34 | 2.35 | | |
| | e | AUC of Z scored dF/F | Box Cox | LME | Genotype | Vole | F(1,1613) = 0.24733 | 6.19E-01 | | | | | | | | | 84.97 | 92.77 | | |
| | g | % of assay engaged in social bouts | None | 2 way RM ANOVA | Genotype x Stim | None | F (1, 17) = 4.926 | P=0.0403 | | WT | Partner | MU | Partner | | 0.0361 | **Partner** | 56.31860454 | 40.02443801 | | |
| | | | | | Genotype | | F (1, 17) = 2.851 | P=0.1096 | | WT | Stranger | MU | Stranger | | 0.7334 | **Stranger** | 3.213392675 | 12.15005527 | | |
| | | | | | Stim | | F (1, 17) = 34.05 | P<0.0001 | | WT | Partner | WT | Stranger | | <0.0001 | | | | | |
| | | | | | Subject | | F (17, 17) = 0.3235 | P=0.9873 | | MU | Partner | MU | Stranger | | 0.0901 | | | | | |
| | h | Social preference index | None | Permutation test on Earth-Movers Distance | Genotype | None | Observed EM Distance = 0.63768867 | 0.0138 | 10000 iterations | | | | | | | | 0.91953704 | 0.606860969 | | |
| | | Side-by-side preference index | None | Permutation test on Earth-Movers Distance | Genotype | None | Observed EM Distance = 0.6753827 | 0.0191 | 10000 iterations | | | | | | | | 0.952052263 | 0.801206288 | | |
| | l | Peak Z scored dF/F | Box Cox | LME | Stim | Vole | F(1,3847) = 12.899 | 0.00033282 | | WT | Partner | WT | Stranger | 0.00017575 | 0.0015806 | **Partner** | 2.42 | | 2.78 | 2.18 |
| | | | | | Group | | F(2,3847) = 5.5641 | 0.0038639 | | WT | Partner | PPref | Partner | 0.0071621 | 0.062643 | **Stranger** | 1.96 | | 2.26 | 1.91 |
| | | | | | Group x Stim | | F(2,3847) = 3.2381 | 0.039345 | | WT | Partner | SPref | Partner | 0.055778 | 0.40342 | | | | | |
| | | | | | | | | | | WT | Stranger | PPref | Stranger | 0.77635 | 1 | | | | | |
| | | | | | | | | | | WT | Stranger | SPref | Stranger | 0.9467 | 1 | | | | | |
| | | | | | | | | | | PPref | Partner | PPref | Stranger | 4.66E-05 | 0.00041931 | | | | | |
| | | | | | | | | | | PPref | Partner | SPref | Partner | 0.0012831 | 0.011489 | | | | | |
| | | | | | | | | | | PPref | Stranger | SPref | Stranger | 0.4507 | 0.99545 | | | | | |
| | | | | | | | | | | SPref | Partner | SPref | Stranger | 0.96524 | 1 | | | | | |
| | m | AUC of Z scored dF/F | Box Cox | LME | Stim | Vole | F(1,3847) = 38.567 | 5.85E-10 | | WT | Partner | WT | Stranger | 6.48E-10 | 5.83E-09 | **Partner** | 124.61 | | 200.75 | 34.13 |
| | | | | | Group | | F(2,3847) = 6.9808 | 0.00094139 | | WT | Partner | PPref | Partner | 0.0007716 | 0.006923 | **Stranger** | 10.72 | | 42.73 | 22.11 |
| | | | | | Group x Stim | | F(2,3847) = 4.1826 | 0.015328 | | WT | Partner | SPref | Partner | 0.12949 | 0.71295 | | | | | |
| | | | | | | | | | | WT | Stranger | PPref | Stranger | 0.79998 | 1 | | | | | |
| | | | | | | | | | | WT | Stranger | SPref | Stranger | 0.94116 | 1 | | | | | |
| | | | | | | | | | | PPref | Partner | PPref | Stranger | 3.22E-09 | 2.90E-08 | | | | | |
| | | | | | | | | | | PPref | Partner | SPref | Partner | 0.00098771 | 0.0088544 | | | | | |
| | | | | | | | | | | PPref | Stranger | SPref | Stranger | 0.59451 | 0.9997 | | | | | |
| | | | | | | | | | | SPref | Partner | SPref | Stranger | 0.57209 | 0.99952 | | | | | |
| | n | Difference in z scored peak dF/F | None | Permutation test on F statistic | Group | None | Observed F = 3.2595 | 0.0644 | 100,000 iterations | WT | | PPref | | 6.97E-02 | 1.95E-01 | | | | | |
| | | | | | | | | | | WT | | SPref | | 4.72E-01 | 8.53E-01 | | | | | |
| | | | | | | | | | | PPref | | SPref | | 1.88E-02 | 5.53E-02 | | | | | |
| | | Difference in AUC | None | Permutation test on F statistic | Group | None | Observed F = 3.0894 | 0.0766 | 100,000 iterations | WT | | PPref | | 6.37E-02 | 1.79E-01 | | | | | |
| | | | | | | | | | | WT | | SPref | | 6.21E-01 | 9.46E-01 | | | | | |
| | | | | | | | | | | PPref | | SPref | | 1.62E-02 | 4.78E-02 | | | | | |
| | o | X: Difference in AUC; Y: % of assay engaged in social bouts with the partner | None | Pearson correlation | n/a | n/a | r = 0.1852 | 0.6085 | | | | | | | | | | | | |
| | | X: Difference in AUC; Y: % of assay engaged in social bouts with the partner | None | Pearson correlation | n/a | n/a | r = 0.9094 | 0.0007 | | | | | | | | | | | | |
| | p | X: Difference in AUC; Y: % of assay engaged in social bouts with the stranger | None | Pearson correlation | n/a | n/a | r = 0.1024 | 0.7783 | | | | | | | | | | | | |
| | | X: Difference in AUC; Y: % of assay engaged in social bouts with the stranger | None | Pearson correlation | n/a | n/a | r = -0.9615 | <0.0001 | | | | | | | | | | | | |

| Figure | Panel | Dependent Variable | Transformation applied | Test | Independent variable(s) | Random effect(s) | Test Statistic | p-value | Notes | Multiple Comparisons | | | | | | | | | |
|---|---|---|---|---|---|---|---|---|---|---|---|---|---|---|---|---|---|---|---|
| | | | | | | | | | | Comparison group 1 | | Comparison group 2 | | Unadjusted p value | Sidak corrected p | Median WT | Median Ppref OxtR-/- | Median Spref OxtR-/- | Median all OxtR-/- |
| | b | % time spent in social bouts during the intro | None | t test | Genotype | None | t(15) = 3.619 | 0.0028 | | | | | | | | 13.25 | 28.62 | 27.34 | 28.12 |
| | c | % of social contact spent in non-AG sniff, AG sniff, or side-by-side contact | None | 2 way RM ANOVA with Geisser-Greenhouse correction | Genotype x Behavior | Vole | F (4, 26) = 2.146 | P=0.1036 | | | | | | | | | | | |
| | | | | | Genotype | | F (2, 13) = 1.133 | P=0.3518 | | | | | | | | | | | |
| | | | | | Behavior | | F (1.167, 15.16) = 10.37 | P=0.0043 | | | | | | | | | | | |
| | | | | | Vole | | F (13, 26) = 2.054e-018 | P>0.9999 | | | | | | | | | | | |
| | d | Left = WT; Pearson correlations between intro behavior (% of social contact) and PPT side-by-side contact | None | Pearson correlation | Intro Non-AG sniff % vs. PPT Partner side-by-side | n/a | r = -0.211790912095149 | 0.556935236 | | | | | | | | | | | |
| | | | | | Intro AG sniff % vs. PPT Partner side-by-side | n/a | r = -0.259088860096925 | 0.469780022 | | | | | | | | | | | |
| | | | | | Intro side-by-side % vs. PPT Partner side-by-side | n/a | r = 0.268601335606747 | 0.453021976 | | | | | | | | | | | |
| | | | | | Intro Non-AG sniff % vs. PPT Stranger side-by-side | n/a | r = 0.445349117231156 | 0.197110273 | | | | | | | | | | | |
| | | | | | Intro AG sniff % vs. PPT Stranger side-by-side | n/a | r = 0.0527829986310332 | 0.884858338 | | | | | | | | | | | |
| | | | | | Intro side-by-side % vs. PPT Stranger side-by-side | n/a | r = -0.237610642018951 | 0.508591771 | | | | | | | | | | | |
| | | Right = OxtR-/-; Pearson correlations between intro behavior and PPT side-by-side contact | None | Pearson correlation | Intro Non-AG sniff % vs. PPT Partner side-by-side | n/a | r = 0.58237266892227 | 0.22519915 | | | | | | | | | | | |
| | | | | | Intro AG sniff % vs. PPT Partner side-by-side | n/a | r = 0.921368200517741 | 0.009031351 | | | | | | | | | | | |
| | | | | | Intro side-by-side % vs. PPT Partner side-by-side | n/a | r = -0.87985358183197 | 0.020785576 | | | | | | | | | | | |
| | | | | | Intro Non-AG sniff % vs. PPT Stranger side-by-side | n/a | r = -0.830552779829332 | 0.040635926 | | | | | | | | | | | |

| | | | | | | | | | | | | | | | | |
|---|---|---|---|---|---|---|---|---|---|---|---|---|---|---|---|---|
| | | | | | Intro AG sniff % vs. PPT Stranger side-by-side | n/a | r = -0.809429699980646 | 0.051015084 | | | | | | | | |
| | | | | | Intro side-by-side % vs. PPT Stranger side-by-side | n/a | r = 0.881864967809125 | 0.020109489 | | | | | | | | |
| | e | Left: X = % of intro engaged in AG sniffing; Y = % of PPT spent in side-by-side contact with partner | None | Linear regression | n/a | n/a | R squared = 0.8553; F(1,5) = 29.55 | 0.0029 | Y = 1.366*X - 3.925 | | | | | | | |
| | | Right: X = % of intro engaged in side-by-side contact; Y = % of PPT spent in side-by-side contact with partner | None | Linear regression | n/a | n/a | R squared = 0.7637; F(1,4) = 12.93 | 0.0228 | Y = -3.295*X + 72.30 | | | | | | | |
| | g | Peak Z scored dF/F | Box Cox | LME | Group | Vole | F(2,596) = 1.159 | 3.15E-01 | | | | | | 2.2 | 1.93 | 2.23 |
| | h | AUC of Z scored dF/F | Box Cox | LME | Group | Vole | F(2,596) = 1.3505 | 0.25991 | | | | | | 74.77 | 40.28 | 83.73 |
| | j | Peak Z scored dF/F | Box Cox | LME | Group | Vole | F(2,355) = 2.0272 | 1.85E-01 | | | | | | 2.62 | 2.13 | 2.74 |
| | k | AUC of Z scored dF/F | Box Cox | LME | Group | Vole | F(2,355) = 3.3965 | 0.034584 | | WT | Ppref | 0.080021 | 0.22137 | 111.88 | 27.15 | 143.86 |
| | | | | | | | | | | WT | Spref | 0.17781 | 0.4442 | | | |
| | | | | | | | | | | Ppref | Spref | 0.013367 | 0.039568 | | | |
| 4 | l | Left: X: Intro mean AG sniff Box Cox normalized AUC; Y: Mean % of PPT engaged in side-by-side contact with partner | None | Linear regression | n/a | n/a | R squared = 0.07353; F(1,8) = 0.6350 | 0.4485 | Y = -1.023*X + 123.1 | | | | | | | |
| | | Right: X: Intro mean AG sniff Box Cox normalized AUC; Y: Mean % of PPT engaged in side-by-side contact with stranger | None | Linear regression | n/a | n/a | R squared = 0.1570; F(1,8) = 1.49 | 0.257 | Y = 0.2144*X - 12.58 | | | | | | | |
| | m | Left: X: Intro mean AG sniff Box Cox normalized AUC; Y: Mean % of PPT engaged in side-by-side contact with partner | None | Linear regression | n/a | n/a | R squared =0.954; F(1,4) = 83.04 | 0.0008 | Y = -10.65*X + 717.9 | | | | | | | |
| | | Right: X: Intro mean AG sniff Box Cox normalized AUC; Y: Mean % of PPT engaged in side-by-side contact with stranger | None | Linear regression | n/a | n/a | R squared = 0.7157; F(1,4) = 10.07 | 0.0337 | Y = 3.238*X - 200.6 | | | | | | | |
| | n | Left: X: Intro mean AG sniff Box Cox normalized AUC; Y: Mean PPT Box Cox normalized partner social bout-elicited AUC | None | Linear regression | n/a | n/a | R squared = 0.3178; F(1,8) = 3.727 | 0.0896 | Y = 0.2171*X + 32.19 | | | | | | | |
| | | Right: X: Intro mean AG sniff Box Cox normalized AUC; Y: Mean PPT Box Cox normalized stranger social bout-elicited AUC | None | Linear regression | n/a | n/a | R squared = 0.03939; F(1,8) = 0.3281 | 0.5825 | Y = 0.09186*X + 39.61 | | | | | | | |
| | o | Left: X: Intro mean AG sniff Box Cox normalized AUC; Y: Mean PPT Box Cox normalized partner social bout-elicited AUC | None | Linear regression | n/a | n/a | R squared = 0.8765; F(1,4) = 28.39 | 0.006 | Y = -0.5231*X + 81.04 | | | | | | | |
| | | Right: X: Intro mean AG sniff Box Cox normalized AUC; Y: Mean PPT Box Cox normalized stranger social bout-elicited AUC | None | Linear regression | n/a | n/a | R squared = 0.09042; F(1,4) = 0.3976 | 0.5625 | Y = -0.08524*X + 51.24 | | | | | | | |
| | p | % time spent in side-by-side contact during the intro | None | Permutation test on F statistic | Group | None | Observed F = 8.6338 | 0.0062 | 10,000 iterations | WT | PPref | 0.4441 | 0.82821 | 6.455598912 | 8.889888889 | 22.01116667 |
| | | | | | | | | | | WT | SPref | 0.0081 | 0.024104 | | | |
| | | | | | | | | | | PPref | SPref | 0.11528 | 0.3075 | | | |
| | | Number of side-by-side instances during the intro | None | GLM with two parameter Poisson distribution | Group | None | F(2,13) = 6.9877 | 0.03038 | Multiple comparisons were performed with a simple GLM with a Poisson distribution. | WT | PPref | <0.0001 | <0.0001 | 36 | 78 | 90 |
| | | | | | | | | | | WT | SPref | <0.0001 | <0.0001 | | | |
| | | | | | | | | | | PPref | SPref | 0.125 | 0.3301 | | | |
| | s | Peak Z scored dF/F | Box Cox | LME | Group | Vole | F(2,274) = 4.3416 | 1.39E-02 | | WT | Ppref | 0.10616 | 0.28587 | 2.35 | 1.64 | 2.7 |
| | | | | | | | | | | WT | Spref | 0.1877 | 0.46403 | | | |
| | | | | | | | | | | Ppref | Spref | 0.0024588 | 0.0073582 | | | |
| | t | AUC of Z scored dF/F | Box Cox | LME | Group | Vole | F(2,274) = 3.1474 | 0.044521 | | WT | Ppref | 0.050643 | 0.14436 | 92.58 | -3.68 | 98.45 |
| | | | | | | | | | | WT | Spref | 0.60206 | 0.93698 | | | |
| | | | | | | | | | | Ppref | Spref | 0.017462 | 0.051477 | | | |
| | | | | | | | | | | Multiple Comparisons | | | | | | |
| **Figure** | **Panel** | **Dependent Variable** | **Transformation applied** | **Test** | **Independent variable(s)** | **Random effect(s)** | **Test Statistic** | **p-value** | **Notes** | **Comparison group 1** | **Comparison group 2** | **Unadjusted p value** | **Sidak corrected p** | **Median WT** | **Median OxtR-/-** | |
| S2 | a | % of assay time spent non-AG sniffing | Log | t test | Genotype | n/a | t(15) = 2.620 | 0.0193 | | | | | | 7.358 | 2.42 | |
| | | % of assay time spent AG sniffing | None | Welch's t test | Genotype | n/a | t(8.902) = 5.763 | 0.0003 | | | | | | 7.298 | 2.572 | |
| | | % of assay time spent in side-by-side contact | None | t test | Genotype | n/a | t(15) = 3.14 | 0.0067 | | | | | | 7.457 | 0.2463 | |
| | b | Transition probability | None | Permutation test | Genotype | None | Non-AG sniff --> AG sniff: t(15) = 1.6304 | 1.12E-01 | | | | | 0.7589 | | | |
| | | | | | Genotype | None | Non-AG sniff --> Side-by-side: t(15) = 1.1749 | 0.2643 | | | | | 0.9749 | | | |
| | | | | | Genotype | None | Non-AG sniff --> No interaction: t(15) = -1.7038 | 0.1094 | | | | | 0.751 | | | |
| | | | | | Genotype | None | AG sniff --> Non-AG sniff: t(15) = 0.3131 | 7.64E-01 | | | | | 1 | | | |
| | | | | | Genotype | None | AG sniff --> Side-by-side: t(15) = 4.0370 | 1.70E-03 | | | | | 0.0202 | | | |
| | | | | | Genotype | None | AG sniff --> No interaction: t(15) = -2.6878 | 2.04E-02 | | | | | 0.2191 | | | |
| | | | | | Genotype | None | Side-by-side --> Non-AG sniff: t(15) = -1.3749 | 2.01E-01 | | | | | 0.9321 | | | |
| | | | | | Genotype | None | Side-by-side --> AG sniff: t(15) = 2.0022 | 6.84E-02 | | | | | 0.5727 | | | |
| | | | | | Genotype | None | Side-by-side --> No interaction: t(15) = 0.6808 | 5.53E-01 | | | | | 0.9999 | | | |

| | | | | | | | | | | | | | | | | | | | |
|---|---|---|---|---|---|---|---|---|---|---|---|---|---|---|---|---|---|---|---|
| | | | | | Genotype | None | No interaction --> Non-AG sniff: t(15) = -1.1777 | 2.47E-01 | | | | | | | 0.9668 | | | | |
| | | | | | Genotype | None | No interaction --> AG sniff: t(15) = 0.6502 | 0.5259 | | | | | | | 0.999 | | | | |
| | | | | | Genotype | None | No interaction --> Side-by-side: t(15) = 1.4082 | 0.1809 | | | | | | | 0.9088 | | | | |
| | f | Peak z-scored dF/F | Box Cox | LME | Genotype | Vole | F(1,126) = 0.0002439 | 0.98756 | | | | | | | | 2.3358 | 2.1395 | | |
| | g | AUC of z-scored dF/F | Box Cox | LME | Genotype | Vole | F(1,126) = 0.012985 | 0.90946 | | | | | | | | 108.276 | 72.5894 | | |
| | i | Number of strikes | None | GLM with a two-parameter generalized poisson distribution | Genotype | None | Z(15) = -1.384 | 0.166 | | | | | | | | 12 | 17 | | |
| | k | Peak z-scored dF/F | Box Cox | LME | Genotype | Vole | F(1,372) = 2.3812 | 0.12365 | | | | | | | | 3.1877 | 3.6923 | | |
| | l | AUC of z-scored dF/F | Box Cox | LME | Genotype | Vole | F(1,372) = 3.7182 | 0.054583 | | | | | | | | 131.9738 | 226.951 | | |
| | | | | | | | | | | Multiple Comparisons | | | | | | | | | |
| **Figure** | **Panel** | **Dependent Variable** | **Transformation applied** | **Test** | **Independent variable(s)** | **Random effect(s)** | **Test Statistic** | **p-value** | **Notes** | **Comparison group 1** | | **Comparison group 2** | | **Unadjusted p value** | **Sidak corrected p** | **Median WT Partner** | **Median WT Stranger** | **Median OxtR-/- Partner** | **Median OxtR-/- Stranger** |
| S3 | b | % of assay spent non-AG sniffing | None | LME | Genotype | Vole | F (1, 19) = 0.9512 | 0.3417 | | WT | Intro | MU | Intro | | 0.0504 | **Intro** | 7.378300678 | 2.419833333 | |
| | | | | | Assay | | F (2, 32) = 1.971 | 0.1559 | | WT | Reunion | MU | Reunion | | >0.9999 | **Reunion** | 3.386217982 | 5.028498022 | |
| | | | | | Genotype x Assay | | F (2, 32) = 4.616 | 0.0173 | | WT | Stranger Exp | MU | Stranger Exp | | 0.9959 | **Stranger Exp** | 1.683583333 | 4.22925 | |
| | | | | | | | | | | WT | Intro | WT | Reunion | | 0.084 | | | | |
| | | | | | | | | | | WT | Intro | WT | Stranger Exp | | 0.0287 | | | | |
| | | | | | | | | | | WT | Reunion | WT | Stranger Exp | | >0.9999 | | | | |
| | | | | | | | | | | MU | Intro | MU | Reunion | | 0.9943 | | | | |
| | | | | | | | | | | MU | Intro | MU | Stranger Exp | | 0.9989 | | | | |
| | | | | | | | | | | MU | Reunion | MU | Stranger Exp | | >0.9999 | | | | |
| | c | % of assay spent AG sniffing | None | LME | Genotype | Vole | F (1, 19) = 5.832 | 0.026 | | WT | Intro | MU | Intro | | <0.0001 | **Intro** | 7.298388889 | 2.572 | |
| | | | | | Assay | | F (2, 32) = 13.01 | <0.0001 | | WT | Reunion | MU | Reunion | | 0.9987 | **Reunion** | 2.034891635 | 2.578503309 | |
| | | | | | Genotype x Assay | | F (2, 32) = 13.69 | <0.0001 | | WT | Stranger Exp | MU | Stranger Exp | | >0.9999 | **Stranger Exp** | 2.481916667 | 2.59075 | |
| | | | | | | | | | | WT | Intro | WT | Reunion | | <0.0001 | | | | |
| | | | | | | | | | | WT | Intro | WT | Stranger Exp | | <0.0001 | | | | |
| | | | | | | | | | | WT | Reunion | WT | Stranger Exp | | 0.9995 | | | | |
| | | | | | | | | | | MU | Intro | MU | Reunion | | >0.9999 | | | | |
| | | | | | | | | | | MU | Intro | MU | Stranger Exp | | >0.9999 | | | | |
| | | | | | | | | | | MU | Reunion | MU | Stranger Exp | | >0.9999 | | | | |
| | d | % of assay spent in side-by-side contact | None | LME | Genotype | Vole | F (1, 51) = 0.5178 | 0.475 | | Intro | | Reunion | | | <0.0001 | **Intro** | 7.456611111 | 0.246277778 | |
| | | | | | Assay | | F (2, 51) = 44.66 | <0.0001 | | Intro | | Stranger Exp | | | 0.998 | **Reunion** | 40.97403032 | 39.60516667 | |
| | | | | | Genotype x Assay | | F (2, 51) = 0.5973 | 0.5541 | | Reunion | | Stranger Exp | | | <0.0001 | **Stranger Exp** | 2.5245 | 2.83225 | |
| | f | Peak Z scored dF/F | Box Cox | LME | Genotype | Vole | F(1,1742) = 0.68597 | 0.40765 | | Intro | | SepReu | | 0.011051 | 0.032789 | **Intro** | 2.45 | 2.29 | |
| | | | | | Assay | | F(1,1742) = 4.1223 | 0.016366 | | Intro | | Stranger Exp | | 0.79275 | 0.9911 | **Reunion** | 2.73 | 2.93 | |
| | | | | | Genotype x Assay | | F(1,1742) = 1.8459 | 0.15819 | | SepReu | | Stranger Exp | | 0.01626 | 0.047991 | **Stranger Exp** | 2.23 | 2.26 | |
| | g | AUC of Z scored dF/F | Box Cox | LME | Genotype | Vole | F(1,1742) = 4.346 | 0.037242 | | WT | Intro | WT | SepReu | 0.0087728 | 0.076241 | **Intro** | 83 | 54.3 | |
| | | | | | Assay | | F(1,1742) = 12.876 | 2.81E-06 | | WT | Intro | WT | Stranger Exp | 0.2313 | 0.90628 | **Reunion** | 167.15 | 219.43 | |
| | | | | | Genotype x Assay | | F(1,1742) = 2.7813 | 0.062231 | | WT | Intro | MU | Intro | 0.2213 | 0.89473 | **Stranger Exp** | 66.73 | 70.12 | |
| | | | | | | | | | | WT | SepReu | WT | Stranger Exp | 0.36105 | 0.98225 | | | | |
| | | | | | | | | | | WT | SepReu | MU | SepReu | 0.64159 | 0.9999 | | | | |
| | | | | | | | | | | WT | Stranger Exp | MU | Stranger Exp | 0.01447 | 0.12294 | | | | |
| | | | | | | | | | | MU | Intro | MU | SepReu | 6.16E-05 | 0.0005547 | | | | |
| | | | | | | | | | | MU | Intro | MU | Stranger Exp | 0.4288 | 0.99353 | | | | |
| | | | | | | | | | | MU | SepReu | MU | Stranger Exp | 1.65E-05 | 0.00014881 | | | | |
| | i | Peak Z scored dF/F | Box Cox | LME | Genotype | Vole | F(1,711) = 1.5138 | 0.21896 | | WT | Intro | WT | SepReu | 0.75489 | 1 | **Intro** | 2.76 | 1.83 | |
| | | | | | Assay | | F(1,711) = 1.0429 | 0.35298 | | WT | Intro | WT | Stranger Exp | 0.05637 | 0.40678 | **Reunion** | 2.45 | 2.96 | |
| | | | | | Genotype x Assay | | F(1,711) = 3.1808 | 0.042142 | | WT | Intro | MU | Intro | 0.00039127 | 0.0035159 | **Stranger Exp** | 2.1 | 2.49 | |
| | | | | | | | | | | WT | SepReu | WT | Stranger Exp | 0.048592 | 0.3613 | | | | |
| | | | | | | | | | | WT | SepReu | MU | SepReu | 0.24312 | 0.91848 | | | | |
| | | | | | | | | | | WT | Stranger Exp | MU | Stranger Exp | 0.29491 | 0.95693 | | | | |
| | | | | | | | | | | MU | Intro | MU | SepReu | 0.11943 | 0.68167 | | | | |
| | | | | | | | | | | MU | Intro | MU | Stranger Exp | 0.09852 | 0.60681 | | | | |
| | | | | | | | | | | MU | SepReu | MU | Stranger Exp | 0.72437 | 0.99999 | | | | |
| | j | AUC of Z scored dF/F | Box Cox | LME | Genotype | Vole | F(1,711) = 3.8669 | 0.049636 | | WT | Intro | WT | SepReu | 0.14184 | 0.74758 | **Intro** | 128.67 | -19.3 | |
| | | | | | Assay | | F(1,711) = 4.8172 | 0.0083554 | | WT | Intro | WT | Stranger Exp | 0.071162 | 0.48541 | **Reunion** | 166.4 | 129.48 | |
| | | | | | Genotype x Assay | | F(1,711) = 3.7309 | 0.024442 | | WT | Intro | MU | Intro | 7.36E-06 | 6.63E-05 | **Stranger Exp** | 56.33 | 64.36 | |
| | | | | | | | | | | WT | SepReu | WT | Stranger Exp | 0.0092628 | 0.080342 | | | | |
| | | | | | | | | | | WT | SepReu | MU | SepReu | 0.11103 | 0.65327 | | | | |

| | | | | | | | | | | | | | | | | | | | |
|---|---|---|---|---|---|---|---|---|---|---|---|---|---|---|---|---|---|---|---|
| | [illegible] | AUC of z scored dF/F | Box Cox | LME | | | | | | WT | Stranger Exp | MU | Stranger Exp | 0.42384 | 0.993 | | | | |
| | | | | | | | | | | MU | Intro | MU | SepReu | 0.0052736 | 0.046474 | | | | |
| | | | | | | | | | | MU | Intro | MU | Stranger Exp | 0.041971 | 0.32016 | | | | |
| | | | | | | | | | | MU | SepReu | MU | Stranger Exp | 0.89807 | 1 | | | | |
| | l | Peak z scored dF/F | Box Cox | LME | Genotype | Vole | F(1,486) = 2.941 | 0.086996 | | | | | | | | **Intro** | 2.3358 | 2.1395 | |
| | | | | | Assay | | F(2,486) = 1.4308 | 0.24012 | | | | | | | | **Reunion** | 2.4484 | 2.7646 | |
| | | | | | Genotype*Assay | | F(2,486) = 1.1706 | 0.31104 | | | | | | | | **Stranger Exp** | 2.0028 | 6.1549 | |
| | m | AUC of z scored dF/F | Box Cox | LME | Genotype | Vole | F(1,486) = 4.7146 | 0.030391 | | | | | | | | **Intro** | 108.276 | 72.5894 | |
| | | | | | Assay | | F(2,486) = 0.95724 | 0.38467 | | | | | | | | **Reunion** | 111.4367 | 142.5169 | |
| | | | | | Genotype*Assay | | F(2,486) = 2.1791 | 0.11425 | | | | | | | | **Stranger Exp** | 9.7616 | 493.5979 | |
| | n | Number of strikes | None | Generalized linear mixed model fit by maximum likelihood (Laplace Approximation) ['glmerMod'] | Genotype | Vole | F(1,35) = 0.975 | 0.33 | | OxtR-/- | Stranger Exp | WT | Stranger Exp | 0.9973 | 1 | | | | |
| | | | | | Assay | | F(1,35) = -4.571 | 4.85E-06 | | OxtR-/- | Stranger Exp | OxtR-/- | Intro | 0.00001 | 0.0001 | | | | |
| | | | | | Genotype*Assay | | F(1,33) = -4.215 | 2.50E-05 | | OxtR-/- | Stranger Exp | WT | Intro | 0.9992 | 1 | | | | |
| | | | | | | | | | | WT | Stranger Exp | OxtR-/- | Intro | 0.2762 | 0.8562 | | | | |
| | | | | | | | | | | WT | Stranger Exp | WT | Intro | 0.9963 | 1 | | | | |
| | | | | | | | | | | OxtR-/- | Intro | WT | Intro | 0.3133 | 0.8951 | | | | |
| | f | Peak z scored dF/F | Box Cox | LME | Genotype | Vole | F(1,701) = 5.6959 | 0.017268 | | | | | | | | **Intro** | 3.1877 | 3.6907 | |
| | | | | | Assay | | F(1,701) = 0.86864 | 0.35165 | | | | | | | | **Stranger Exp** | 3.3316 | 3.7474 | |
| | | | | | Genotype*Assay | | F(1,701) = 0.0045443 | 0.94627 | | | | | | | | | | | |
| | g | AUC of z scored dF/F | Box Cox | LME | Genotype | Vole | F(1,701) = 9.7436 | 0.0018734 | | | | | | | | **Intro** | 131.9738 | 225.396 | |
| | | | | | Assay | | F(1,701) = 2.5476 | 0.11091 | | | | | | | | **Stranger Exp** | 211.7916 | 261.4536 | |
| | | | | | Genotype*Assay | | F(1,701) = 0.083782 | 0.77232 | | | | | | | | | | | |
| **Figure** | **Panel** | **Dependent Variable** | **Transformation applied** | **Test** | **Independent variable(s)** | **Random effect(s)** | **Test Statistic** | **p-value** | **Notes** | **Multiple Comparisons** | | | | | | | | | |
| | | | | | | | | | | **Comparison group 1** | | **Comparison group 2** | | **Unadjusted p value** | **Sidak corrected p** | **Median WT** | **Median OxtR-/-** | | |
| **S4** | b | Proportion of females displaying mating behavior | None | Binomial test | Genotype | n/a | n/a | 0.1195 | | | | | | | | | | | |
| | c | % of assay time spent in mating behavior | None | t test | Genotype | n/a | t(12) = 0.1785 | 0.8613 | | | | | | | | 7.663 | 9.056 | | |
| | | Number of mating bouts | None | GLM with a two-parameter generalized poisson distribution | Genotype | n/a | Z(12) = 0.663 | 0.508 | | | | | | | | 36 | 36 | | |
| | d | % of time engaged in social bouts | None | T Test | Genotype | None | t(16) = 0.7287 | 4.77E-01 | | | | | | | | 40.73 | 32.4 | | |
| | | Number of social bouts | None | GLM with a two-parameter generalized poisson distribution | Genotype | None | Z(16) = 1.146 | 2.52E-01 | | | | | | | | 89 | 80.5 | | |
| | | Median duration of social bouts | None | Welch's t test | Genotype | None | t(13.06) = 1.21 | 2.48E-01 | | | | | | | | 2.175 | 2.663 | | |
| | e | % of assay time spent non-AG sniffing | None | t test | Genotype | n/a | t(16) = 0.3501 | 0.7309 | | | | | | | | 3.585 | 2.759 | | |
| | | % of assay time spent AG sniffing | None | t test | Genotype | n/a | t(16) = 0.3135 | 0.758 | | | | | | | | 2.577 | 1.545 | | |
| | | % of assay time spent in side-by-side contact | None | t test | Genotype | n/a | t(16) = 0.485 | 0.6342 | | | | | | | | 30.21 | 21.78 | | |
| | g | Peak z-scored dF/F | Box Cox | LME | Genotype | Vole | F(1,40) = 0.18806 | 0.66687 | | | | | | | | 1.024 | 1.7869 | | |
| | h | AUC of z-scored dF/F | Box Cox | LME | Genotype | Vole | F(1,40) = 2.0627 | 0.15872 | | | | | | | | -273.8318 | -129.5171 | | |
| | j | Peak z-scored dF/F | Box Cox | LME | Genotype | Vole | F(1,475) = 0.57902 | 0.44707 | | | | | | | | 1.378 | 1.1845 | | |
| | k | AUC of z-scored dF/F | Box Cox | LME | Genotype | Vole | F(1,475) = 0.76687 | 0.38163 | | | | | | | | -103.6138 | -119.1052 | | |
| | m | Peak z-scored dF/F | Box Cox | LME | Genotype | Vole | F(1,1282) = 0.053793 | 0.81663 | | | | | | | | 2.2066 | 2.2585 | | |
| | n | AUC of z-scored dF/F | Box Cox | LME | Genotype | Vole | F(1,1282) = 0.11397 | 0.73572 | | | | | | | | 79.1489 | 58.361 | | |
| | p | Peak z-scored dF/F | Box Cox | LME | Genotype | Vole | F(1,654) = 3.0119 | 0.083125 | | | | | | | | 2.2435 | 2.0774 | | |
| | q | AUC of z-scored dF/F | Box Cox | LME | Genotype | Vole | F(1,654) = 2.9562 | 0.086021 | | | | | | | | 105.632 | 49.5363 | | |
| | s | Peak z-scored dF/F | Box Cox | LME | Genotype | Vole | F(1,219) = 0.47972 | 0.48928 | | | | | | | | 2.4015 | 2.2628 | | |
| | t | AUC of z-scored dF/F | Box Cox | LME | Genotype | Vole | F(1,219) = 0.64224 | 0.42377 | | | | | | | | 110.1607 | 91.756 | | |
| | v | Peak z-scored dF/F | Box Cox | LME | Genotype | Vole | F(1,402) = 4.2097 | 0.04084 | | | | | | | | 1.9605 | 2.5162 | | |
| | w | AUC of z-scored dF/F | Box Cox | LME | Genotype | Vole | F(1,402) = 5.2625 | 0.022304 | | | | | | | | 52.3248 | 64.2403 | | |
| **Figure** | **Panel** | **Dependent Variable** | **Transformation applied** | **Test** | **Independent variable(s)** | **Random effect(s)** | **Test Statistic** | **p-value** | **Notes** | **Multiple Comparisons** | | | | | | | | | |
| | | | | | | | | | | **Comparison group 1** | | **Comparison group 2** | | **Unadjusted p value** | **Sidak corrected p** | **Median WT** | **Median OxtR-/-** | | |
| **S5** | a | % of assay length spent non-anogenital sniffing | None | 2 way RM ANOVA | Genotype x Stim | None | F (1, 16) = 0.06982 | P=0.7950 | | | | | | | | 0.948800926 | 0.700888889 | 0.745881525 | 0.248478299 |
| | | | | | Genotype | | F (1, 16) = 3.312 | P=0.0875 | | | | | | | | | | | |
| | | | | | Stim | | F (1, 16) = 1.608 | P=0.2230 | | | | | | | | | | | |
| | | | | | Subject | | F (16, 16) = 0.3348 | P=0.9824 | | | | | | | | | | | |
| | | % of assay length spent anogenital sniffing | None | 2 way RM ANOVA | Genotype x Stim | None | F (1, 16) = 1.367 | P=0.2594 | | | | | | | | 0.287426203 | 0.396703266 | 0.430100372 | 0.076668642 |
| | | | | | Genotype | | F (1, 16) = 0.1607 | P=0.6938 | | | | | | | | | | | |
| | | | | | Stim | | F (1, 16) = 0.4568 | P=0.5088 | | | | | | | | | | | |
| | | | | | Subject | | F (16, 16) = 0.6528 | P=0.7986 | | | | | | | | | | | |
| | | % of assay length spent in side-by-side contact | None | 2 way RM ANOVA | Genotype x Stim | None | F (1, 16) = 0.09258 | P=0.7648 | | | | | | | | 65.0233709 | 2.154766883 | 58.858926 | 0.067810185 |
| | | | | | Genotype | | F (1, 16) = 0.2397 | P=0.6310 | | | | | | | | | | | |
| | | | | | Stim | | F (1, 16) = 115.1 | P<0.0001 | | | | | | | | | | | |
| | | | | | Subject | | F (16, 16) = 0.9459 | P=0.5436 | | | | | | | | | | | |
| | b | Number of attacks directed at partner | None | GLM with a two-parameter generalized poisson distribution | Genotype | None | Z(16) = -1.333 | 0.18252 | | | | | | | | | 2 | | 2 |
| | | Number of attacks directed at stranger | None | GLM with a two-parameter generalized poisson distribution | Genotype | None | Z(16) = -0.032 | 0.974527 | | | | | | | | | | | |
| | | Difference between partner and stranger directed attacks | None | GLM with a two-parameter generalized poisson distribution | Genotype | None | Z(16) = 1.347 | 0.1779 | | | | | | | | | | | |

| | | | | | | | | | | | | | | | | | | | |
|---|---|---|---|---|---|---|---|---|---|---|---|---|---|---|---|---|---|---|---|
| | c | Porportions of females that showed any side-by-side contact events with the stranger | None | Binomial Test | Genotype | None | n/a | <0.0001 | | | | | | | | | | | |
| | f | Peak Z scored dF/F | Box Cox | LME | Genotype | Vole | F(1,1134) = 6.9989 | 0.0082689 | | | | | | | | 2.6297 | 2.1976 | 2.6557 | 1.803 |
| | | | | | Stim | | F(1,1134) = 16.422 | 5.41E-05 | | | | | | | | | | | |
| | | | | | Genotype x Stim | | F(1,1134) = 1.383 | 0.23984 | | | | | | | | | | | |
| | g | AUC of Z scored dF/F | Box Cox | LME | Genotype | Vole | F(1,1134) = 0.64629 | 0.42161 | | | | | | | | 118.2063 | 41.4769 | 121.5318 | 11.3818 |
| | | | | | Stim | | F(1,1134) = 14.685 | 1.34E-04 | | | | | | | | | | | |
| | | | | | Genotype x Stim | | F(1,1134) = 1.5517 | 0.21314 | | | | | | | | | | | |
| | i | Peak Z scored dF/F | Box Cox | LME | Genotype | Vole | F(1,454) = 1.7127 | 0.1913 | | | | | | | | 2.4843 | 2.4451 | 2.3103 | 2.035 |
| | | | | | Stim | | F(1,454) = 0.43978 | 5.08E-01 | | | | | | | | | | | |
| | | | | | Genotype x Stim | | F(1,454) = 0.19153 | 0.66185 | | | | | | | | | | | |
| | j | AUC of Z scored dF/F | Box Cox | LME | Genotype | Vole | F(1,454) = 0.28644 | 0.59277 | | | | | | | | 114.1553 | 82.3693 | 104.9737 | 11.0248 |
| | | | | | Stim | | F(1,454) = 1.3013 | 2.55E-01 | | | | | | | | | | | |
| | | | | | Genotype x Stim | | F(1,454) = 0.14474 | 0.70379 | | | | | | | | | | | |
| | | | | | | | | | | Multiple Comparisons | | | | | | | | | |
| **Figure** | **Panel** | **Dependent Variable** | **Transformation applied** | **Test** | **Independent variable(s)** | **Random effect(s)** | **Test Statistic** | **p-value** | **Notes** | **Comparison group 1** | | **Comparison group 2** | | **Unadjusted p value** | **Sidak corrected p** | **Median WT** | **Median OxtR-/-** | | |
| S6 | h | % of assay time spent non-AG sniffing | None | t test | Genotype | n/a | t(18) = 2.848 | 0.0107 | | | | | | | | 3.204 | 5.178 | | |
| | | % of assay time spent AG sniffing | Log(y) | t test | Genotype | n/a | t(18) = 1.857 | 0.0798 | | | | | | | | 1.901 | 4.488 | | |
| | | % of assay time spent in side-by-side contact | None | Mann Whitney test | Genotype | n/a | U = 11 | 0.0023 | | | | | | | | 6.451 | 13.7 | | |
| | f | % of social contact by behavior | None | 2 way RM ANOVA | Genotype x Behavior | None | F (2, 36) = 1.824 | P=0.1759 | | | | | | | | | | | |
| | | | | | Genotype | | F (1, 18) = 1.088 | P=0.3107 | | | | | | | | | | | |
| | | | | | Behavior | | F (1.143, 20.57) = 14.83 | P=0.0007 | | | | | | | | | | | |
| | | | | | Subject | | F (18, 36) = 3.247e-018 | P>0.9999 | | | | | | | | | | | |
| | g | % of social bouts initiated by behavior | None | 2 way RM ANOVA | Genotype x Behavior | None | F (2, 36) = 0.1504 | P=0.8609 | | | | | | | | | | | |
| | | | | | Genotype | | F (1, 18) = 0.5916 | P=0.4518 | | | | | | | | | | | |
| | | | | | Behavior | | F (1.654, 29.77) = 18.23 | P<0.0001 | | | | | | | | | | | |
| | | | | | Subject | | F (18, 36) = 4.503e-014 | P>0.9999 | | | | | | | | | | | |
| | e | % of time engaged in social bouts | None | 2 way ANOVA | Sex x Genotype | n/a | F (1, 33) = 40.42 | P<0.0001 | | Female | WT | Female | OxtR-/- | | 0.0005 | | | | |
| | | | | | Sex | | F (1, 33) = 0.9379 | P=0.3399 | | Female | WT | Male | WT | | 0.0032 | | | | |
| | | | | | Genotype | | F (1, 33) = 0.04213 | P=0.8386 | | Female | WT | Male | OxtR-/- | | 0.9963 | | | | |
| | | | | | | | | | | Female | OxtR-/- | Male | WT | | 0.9503 | | | | |
| | | | | | | | | | | Female | OxtR-/- | Male | OxtR-/- | | <0.0001 | | | | |
| | | | | | | | | | | Male | WT | Male | OxtR-/- | | 0.0004 | | | | |
| | a | Proportion of males displaying any strikes | None | Binomial test with Wilson/Brown method of calculating confidence intervals | Genotype | n/a | WT proportions: 45.45% no Stranger Exp, 54.55% Stranger Exp | 0.0008 | | | | | | | | | | | |
| | | | | | | | OxtR-/- proportions: 100% no Stranger Exp, 0% Stranger Exp | | | | | | | | | | | | |
| | c | Peak z scored dF/F | Box Cox | LME to test WT difference from 0 dF/F | n/a | Vole | F(1,11) = 77.252 | 2.64E-06 | | | | | | | | 3.7159 | | | |
| | d | AUC of z scored dF/F | Box Cox | LME to test WT difference from 0 AUC | n/a | Vole | F(1,11) = 53.339 | 1.54E-05 | | | | | | | | 450.2758 | | | |
| | j | Peak z scored dF/F | Box Cox | LME | Genotype | Vole | F(1,791) = 0.0008914 | 0.97619 | | | | | | | | 2.208 | 2.2811 | | |
| | k | AUC of z scored dF/F | Box Cox | LME | Genotype | Vole | F(1,791) = 0.024107 | 0.87665 | | | | | | | | 74.769 | 91.2702 | | |
| | m | Peak z scored dF/F | Box Cox | LME | Genotype | Vole | F(1,472) = 0.095612 | 0.7573 | | | | | | | | 2.5466 | 2.4116 | | |
| | n | AUC of z scored dF/F | Box Cox | LME | Genotype | Vole | F(1,472) = 0.66784 | 0.41422 | | | | | | | | 101.4363 | 95.7307 | | |
| | p | Peak z scored dF/F | Box Cox | LME | Genotype | Vole | F(1,340) = 0.33023 | 0.5659 | | | | | | | | 2.2947 | 2.3228 | | |
| | q | AUC of z scored dF/F | Box Cox | LME | Genotype | Vole | F(1,340) = 0.4535 | 0.50114 | | | | | | | | 79.1638 | 98.1405 | | |
| | r | % of time engaged in social bouts | None | T Test | Genotype | None | t(21) = 1.331 | 1.98E-01 | | | | | | | | 44.34 | 54.11 | | |
| | | Number of social bouts | None | GLM with a two-parameter generalized poisson distribution | Genotype | None | Z(21) = 0.623 | 5.33E-01 | | | | | | | | 66.5 | 64 | | |
| | | Median duration of social bouts | Log(y) | T test | Genotype | None | t(21) = 1.703 | 1.03E-01 | | | | | | | | 2.775 | 4.375 | | |
| | t | Peak z scored dF/F | Box Cox | LME | Genotype | Vole | F(1,1420) = 0.86174 | 0.35341 | | | | | | | | 2.5523 | 2.6 | | |
| | u | AUC of z scored dF/F | Box Cox | LME | Genotype | Vole | F(1,1420) = 0.90524 | 0.34154 | | | | | | | | 123.2521 | 143.2993 | | |
| | v | % of assay time spent non-AG sniffing | None | t test | Genotype | n/a | t(21) = 0.7514 | 0.4608 | | | | | | | | 3.619 | 2.797 | | |
| | | % of assay time spent AG sniffing | None | Mann Whitney test | Genotype | n/a | U = 54 | 0.4865 | | | | | | | | 4.428 | 7.693 | | |
| | | % of assay time spent in side-by-side contact | None | t test | Genotype | n/a | t(21) = 1.174 | 0.2536 | | | | | | | | 25.84 | 39.35 | | |
| | w | Proportion of males displaying mating behavior | None | Binomial test with Wilson/Brown method of calculating confidence intervals | Genotype | n/a | WT proportions: 41.67% no mating, 58.33% mating | >0.9999 | | | | | | | | | | | |
| | x | % of assay time spent in mating behavior | None | t test | Genotype | n/a | t(11) = 1.013 | 0.3329 | | | | | | | | 4.232 | 0.3907 | | |
| | | Number of mating bouts | None | GLM with a two-parameter generalized poisson distribution | Genotype | None | Z(11) = 1.039 | 0.299 | | | | | | | | 9 | 7.5 | | |
| | z | Peak z scored dF/F | Box Cox | LME | Genotype | Vole | F(1,34) = 2.6967 | 0.10978 | | | | | | | | 1.83 | 0.7269 | | |
| | aa | AUC of z scored dF/F | Box Cox | LME | Genotype | Vole | F(1,34) = 10.088 | 0.0031688 | | | | | | | | -93.8763 | -502.668 | | |
| | cc | Peak z scored dF/F | Box Cox | LME | Genotype | Vole | F(1,197) = 5.3218 | 0.0221 | | | | | | | | 1.3983 | 2.0371 | | |
| | dd | AUC of z scored dF/F | Box Cox | LME | Genotype | Vole | F(1,197) = 2.9359 | 0.088201 | | | | | | | | -92.0982 | -14.2155 | | |
| | | | | | | | | | | Multiple Comparisons | | | | | | | | | |
| **Figure** | **Panel** | **Dependent Variable** | **Transformation applied** | **Test** | **Independent variable(s)** | **Random effect(s)** | **Test Statistic** | **p-value** | **Notes** | **Comparison group 1** | | **Comparison group 2** | | **Unadjusted p value** | **Sidak corrected p** | **Medians** | **Partner** | **Stranger** | |
| | a | % of assay time spent non-AG sniffing | None | 2 Way RM ANOVA | Genotype*Stim | Subject | F (1, 17) = 6.463 | P=0.0210 | | WT | Partner | OxtR-/- | Partner | | 0.7566 | **WT** | 0.8325 | 0.6733 | |
| | | | | | Genotype | | F (1, 17) = 1.761 | P=0.2020 | | WT | Stranger | OxtR-/- | Stranger | | 0.0337 | **OxtR-/-** | 0.4419 | 1.0198 | |
| | | | | | Stim | | F (1, 17) = 4.275 | P=0.0542 | | WT | Partner | WT | Stranger | | 0.995 | | | | |
| | | | | | Subject | | F (17, 17) = 0.7445 | P=0.7252 | | OxtR-/- | Partner | OxtR-/- | Stranger | | 0.0219 | | | | |
| | | % of assay time spent AG sniffing | None | 2 Way RM ANOVA | Genotype*Stim | Subject | F (1, 17) = 0.5566 | P=0.4658 | | | | | | | | **WT** | 0.3039 | 0.7269 | |
| | | | | | Genotype | | F (1, 17) = 0.06710 | P=0.7987 | | | | | | | | **OxtR-/-** | 0.4419 | 1.0198 | |
| | | | | | Stim | | F (1, 17) = 5.841 | P=0.0272 | | | | | | | | | | | |
| | | | | | Subject | | F (17, 17) = 0.9432 | P=0.5472 | | | | | | | | | | | |
| | | % of assay time spent in side-by-side contact | None | 2 Way RM ANOVA | Genotype*Stim | Subject | F (1, 17) = 4.721 | P=0.0442 | | WT | Partner | OxtR-/- | Partner | | 0.008 | **WT** | 54.26753176 | 1.036155721 | |
| | | | | | Genotype | | F (1, 17) = 3.345 | P=0.0850 | | WT | Stranger | OxtR-/- | Stranger | | 0.4141 | **OxtR-/-** | 39.27973307 | 4.591887409 | |
| | | | | | Stim | | F (1, 17) = 39.73 | P<0.0001 | | WT | Partner | WT | Stranger | | <0.0001 | | | | |

| | | | | | | | | | | | | | | | | | | |
|---|---|---|---|---|---|---|---|---|---|---|---|---|---|---|---|---|---|---|
| S7 | | | | | Subject | | F (17, 17) = 0.4212 | P=0.9582 | | OxtR-/- | Partner | OxtR-/- | Stranger | | 0.0112 | | | |
| | b | Attacks on stranger | None | GLM with a two-parameter generalized poisson distribution | Genotype | None | Z(17) = 1.000 | 0.31732 | | | | | | | | | | |
| | d | Peak z scored dF/F | Box Cox | LME | Stim | Vole | F(1,1354) = 8.169 | 0.0043266 | | | | | | | | **WT** | 2.7039 | 2.373 |
| | | | | | Group | | F(2,1354) = 0.63889 | 0.52803 | | | | | | | | **Ppref** | 3.6348 | 2.4499 |
| | | | | | Stim*Group | | F(2,1354) = 0.12453 | 0.88292 | | | | | | | | **Spref** | 3.4087 | 2.3369 |
| | | AUC of z scored dF/F | Box Cox | LME | Stim | Vole | F(1,1354) =15.785 | 7.47E-05 | | | | | | | | **WT** | 192.7423 | 86.2053 |
| | | | | | Group | | F(2,1354) = 0.72701 | 0.48354 | | | | | | | | **Ppref** | 283.6434 | 123.4179 |
| | | | | | Stim*Group | | F(2,1354) = 0.046833 | 0.95425 | | | | | | | | **Spref** | 164.2437 | 118.291 |
| | f | Peak z scored dF/F | Box Cox | LME | Stim | Vole | F(1,1383) = 6.9762 | 0.008353 | | | | | | | | **WT** | 2.2399 | 2.5964 |
| | | | | | Group | | F(2,1383) = 0.0071919 | 0.99283 | | | | | | | | **Ppref** | 2.2094 | 2.9308 |
| | | | | | Stim*Group | | F(2,1383) = 0.071645 | 0.93087 | | | | | | | | **Spref** | 2.5391 | 3.025 |
| | | AUC of z scored dF/F | Box Cox | LME | Stim | Vole | F(1,1383) = 8.5472 | 0.0035168 | | | | | | | | **WT** | 107.6244 | 208.9991 |
| | | | | | Group | | F(2,1383) = 0.10878 | 0.89693 | | | | | | | | **Ppref** | 183.2685 | 240.2516 |
| | | | | | Stim*Group | | F(2,1383) = 0.55498 | 0.57421 | | | | | | | | **Spref** | 173.4334 | 318.5029 |
| | l | Peak z scored dF/F | Box Cox | LME | Stim | Vole | F(1,2094) = 8.0273 | 0.0046519 | | WT | Partner | WT | Stranger | 0.0013263 | 0.011874 | **WT** | 2.5071 | 1.9606 |
| | | | | | Group | | F(2,2094) = 4.0128 | 0.018222 | | WT | Partner | PPref | Partner | 0.12586 | 0.70198 | **Ppref** | 2.8249 | 1.9555 |
| | | | | | Stim*Group | | F(2,2094) = 4.0205 | 0.018083 | | WT | Partner | SPref | Partner | 0.0086734 | 0.075407 | **Spref** | 1.8994 | 1.8549 |
| | | | | | | | | | | WT | Stranger | PPref | Stranger | 0.3452 | 0.97787 | | | |
| | | | | | | | | | | WT | Stranger | SPref | Stranger | 0.37948 | 0.98636 | | | |
| | | | | | | | | | | PPref | Partner | PPref | Stranger | 0.00014643 | 0.0013171 | | | |
| | | | | | | | | | | PPref | Partner | SPref | Partner | 0.0013708 | 0.01227 | | | |
| | | | | | | | | | | PPref | Stranger | SPref | Stranger | 0.86416 | 1 | | | |
| | | | | | | | | | | SPref | Partner | SPref | Stranger | 0.56992 | 0.9995 | | | |
| | m | AUC of z scored dF/F | Box Cox | LME | Stim | Vole | F(1,2094) = 22.637 | 2.09E-06 | | WT | Partner | WT | Stranger | 2.70E-07 | 2.43E-06 | **WT** | 131.071 | 11.91 |
| | | | | | Group | | F(2,2094) = 3.8898 | 0.020597 | | WT | Partner | PPref | Partner | 0.034164 | 0.26864 | **Ppref** | 215.0337 | 8.7216 |
| | | | | | Stim*Group | | F(2,2094) = 4.8412 | 0.0079862 | | WT | Partner | SPref | Partner | 0.024732 | 0.20179 | **Spref** | -30.8966 | 28.5899 |
| | | | | | | | | | | WT | Stranger | PPref | Stranger | 0.778 | 1 | | | |
| | | | | | | | | | | WT | Stranger | SPref | Stranger | 0.93276 | 1 | | | |
| | | | | | | | | | | PPref | Partner | PPref | Stranger | 3.35E-07 | 3.01E-06 | | | |
| | | | | | | | | | | PPref | Partner | SPref | Partner | 0.0015036 | 0.013452 | | | |
| | | | | | | | | | | PPref | Stranger | SPref | Stranger | 0.74138 | 0.99999 | | | |
| | | | | | | | | | | SPref | Partner | SPref | Stranger | 0.95165 | 1 | | | |
| | o | Peak z scored dF/F | Box Cox | LME | Stim | Vole | F(1,1219) = 3.7933 | 0.051688 | | | | | | | | **WT** | 2.321 | 1.9035 |
| | | | | | Group | | F(2,1219) = 1.5263 | 0.21776 | | | | | | | | **Ppref** | 2.4067 | 2.4721 |
| | | | | | Stim*Group | | F(2,1219) = 0.49648 | 0.60879 | | | | | | | | **Spref** | 2.1985 | 1.9782 |
| | p | AUC of z scored dF/F | Box Cox | LME | Stim | Vole | F(1,1219) = 6.5923 | 0.010361 | | | | | | | | **WT** | 134.6529 | -0.4092 |
| | | | | | Group | | F(2,1219) = 0.97463 | 0.37763 | | | | | | | | **Ppref** | 148.73 | 59.1593 |
| | | | | | Stim*Group | | F(2,1219) = 0.12033 | 0.88664 | | | | | | | | **Spref** | 99.2955 | 18.6906 |

| **Figure** | **Panel** | **Dependent Variable** | **Transformation applied** | **Test** | **Independent variable(s)** | **Random effect(s)** | **Test Statistic** | **p-value** | **Notes** | **Multiple Comparisons** | | | | | | | | | |
|---|---|---|---|---|---|---|---|---|---|---|---|---|---|---|---|---|---|---|---|
| | | | | | | | | | | **Comparison group 1** | | **Comparison group 2** | | **Unadjusted p value** | **Sidak corrected p** | **Median category** | **Median WT** | **Median Oxtr-/- (all) or Oxtr-/- Ppref** | **Median Oxtr-/- Spref** |
| | a | % of assay spent non-AG sniffing | None | LME | Group | Vole | F (2, 15) = 1.305 | 0.3003 | | | | | | | | **Intro** | 2.791138889 | 6.055891526 | 4.007782966 |
| | | | | | Assay | | F (2, 25) = 2.589 | 0.0951 | | | | | | | | **Reunion** | 3.737083165 | 2.577055556 | 2.32 |
| | | | | | Group x Assay | | F (4, 25) = 2.083 | 0.1132 | | | | | | | | **Stranger exposure** | 1.541541667 | 2.256541667 | 4.776 |
| | b | % of assay spent AG sniffing | None | LME | Group | Vole | F (2, 15) = 0.9515 | 0.4083 | | WT | Intro | Ppref | Intro | 0.0176 | 0.273 | **Intro** | 1.758694444 | 5.534777778 | 2.673615744 |
| | | | | | Assay | | F (2, 25) = 0.2064 | 0.8149 | | WT | Intro | Spref | Intro | 0.8644 | >0.9999 | **Reunion** | 3.273305556 | 3.533444444 | 3.358277778 |
| | | | | | Group x Assay | | F (4, 25) = 3.856 | 0.0142 | | Ppref | Intro | Spref | Intro | 0.0762 | 0.7599 | **Stranger exposure** | 4.45475 | 2.668038301 | 4.985833333 |
| | | | | | | | | | | WT | Reunion | Ppref | Reunion | 0.1191 | 0.898 | | | | |
| | | | | | | | | | | WT | Reunion | Spref | Reunion | 0.9971 | >0.9999 | | | | |
| | | | | | | | | | | Ppref | Reunion | Spref | Reunion | 0.2256 | 0.99 | | | | |
| | | | | | | | | | | WT | Stranger Exp | PPref | Stranger Exp | 0.3727 | 0.9998 | | | | |
| | | | | | | | | | | WT | Stranger Exp | SPref | Stranger Exp | 0.3139 | 0.9989 | | | | |
| | | | | | | | | | | Ppref | Stranger Exp | Spref | Stranger Exp | 0.1216 | 0.903 | | | | |
| | | | | | | | | | | WT | Intro | WT | Reunion | 0.5917 | >0.9999 | | | | |
| | | | | | | | | | | WT | Intro | WT | Stranger Exp | 0.0653 | 0.7032 | | | | |
| | | | | | | | | | | WT | Reunion | WT | Stranger Exp | 0.2258 | 0.99 | | | | |
| | | | | | | | | | | Ppref | Intro | PPref | Reunion | 0.3938 | 0.9999 | | | | |
| | | | | | | | | | | Ppref | Intro | Ppref | Stranger Exp | 0.0141 | 0.2251 | | | | |
| | | | | | | | | | | Ppref | Reunion | Ppref | Stranger Exp | 0.053 | 0.6246 | | | | |
| | | | | | | | | | | Spref | Intro | Spref | Reunion | 0.9038 | >0.9999 | | | | |
| | | | | | | | | | | Spref | Intro | Spref | Stranger Exp | 0.0544 | 0.6343 | | | | |
| | | | | | | | | | | Spref | Reunion | Spref | Stranger Exp | 0.0695 | 0.7265 | | | | |
| | c | % of assay spent in side-by-side contact | None | LME | Group | Vole | F (2, 15) = 0.7959 | 0.4693 | | Intro | | Reunion | | | <0.0001 | **Intro** | 6.455598912 | 8.889888889 | 22.01116667 |
| | | | | | Assay | | F (2, 25) = 62.37 | <0.0001 | | Intro | | Stranger Exp | | | 0.9708 | **Reunion** | 57.51216667 | 48.24438889 | 81.71772222 |
| | | | | | Group x Assay | | F (4, 25) = 0.8275 | 0.5202 | | Reunion | | Stranger Exp | | | <0.0001 | **Stranger exposure** | 1.933458333 | 12.14811741 | 6.367166667 |
| | d | % of assay time spent in mating behavior | None | t test | Genotype | n/a | t(10) = 1.961 | 0.0783 | | | | | | | | | 4.232 | 0.2988 | |
| | | Number of mating bouts | None | GLM with a two-parameter generalized poisson distribution | Genotype | n/a | Z(10) = 1.232 | 0.218 | | | | | | | | | 9 | 3 | |
| | e | Number of strikes (Only comparing WT Intro to WT Stranger Exp) | None | GLM with 2 parameter Poisson distribution | Assay | None | F(1,18) = -1.445 | 0.14838 | | | | | | | | | | | |
| | g | No comparisons made | | | | | | | | | | | | | | | | | |
| | h | No comparisons made | | | | | | | | | | | | | | | | | |

| | | | | | | | | | | | | | | | | | | | |
|---|---|---|---|---|---|---|---|---|---|---|---|---|---|---|---|---|---|---|---|
| | j | Peak z scored dF/F | Box Cox | LME | Group | Vole | F(2,1458) = 3.36 | 0.035006 | | WT | | PPref | | 0.18823 | 0.46506 | **Intro** | 2.2028 | 1.8874 | 2.2343 |
| | | | | | Assay | | F(2,1458) = 0.51377 | 0.59834 | | WT | | SPref | | 0.029897 | 0.087036 | **Reunion** | 2.362 | 2.4848 | 2.0366 |
| | | | | | Assay*Group | | F(4,1458) = 1.649 | 0.15948 | | PPref | | SPref | | 0.37589 | 0.7569 | **Stranger exposure** | 2.642 | 2.0908 | 2.0945 |
| | k | AUC of z scored dF/F | Box Cox | LME | Group | Vole | F(2,1458) = 6.8966 | 0.0010445 | | WT | | PPref | | 0.055624 | 0.15776 | **Intro** | 74.769 | 35.1572 | 83.7332 |
| | | | | | Assay | | F(2,1458) = 3.7174 | 0.024529 | | WT | | SPref | | 0.00094341 | 0.0028276 | **Reunion** | 177.4331 | 140.7544 | 60.5692 |
| | | | | | Assay*Group | | F(4,1458) = 2.0888 | 0.079977 | | PPref | | SPref | | 0.2 | 0.48799 | **Stranger exposure** | 126.7499 | 84.0099 | -8.4426 |
| | | | | | | | | | | Intro | | Reunion | | 0.0017304 | 0.0051821 | | | | |
| | | | | | | | | | | Intro | | Stranger Exp | | 0.38158 | 0.76349 | | | | |
| | | | | | | | | | | Reunion | | Stranger Exp | | 0.032906 | 0.095506 | | | | |
| | m | Peak z scored dF/F | Box Cox | LME | Group | Vole | F(2,843) = 0.40446 | 0.66747 | | Intro | | Reunion | | 0.017762 | 0.052345 | | 2.6225 | 2.1131 | 2.7406 |
| | | | | | Assay | | F(2,843) = 3.4396 | 0.032529 | | Intro | | Stranger Exp | | 0.44973 | 0.83338 | | 2.5961 | 3.5185 | 3.3379 |
| | | | | | Assay*Group | | F(4,843) = 1.7278 | 0.14176 | | Reunion | | Stranger Exp | | 0.0069082 | 0.020582 | | 2.4298 | 2.2098 | 2.1879 |
| | n | AUC of z scored dF/F | Box Cox | LME | Group | Vole | F(2,843) = 0.25972 | 0.77133 | | WT | Intro | WT | SepReu | 0.060571 | 0.67525 | **Intro** | 111.8825 | 21.5275 | 143.8561 |
| | | | | | Assay | | F(2,843) = 8.5515 | 0.00021053 | | WT | Intro | WT | Stranger Exp | 0.7955 | 1 | **Reunion** | 189.1582 | 322.6815 | 309.2118 |
| | | | | | Assay*Group | | F(4,843) = 2.6395 | 0.032728 | | WT | Intro | PPref | Intro | 0.030794 | 0.4305 | **Stranger exposure** | 77.8033 | 48.8651 | 87.3153 |
| | | | | | | | | | | WT | Intro | SPref | Intro | 0.1781 | 0.97071 | | | | |
| | | | | | | | | | | WT | SepReu | WT | Stranger Exp | 0.067542 | 0.716 | | | | |
| | | | | | | | | | | WT | SepReu | PPref | SepReu | 0.084827 | 0.79721 | | | | |
| | | | | | | | | | | WT | SepReu | SPref | SepReu | 0.80842 | 1 | | | | |
| | | | | | | | | | | WT | Stranger Exp | PPref | Stranger Exp | 0.3712 | 0.99976 | | | | |
| | | | | | | | | | | WT | Stranger Exp | SPref | Stranger Exp | 0.60229 | 1 | | | | |
| | | | | | | | | | | PPref | Intro | PPref | SepReu | 4.09E-07 | 7.37E-06 | | | | |
| | | | | | | | | | | PPref | Intro | PPref | Stranger Exp | 0.87814 | 1 | | | | |
| | | | | | | | | | | PPref | Intro | SPref | Intro | 0.0085104 | 0.14259 | | | | |
| | | | | | | | | | | PPref | SepReu | PPref | Stranger Exp | 0.0012947 | 0.02305 | | | | |
| | | | | | | | | | | PPref | SepReu | SPref | SepReu | 0.34059 | 0.99944 | | | | |
| | | | | | | | | | | PPref | Stranger Exp | SPref | Stranger Exp | 0.63846 | 1 | | | | |
| | | | | | | | | | | SPref | Intro | SPref | SepReu | 0.67247 | 1 | | | | |
| | | | | | | | | | | SPref | Intro | SPref | Stranger Exp | 0.13101 | 0.92016 | | | | |
| | | | | | | | | | | SPref | SepReu | SPref | Stranger Exp | 0.12728 | 0.91375 | | | | |
| | o | % time spent in social bouts during the intro | None | t test | Genotype | None | t(15) = 4.046 | 0.0011 | | | | | | | | | 13.25 | 28.62 | |
| | p | % of assay time spent non-AG sniffing | None | Permutation test on F statistic | Group | n/a | Observed F(2,13) = 3.1087 | 0.0739 | | WT | | Ppref | | 0.0095 | 0.02823 | | 2.791138889 | 6.055891526 | 4.007782966 |
| | | | | | | | | | | WT | | Spref | | 0.1594 | 0.40603 | | | | |
| | | | | | | | | | | Ppref | | Spref | | 0.775 | 0.98861 | | | | |
| | | % of assay time spent AG sniffing | None | Permutation test on F statistic | Group | n/a | Observed F(2,13) = 4.955 | 0.0283 | 10,000 iterations | WT | | Ppref | | 0.0107 | 0.031758 | | 1.758694444 | 5.534777778 | 2.673615744 |
| | | | | | | | | | | WT | | Spref | | 0.8024 | 0.99228 | | | | |
| | | | | | | | | | | Ppref | | Spref | | 0.12778 | 0.33644 | | | | |
| | | % of assay time spent in side-by-side contact | None | Permutation test on F statistic | Group | n/a | Observed F(2,13) = 8.6338 | 0.0062 | 10,000 iterations | WT | | PPref | | 0.4441 | 0.82821 | | 6.455598912 | 8.889888889 | 22.01116667 |
| | | | | | | | | | | WT | | SPref | | 0.0081 | 0.024104 | | | | |
| | | | | | | | | | | PPref | | SPref | | 0.11528 | 0.3075 | | | | |
| | q | Transition probability | None | Permutation test on F statistic | Genotype | None | Non-AG sniff --> AG sniff: F(2,13) = 1.43991340732870 | 2.75E-01 | | | | | | | 0.9788 | | | | |
| | | | | | Genotype | None | Non-AG sniff --> Side-by-side: F(2,13) = 3.58512032922330 | 0.0569 | | | | | | | 0.5049 | | | | |
| | | | | | Genotype | None | Non-AG sniff --> No interaction: F(2,13) = 3.57330620058973 | 0.0566 | | | | | | | 0.503 | | | | |
| | | | | | Genotype | None | AG sniff --> Non-AG sniff: F(2,13) = 2.28109495011194 | 1.28E-01 | | | | | | | 0.8062 | | | | |
| | | | | | Genotype | None | AG sniff --> Side-by-side: F(2,13) = 1.75010845473656 | 2.26E-01 | | | | | | | 0.9534 | | | | |
| | | | | | Genotype | None | AG sniff --> No interaction: F(2,13) = 1.48155671372798 | 2.46E-01 | | | | | | | 0.9661 | | | | |
| | | | | | Genotype | None | Side-by-side --> Non-AG sniff: F(2,13) = 1.36013247181587 | 2.90E-01 | | | | | | | 0.9836 | | | | |
| S8 | | | | | Genotype | None | Side-by-side --> AG sniff: F(2,13) = 2.23717796011578 | 1.47E-01 | | | | | | | 0.8512 | | | | |
| | | | | | Genotype | None | Side-by-side --> No interaction: F(2,13) = 5.66318891976579 | 1.27E-02 | | | | | | | 0.1422 | | | | |
| | | | | | Genotype | None | No interaction --> Non-AG sniff: F(2,13) = 0.131503761927990 | 8.92E-01 | | | | | | | 1 | | | | |
| | | | | | Genotype | None | No interaction --> AG sniff: F(2,13) = 2.29041426349541 | 0.1286 | | | | | | | 0.8083 | | | | |
| | | | | | Genotype | None | No interaction --> Side-by-side: F(2,13) = 1.07568177754074 | 0.3555 | | | | | | | 0.9949 | | | | |
| | | Left = WT; Pearson correlations between intro behavior (% of social contact) and PPT side-by-side contact | None | Pearson correlation | Intro Non-AG sniff % vs. PPT Partner side-by-side | n/a | r = -0.211790912095149 | 0.556935236 | | | | | | | | | | | |
| | | | | | Intro AG sniff % vs. PPT Partner side-by-side | n/a | r = -0.259088860096925 | 0.469780022 | | | | | | | | | | | |
| | | | | | Intro side-by-side % vs. PPT Partner side-by-side | n/a | r = 0.268601335606747 | 0.453021976 | | | | | | | | | | | |
| | | | | | Intro Non-AG sniff % vs. PPT Stranger side-by-side | n/a | r = 0.445349117231156 | 0.197110273 | | | | | | | | | | | |

| | | | | | | | | |
|---|---|---|---|---|---|---|---|---|
| r | | | | Intro AG sniff % vs. PPT Stranger side-by-side | n/a | r = 0.0527829986310332 | 0.884858338 | |
| | | | | Intro side-by-side % vs. PPT Stranger side-by-side | n/a | r = -0.237610642018951 | 0.508591771 | |
| | Right = OxtR-/-; Pearson correlations between intro behavior (% of social contact) and PPT side-by-side contact | None | Pearson correlation | Intro Non-AG sniff % vs. PPT Partner side-by-side | n/a | r = 0.58237266892227 | 0.22519915 | |
| | | | | Intro AG sniff % vs. PPT Partner side-by-side | n/a | r = 0.921368200517741 | 0.009031351 | |
| | | | | Intro side-by-side % vs. PPT Partner side-by-side | n/a | r = -0.87985358183197 | 0.020785576 | |
| | | | | Intro Non-AG sniff % vs. PPT Stranger side-by-side | n/a | r = -0.830552779829332 | 0.040635926 | |
| | | | | Intro AG sniff % vs. PPT Stranger side-by-side | n/a | r = -0.809429699980646 | 0.051015084 | |
| | | | | Intro side-by-side % vs. PPT Stranger side-by-side | n/a | r = 0.881864967809125 | 0.020109489 | |
| s | OxtR-/- including the single male with no partner preference; Pearson correlations between intro behavior (% of social contact) and PPT side-by-side contact | None | Pearson correlation | Intro Non-AG sniff % vs. PPT Partner side-by-side | n/a | r = 0.615666518812657 | 0.141061052 | |
| | | | | Intro AG sniff % vs. PPT Partner side-by-side | n/a | r = 0.924807973725302 | 0.002858651 | |
| | | | | Intro side-by-side % vs. PPT Partner side-by-side | n/a | r = -0.881165802837682 | 0.008761617 | |
| | | | | Intro Non-AG sniff % vs. PPT Stranger side-by-side | n/a | r = -0.749206084232941 | 0.05257222 | |
| | | | | Intro AG sniff % vs. PPT Stranger side-by-side | n/a | r = -0.772965961902528 | 0.041563072 | |
| | | | | Intro side-by-side % vs. PPT Stranger side-by-side | n/a | r = 0.819273228472236 | 0.024133092 | |
| t | Left: X = % of intro engaged in AG sniffing; Y = % of PPT spent in side-by-side contact with partner | None | Linear regression | n/a | n/a | R squared = 0.8553; F(1,5) = 29.55 | 0.0029 | Y = 1.366*X - 3.925 |
| | Right: X = % of intro engaged in side-by-side contact; Y = % of PPT spent in side-by-side contact with partner | None | Linear regression | n/a | n/a | R squared = 0.7765; F(1,5) = 17.37 | 0.0088 | Y = -0.9345*X + 79.16 |
| u | Left: X: Intro mean AG sniff Box Cox normalized AUC; Y: Mean % of PPT engaged in side-by-side contact with partner | None | Linear regression | n/a | n/a | R squared =0.5749; F(1,5) = 6.763 | 0.0482 | Y = -5.019*X + 355.6 |
| | Right: X: Intro mean AG sniff Box Cox normalized AUC; Y: Mean % of PPT engaged in side-by-side contact with stranger | None | Linear regression | n/a | n/a | R squared = 0.2680; F(1,5) = 1.831 | 0.234 | Y = 1.166*X - 67.12 |
| v | Left: X: Intro mean AG sniff Box Cox normalized AUC; Y: Mean PPT Box Cox normalized partner social bout-elicited AUC | None | Linear regression | n/a | n/a | R squared = 0.01949; F(1,5) = 0.09936 | 0.7653 | Y = -0.05104*X + 50.65 |
| | Right: X: Intro mean AG sniff Box Cox normalized AUC; Y: Mean PPT Box Cox normalized stranger social bout-elicited AUC | None | Linear regression | n/a | n/a | R squared = 0.1465; F(1,5) = 0.8583 | 0.3967 | Y = 0.08313*X + 40.40 |
| w | Left: X: Intro mean sniff to huddle transition Box Cox normalized peak; Y: Mean % of PPT engaged in side-by-side contact with partner | None | Linear regression | n/a | n/a | R squared =0.8499; F(1,5) = 28.31 | 0.0031 | Y = -198.6*X + 335.4 |
| | Right: X: Intro mean sniff to huddle transition Box Cox normalized peak; Y: Mean % of PPT engaged in side-by-side contact with stranger | None | Linear regression | n/a | n/a | R squared =0.7330; F(1,5) = 13.73 | 0.0139 | Y = 62.72*X - 88.52 |
| x | Left: X: Intro mean sniff to huddle transition Box Cox normalized peak; Y: Mean PPT Box Cox normalized partner social bout-elicited AUC | None | Linear regression | n/a | n/a | R squared = 0.2177; F(1,5) = 1.391 | 0.2913 | Y = -5.552*X + 55.99 |
| | Right: X: Intro mean sniff to huddle transition Box Cox normalized peak; Y: Mean PPT Box Cox normalized stranger social bout-elicited AUC | None | Linear regression | n/a | n/a | R squared = 0.1105; F(1,5) = 0.6212 | 0.4663 | Y = 2.350*X + 42.21 |
| y | Left: X: Intro mean sniff to huddle transition Box Cox normalized peak; Y: Mean % of PPT engaged in side-by-side contact with partner | None | Linear regression | n/a | n/a | R squared = 0.02264; F(1,8) = 0.1853 | 0.6782 | Y = 7.295*X + 44.54 |
| | Right: X: Intro mean sniff to huddle transitionBox Cox normalized peak; Y: Mean % of PPT engaged in side-by-side contact with stranger | None | Linear regression | n/a | n/a | R squared = 0.3230; F(1,8) = 3.817 | 0.0865 | Y = 3.953*X - 4.703 |
| z | Left: X: Intro mean sniff to huddle transition Box Cox normalized peak; Y: Mean % of PPT engaged in side-by-side contact with partner | None | Linear regression | n/a | n/a | R squared =0.8808; F(1,4) = 29.56 | 0.0056 | Y = -220.4*X + 367.6 |
| | Right: X: Intro mean sniff to huddle transition Box Cox normalized peak; Y: Mean % of PPT engaged in side-by-side contact with stranger | None | Linear regression | n/a | n/a | R squared =0.9049; F(1,4) = 38.08 | 0.0035 | Y = 78.45*X - 111.7 |
| aa | Left: X: Intro mean sniff to huddle transition Box Cox normalized peak; Y: Mean PPT Box Cox normalized partner social bout-elicited AUC | None | Linear regression | n/a | n/a | R squared = 0.07793; F(1,8) = 0.6761 | 0.4348 | Y = 1.381*X + 44.27 |

| | | | | | | | | | |
|---|---|---|---|---|---|---|---|---|---|
| | aa | Right: X: Intro mean sniff to huddle transition Box Cox normalized peak; Y: Mean PPT Box Cox normalized stranger social bout-elicited AUC | None | Linear regression | n/a | n/a | R squared = 0.0005291; F(1,8) = 0.004235 | 0.9497 | Y = -0.1368*X + 45.85 |
| | bb | Left: X: Intro mean sniff to huddle transition Box Cox normalized peak; Y: Mean PPT Box Cox normalized partner social bout-elicited AUC | None | Linear regression | n/a | n/a | R squared = 0.7015; F(1,4) = 9.398 | 0.0375 | Y = -10.08*X + 62.67 |
| | bb | Right: X: Intro mean sniff to huddle transition Box Cox normalized peak; Y: Mean PPT Box Cox normalized stranger social bout-elicited AUC | None | Linear regression | n/a | n/a | R squared = 0.003073; F(1,4) = 0.01233 | 0.9169 | Y = 0.3386*X + 45.17 |

| Figure | Panel | Dependent Variable | Transformation applied | Test | Independent variable(s) | Random effect(s) | Test Statistic | p-value | Notes | Multiple Comparisons | | | | | | |
|---|---|---|---|---|---|---|---|---|---|---|---|---|---|---|---|---|
| | | | | | | | | | | Comparison group 1 | Comparison group 2 | Unadjusted p value | Sidak corrected p | Median WT | Median OxtR-/- (Ppref) | Median Oxtr-/- Spref |
| S9 | b | AUC of dF/F | Box Cox | LME | Time (Pre/Post) | Vole; Trace | F(1,1118) = 129.39 | 1.90E-28 | | | | | | | | |
| | c | Difference in pre/post AUC by animal | None | One sample t test | n/a | None | t(36) = -6.8647 | 4.94E-08 | | | | | | | | |
| | d | Median dF/F | Box Cox | LME | Time (Pre/Post) | Vole; Trace | F(1,1118) = 110.38 | 1.10E-24 | | | | | | | | |
| | e | Difference in pre/post median by animal | None | One sample t test | n/a | None | t(36) = -6.7933 | 6.14E-08 | | | | | | | | |
| | g | dF/F | Box Cox | Repeated LMEs at each time point comparing mean dF/F to the pre-assay (-10s to 0s) mean dF/F value of 0.0237 | n/a | Vole | n/a | n/a | Benjamini-Yekutieli correction applied to resulting p values | | | | | | | |
| | i | Peak z scored dF/F | Box Cox | LME | Genotype | None | F(1,15) = 2.4348 | 1.40E-01 | | | | | | 3.8632 | 8.0665 | |
| | j | Z scored dF/F AUC | Box Cox | LME | Genotype | None | F(1,15) = 0.21366 | 6.51E-01 | | | | | | 1.21E+03 | 1.29E+04 | |
| | l | Peak z scored dF/F | Box Cox | LME | Group | None | F(2,11) = 0.90879 | 4.31E-01 | | | | | | 4.0634 | 3.8172 | 5.4219 |
| | m | Z scored dF/F AUC | Box Cox | LME | Group | None | F(2,11) = 1.2492 | 3.24E-01 | | | | | | 1.27E+03 | -1.42E+03 | 4.22E+03 |

### Table 2: Voles utilized in this study

| Animal | Sex | Genotype | Intro | Timed Mating | PPT | Separation/Reunification | Stranger Exposure |
|---|---|---|---|---|---|---|---|
| V1259 | M | WT | Included | Included | Excluded. Experimental failure. Inccorect behavior timeline. | Excluded. Experimental failure. Inccorect behavior timeline. | Excluded. Experimental failure. Inccorect behavior timeline. |
| V1618 | M | WT | Excluded. Equipment failure. Camera was not centered over the cage. | Included | Excluded. Experimental failure. Inccorect behavior timeline. | Excluded. Experimental failure. Inccorect behavior timeline. | Excluded. Experimental failure. Inccorect behavior timeline. |
| V2210 | M | WT | Included | Included | Included | Excluded. Data lost. | Included |
| V2211 | M | WT | Included | Included | Included | Excluded. Data lost. | Included |
| V3950 | M | WT | Included | Included | Included | Included | Included |
| V5473 | M | WT | Included | Included | Included | Included | Included |
| V6074 | M | WT | Included | Included | Included | Included | Included |
| V6077 | M | WT | Included | Included | Included | Included | Included |
| V6078 | M | WT | Included | Included | Included | Included | Included |
| V6080 | M | WT | Included | Included | Included | Included | Included |
| V9935 | M | WT | Included | Included | Included | Included | Included |
| V9936 | M | WT | Included | Included | Included | Included | Included |
| B1627 | M | MU | Included | Included | Included | Included | Included |
| B1664 | M | MU | Included | Included | Included | Included | Included |
| B1669 | M | MU | Excluded. Behavioral outlier. 3.45 scaled median absolute deviations below the median for total % of assay time spent in social bouts. | Included | Included | Included | Included |
| V3974 | M | MU | Excluded. Behavioral outlier. 3.27 scaled median absolute deviations below the median for total % of assay time spent in social bouts. | Included | Excluded. Experimental equipment failure. Rotary joint placed too high for the first 25 minutes; restricted motion. 2 severe tangles with the tethered partner. | Included | Included |
| V3975 | M | MU | Included | Included | Included | Included | Included |
| V3977 | M | MU | Included | Included | Excluded. Poor signal quality. | Included | Excluded. Poor signal quality. |
| V4112 | M | MU | Excluded. Poor signal quality. | Included | Included | Included | Excluded. Poor signal quality. |
| V5161 | M | MU | Included | Included | Included | Included | Included |
| V5168 | M | MU | Included | Included | Included | Included | Included |
| V5456 | M | MU | Included | Included | Included | Included | Included |
| V5457 | M | MU | Included | Included | Included | Included | Included |
| V5488 | M | MU | Included | Excluded. Poor signal quality. Excessive chewing of the fiber by the partner. | Excluded. Poor signal quality. | Included | Included |
| V3580 | F | WT | Included | Excluded. Implant failure. | Included | Excluded. Implant failure. | Excluded. Implant failure. |
| V3582 | F | WT | Excluded. Poor signal quality. | Included | Included | Included | Included |
| V3583 | F | WT | Included | Included | Excluded. Poor signal quality. | Included | Included |
| V3812 | F | WT | Included | Included | Included | Included | Included |
| V4011 | F | WT | Included | Included | Excluded. Behavioral outlier. 4.44 scaled median absolute deviations below the median for (partner % of assay time spent in social bouts) - (stranger % of assay time spent in social bouts). | Included | Included |
| V4480 | F | WT | Included | Included | Included | Included | Included |

| | | | | | | | |
|---|---|---|---|---|---|---|---|
| **V4485** | F | WT | Excluded. Poor signal quality. | Excluded. Poor signal quality. | Included | Included | Included |
| **V4494** | F | WT | Included | Included | Included | Included | Included |
| **V5673** | F | WT | Included | Included | Included | Included | Included |
| **V5678** | F | WT | Included | Included | Included | Included | Included |
| **B1625** | F | MU | Included | Included | Included | Included | Included |
| **B1661** | F | MU | Included | Included | Excluded. Behavioral outlier. 4.65 scaled median absolute deviations below the median for (partner % of assay time spent in social bouts) - (stranger % of assay time spent in social bouts). | Included | Included |
| **V3878** | F | MU | Included | Included | Included | Included | Included |
| **V4058** | F | MU | Included | Included | Included | Included | Included |
| **V4064** | F | MU | Excluded. Behavioral outlier. 3.5 scaled median absolute deviations above the median for total % of assay time spent in social bouts. | Excluded. Poor signal quality. | Included | Included | Included |
| **V4528** | F | MU | Included | Included | Included | Included | Included |
| **V6403** | F | MU | Included | Included | Included | Included | Included |
| **V6413** | F | MU | Included | Included | Included | Included | Included |
| **V8540** | F | MU | Included | Included | Included | Included | Included |
| **V8546** | F | MU | Excluded. Behavioral outlier. 4.39 scaled median absolute deviations above the median for total % of assay time spent in social bouts. | Included | Included | Included | Included |
| **V8547** | F | MU | Included | Included | Included | Included | Included |
