## Extended Data for "Oxytocin receptor function regulates neural signatures of pair bonding and fidelity in the nucleus accumbens"

Kimberly L. P. Long[1], Nerissa E. G. Hoglen[1,2], Alex J. Keip[1,2], Robert M. Klinkel[1], DéJenaé L. See[1], Joseph Maa[3], Jenna C. Wong[3], Michael Sherman[1], and Devanand S. Manoli[1]*

[1]Department of Psychiatry and Behavioral Sciences, Center for Integrative Neuroscience, Weill Institute for Neurosciences, and Kavli Institute for Fundamental Neuroscience, University of California, San Francisco; San Francisco, CA 95158, USA.

[2]Neurosciences Graduate Program, University of California, San Francisco; San Francisco, CA 95158, USA.

[3]Department of Molecular and Cell Biology, University of California, Berkeley; Berkeley CA 94720, USA.

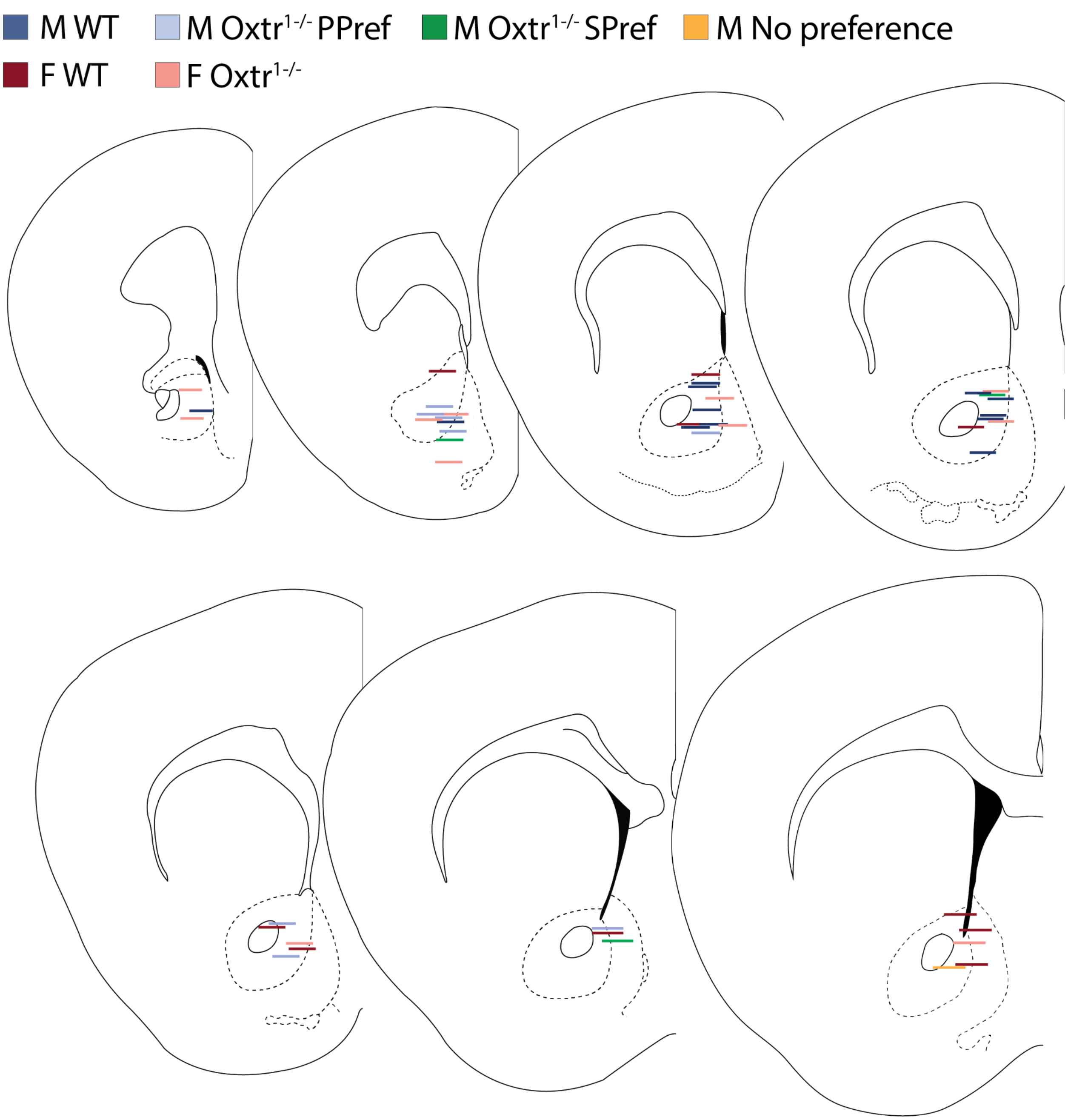


**Extended Data Figure 1: Fiber locations in the prairie vole nucleus accumbens.** M, male; F, female; PPref, partner-preferring; SPref, stranger-preferring.

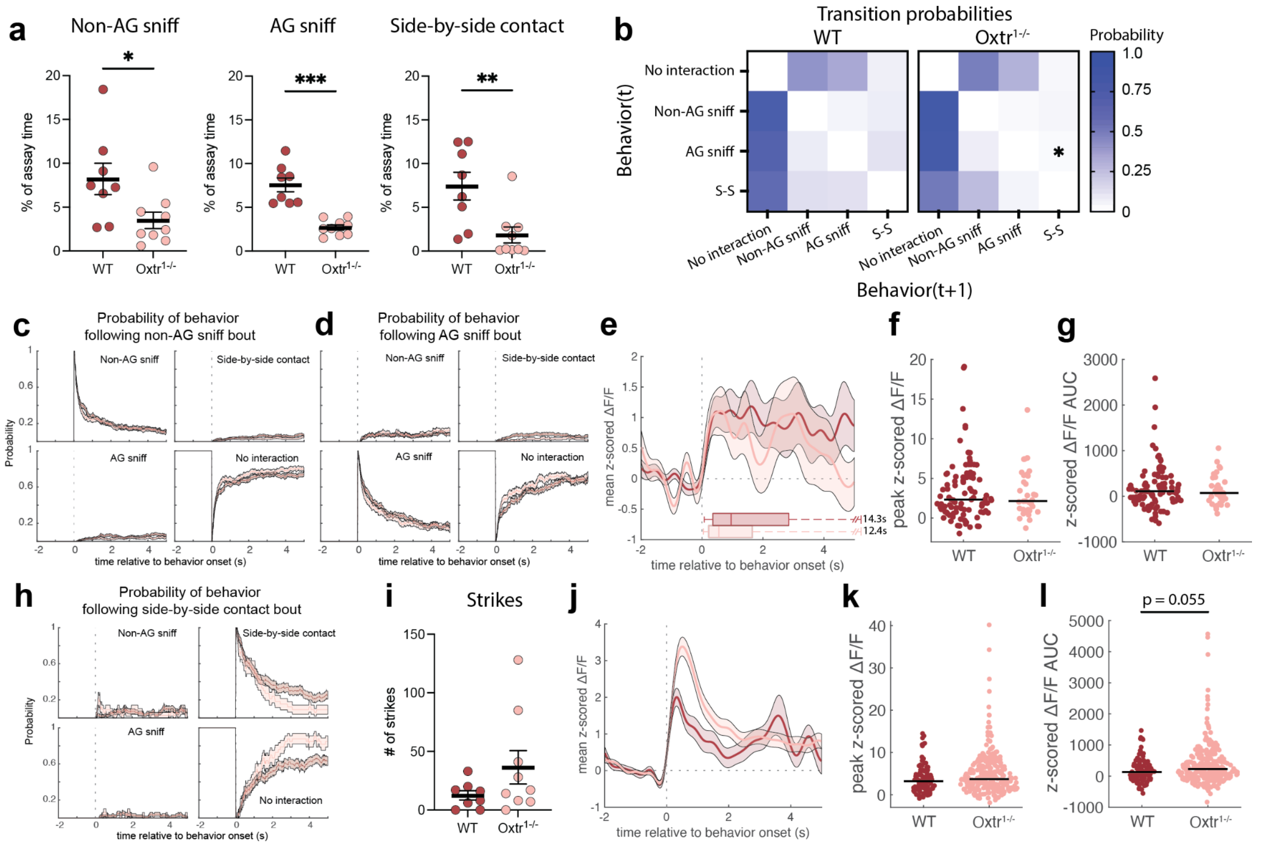


**Extended Data Figure 2: Additional data from female introductions. a,** Percent of introduction assay time spent engaged in non-anogenital (AG) sniffing, AG sniffing, or side-by-side contact (for all plots, WT n=8 voles, $Oxtr^{1-/-}$ n=9 voles). **b,** Heat maps of transition probabilities from Markov modeling of behavior. **c,** Probability of social touch behaviors following the onset of a non-AG sniff bout. Data are related to that shown in Fig. 1j. **d,** Probability of social touch behaviors following the onset of an AG sniff bout. Data are related to that shown in Fig. 1m. **e,** Mean z-scored ΔF/F (+/- s.e.m.) traces at the onset of social bouts initiated by side-by-side contact. At the base of the plot is an adjusted boxplot of the durations of the initiating behavior (WT n=95 traces, $Oxtr^{1-/-}$ n=33 traces). **f,** Peak z-scored ΔF/F values. **g,** AUC values of z-scored ΔF/F traces. **h,** Probability of social touch behaviors following the onset of a side-by-side contact bout. **i,** Number of strikes exhibited by females. **j,** Mean z-scored ΔF/F traces at the onset of strikes (WT n=93 traces/7 voles, $Oxtr^{1-/-}$ n=281 traces/8 voles). **k**, Peak z-scored ΔF/F values. **l,** AUC values of z-scored ΔF/F traces. Detailed statistics are presented in Extended Data File 1. *p<0.05, **p<0.01,

***p<0.001, ****p<0.0001. AG, anogenital; WT, wild-type; S-S, side-by-side contact; AUC, area under the curve.

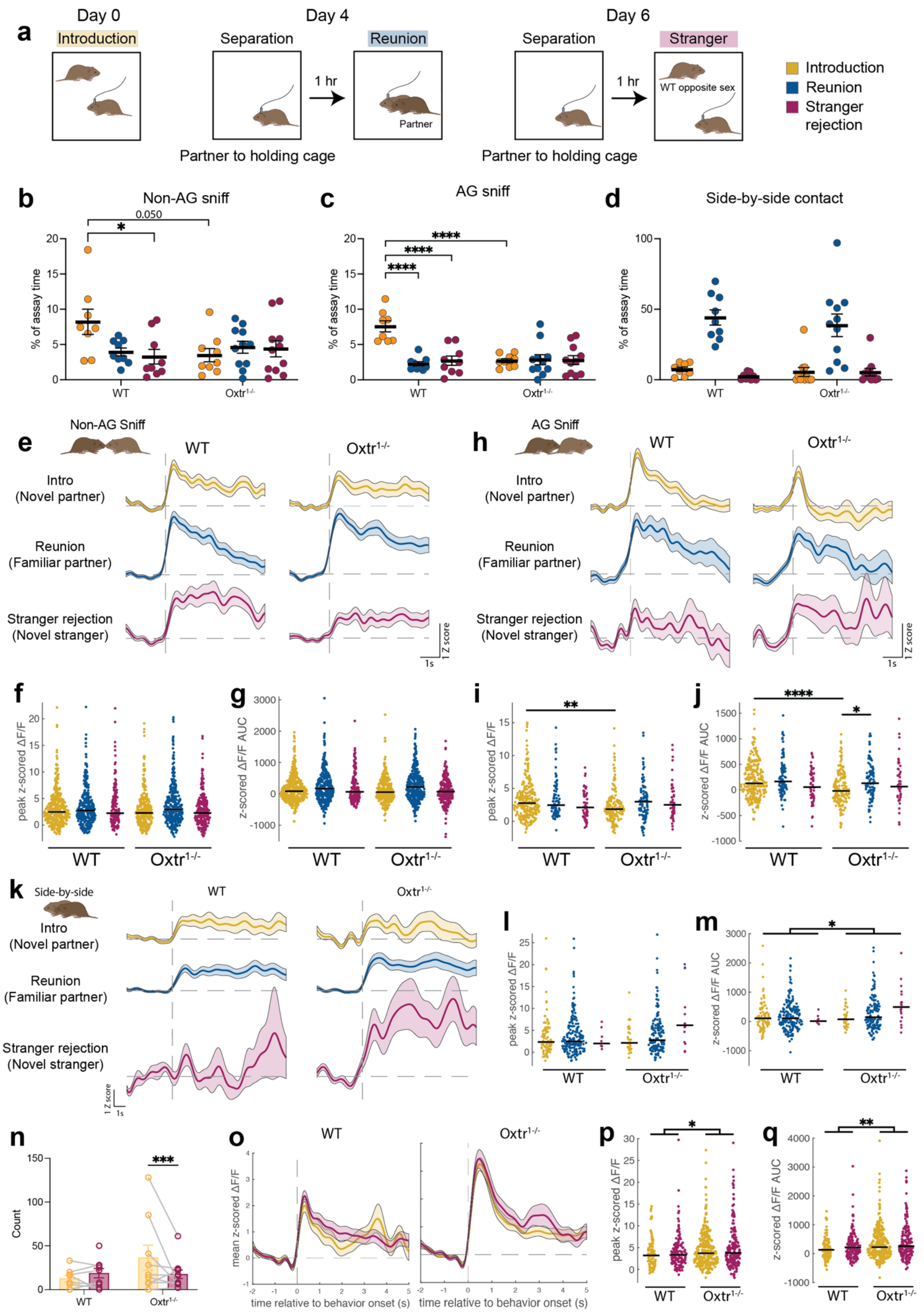

a
Day 0
Introduction
Day 4
Separation
Reunion
1 hr
Partner
Partner to holding cage
Day 6
Separation
Stranger
1 hr
WT opposite sex
Partner to holding cage
Introduction
Reunion
Stranger rejection
b
Non-AG sniff
c
AG sniff
d
Side-by-side contact
% of assay time
WT
Oxtr-/-
e
Non-AG Sniff
h
AG Sniff
Intro (Novel partner)
Reunion (Familiar partner)
Stranger rejection (Novel stranger)
1 Z score
1s
peak z-scored ΔF/F
z-scored ΔF/F AUC
k
Side-by-side
n
Count
o
mean z-scored ΔF/F
time relative to behavior onset (s)

**Extended Data Figure 3: Cross-assay analyses in females.** **a**, Schematic of introduction, reunion, and stranger rejection paradigms. **b**, Percent of assay time spent non-AG sniffing the stimulus animal across assays. **c**, Percent of time spent AG sniffing the stimulus animal across assays. **d**, Percent of time spent in side-by-side contact with the stimulus animal across assays (main effect of assay). **e,** Mean ΔF/F PETH aligned to non-AG sniffs during the introduction, reunion, and stranger rejection assays. Yellow, intro (WT n=394 traces from 8 animals; $Oxtr^{1-/-}$ n=354 traces from 9 animals); blue, reunion (WT n=326 traces from 9 animals; $Oxtr^{1-/-}$ n=237 traces from 10 animals); purple, stranger rejection (WT n=164 traces from 8 animals; $Oxtr^{1-/-}$ n=47 traces from 10 animals). **f**, Peak z-scored ΔF/F values (main effect of assay). **g**, AUC values from z-scored ΔF/F traces (main effect of assay). **h,** Mean ΔF/F PETH aligned to AG sniffs during the intro, reunion, and stranger rejection assays. (Intro: WT n=268 traces from 8 animals, $Oxtr^{1-/-}$ n=161 traces from 9 animals; Reunion: WT n=86 traces from 9 animals, $Oxtr^{1-/-}$ n=101 traces from 10 animals; Stranger rejection: WT n=54 traces from 8 animals, $Oxtr^{1-/-}$ n=47 traces from 10 animals). **i**, Peak ΔF/F values. **j**, AUC values from z-scored ΔF/F traces. **k**, Mean ΔF/F traces surrounding social bouts initiated with side-by-side contact during the introduction, reunion, and stranger rejection assays (Intro: WT n=95 traces from 8 animals; $Oxtr^{1-/-}$ n=33 traces from 6 animals; Reunion: WT n=190 traces from 9 animals; $Oxtr^{1-/-}$ n=148 traces from 11 animals; Stranger rejection: WT n=8 traces from 3 animals; $Oxtr^{1-/-}$ n=18 traces from 6 animals). **l**, Peak ΔF/F values. **m**, AUC values from z-scored ΔF/F traces (significant main effect of genotype). **n**, Number of strikes in the introduction and stranger rejection assays. **o,** Mean ΔF/F traces surrounding strikes. (Intro: WT n=93 traces from 6 animals; $Oxtr^{1-/-}$ n=278 traces from 8 animals; Stranger rejection: WT n=152 traces from 8 animals; $Oxtr^{1-/-}$ n=182 traces from 11 animals). **p**, Peak ΔF/F values (significant main effect of genotype). **q**, AUC values from z-scored ΔF/F traces (significant main effect of genotype). Detailed statistics are presented in Extended Data File 1. $^{*}p<0.05$, $^{**}p<0.01$, $^{***}p<0.001$, $^{****}p<0.0001$. WT, wild-type; AUC, area under the curve; AG, anogenital; PETH, peri-event time histogram.

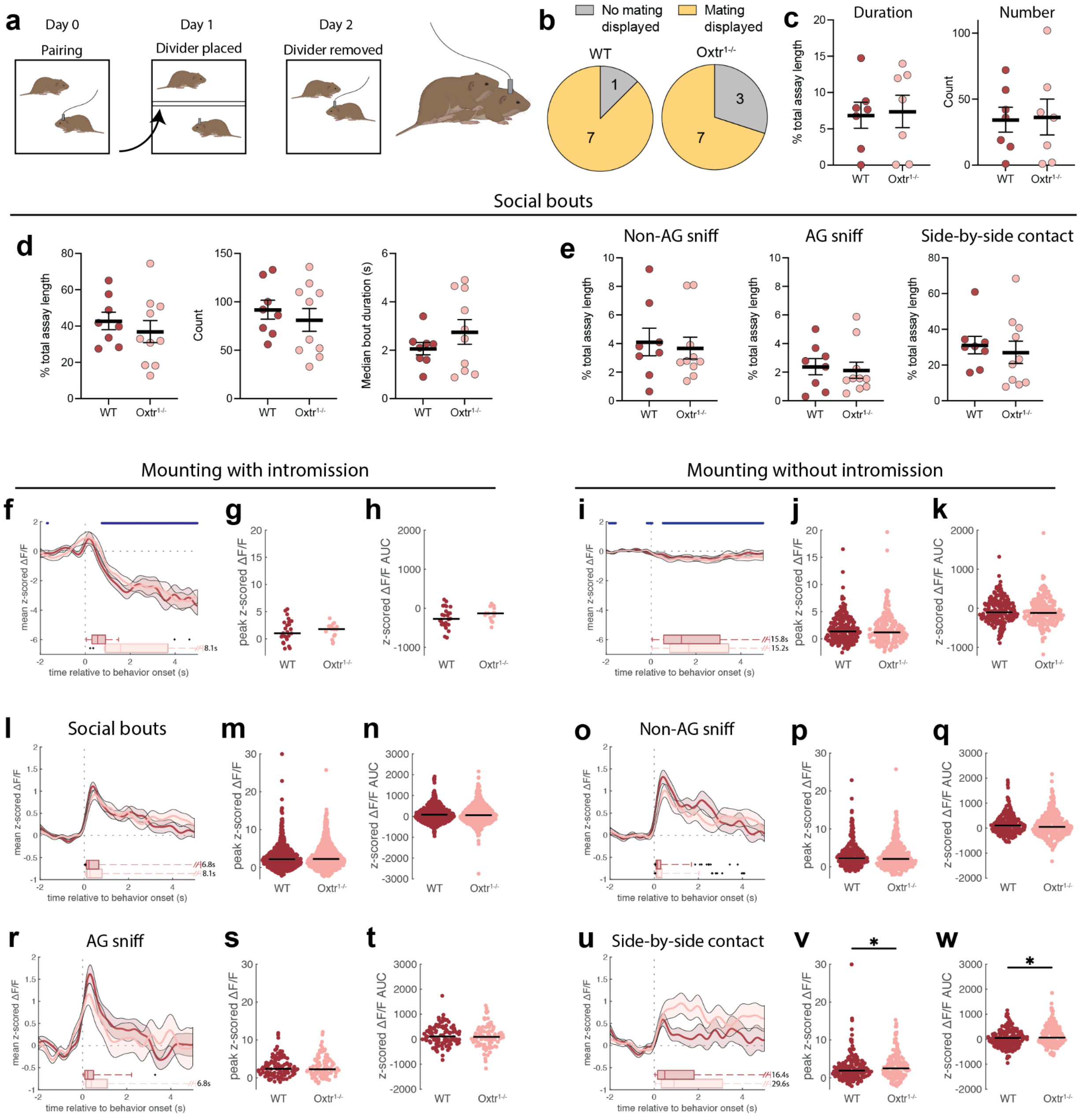


**Extended Data Figure 4: Additional data from female timed mating assays. a,** Timed mating procedure. Roughly 24 hours after introductions, we placed a clear plastic barrier with small holes in the cage to divide the animals into two compartments and to allow limited social contact. The following day, the focal animal was habituated to the patch cable, the barrier was removed, and behavior was recorded for 30 minutes. **b,** Mounting and intromission were scored when a female received mounting or intromission. Pie charts show the proportion of females that received mating behavior. **c,** Left, percent of assay time spent engaged in mating behavior, plotting only the females that received any mating attempts. Right, number of mating bouts, plotting only females that received any mating attempts (n=7 per group). **d**, Quantification of social bouts. Left, total amount of assay time spent engaged in social bouts. Middle, number of social bouts. Right, median duration of social bouts (for subsequent plots, WT n=8 voles, $Oxtr^{1-/-}$ n=10 voles). **e,** Percent of timed mating assay time spent engaged in non-anogenital (AG) sniffing, AG sniffing, or side-by-side contact. **f,** Mean z-scored ΔF/F traces at the onset of mounts that transition to intromission. Blue bars at the top of the plot indicate time points at which the combined WT and $Oxtr^{1-/-}$ mean z-scored ΔF/F is significantly less than 0. At the base of the plot is an adjusted boxplot of the durations of the initiating behavior (WT n=26 traces/3 voles, $Oxtr^{1-/-}$ n=16 traces/3 voles). **g,** Peak z-scored ΔF/F values. **h,** AUC values of z-scored ΔF/F traces. **i,** Mean z-scored ΔF/F traces at the onset of mounts that do not transition to intromission (WT n=233 traces/7 voles, $Oxtr^{1-/-}$ n=244 traces/7 voles). **j,** Peak z-scored ΔF/F values. **k,** AUC values of z-scored ΔF/F traces. **l,** Mean z-scored ΔF/F traces at the onset of social bouts (WT n=621 traces, $Oxtr^{1-/-}$ n=663 traces). **m,** Peak z-scored ΔF/F values. **n,** AUC values of z-scored ΔF/F traces. **o,** Mean z-scored ΔF/F traces at the onset of social bouts initiated by non-AG sniffing (WT n=278 traces, $Oxtr^{1-/-}$ n=378 traces). **p,** Peak z-scored ΔF/F values. **q,** AUC values of z-scored ΔF/F traces. **r,** Mean z-scored ΔF/F traces at the onset of social bouts initiated by AG sniffing (WT n=118 traces, $Oxtr^{1-/-}$ n=103 traces). **s,** Peak z-scored ΔF/F values. **t,** AUC values of z-scored ΔF/F traces. **u,** Mean z-scored ΔF/F traces at the onset of social bouts initiated by side-by-side contact (WT n=224 traces, $Oxtr^{1-/-}$ n=180 traces). **v,** Peak z-scored ΔF/F values. **w,** AUC values of z-scored ΔF/F traces. Detailed statistics are presented in Extended Data File 1. *$p<0.05$. WT, wild-type; AUC, area under the curve; AG, anogenital.

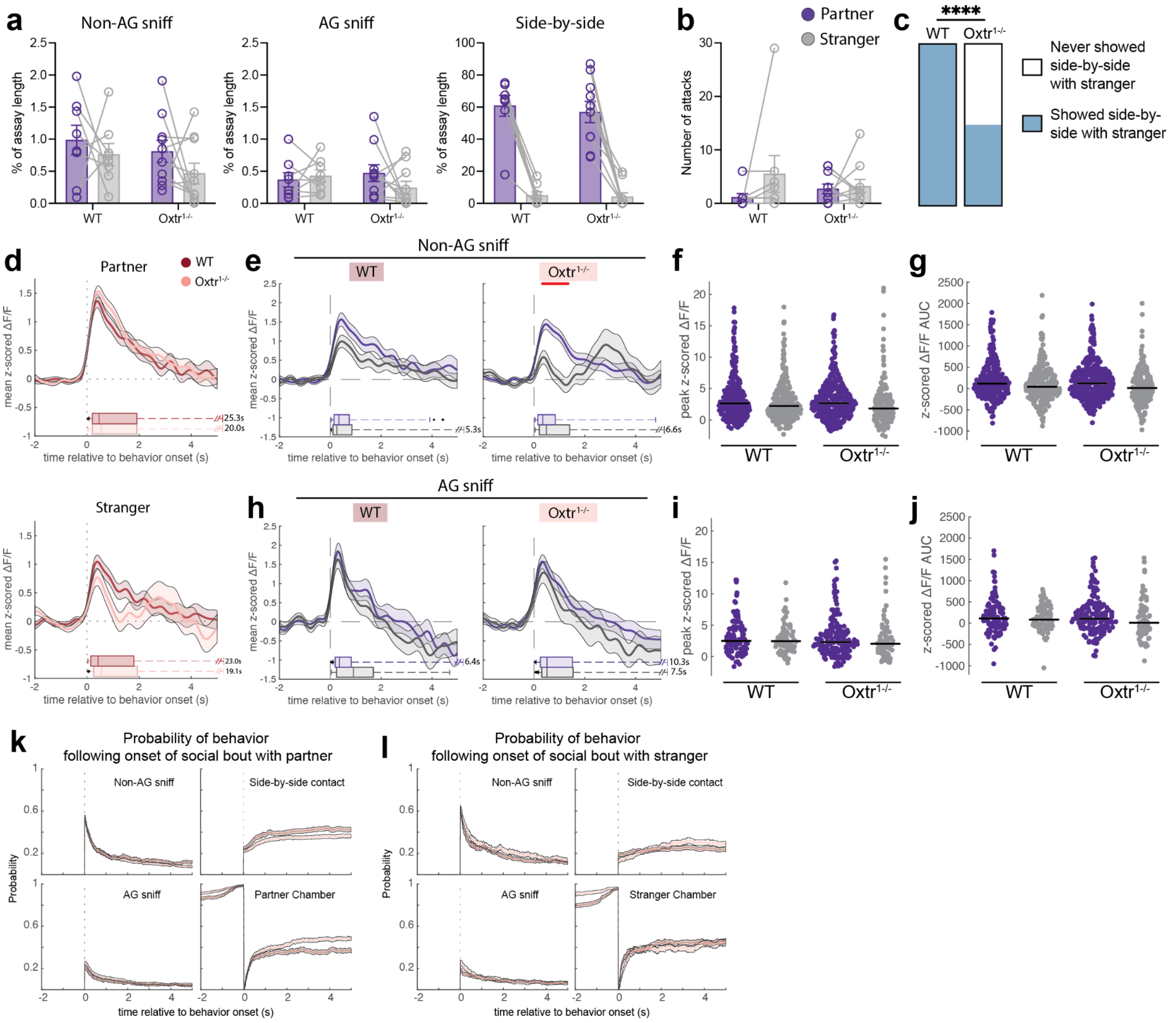


**Extended Data Figure 5: Additional data from female partner preference tests. a,** Percent of PPT assay time spent engaged in non-anogenital (AG) sniffing, AG sniffing, or side-by-side contact. Note the difference in the y-axis for side-by-side contact (for all plots, WT n=8 voles, Oxtr$^{1-/-}$ n=10 voles). **b,** Number of attacks against the partner or stranger male. **c**, Proportion of females that displayed at least one bout of side-by-side contact with the stranger male. **d,** Mean z-scored ΔF/F (+/- s.e.m.) traces at the onset of social bouts with either the partner (top) or stranger (bottom). Data are the same as those plotted Fig. 2f. At the base of the plot is an adjusted boxplot of the durations of the initiating behavior. **e,** Mean z-scored ΔF/F traces at the onset of social bouts initiated by non-AG sniffing. The red line indicates time points at which mean z-scored

$\Delta F/F$ differs between partner and and stranger ($WT_{Partner}$ n=291 traces, $WT_{Stranger}$ n=276 traces, $Oxtr^{1-/-}_{Partner}$ n=383 traces, $Oxtr^{1-/-}_{Stranger}$ n=188 traces). **f,** Peak z-scored $\Delta F/F$ values (significant main effects of stimulus animal and genotype). **g,** AUC values of z-scored $\Delta F/F$ traces (significant main effect of stimulus animal). **h,** Mean z-scored $\Delta F/F$ (+/- s.e.m.) traces at the onset of social bouts initiated by AG sniffing ($WT_{Partner}$ n=113 traces, $WT_{Stranger}$ n=83 traces, $Oxtr^{1-/-}_{Partner}$ n=185 traces, $Oxtr^{1-/-}_{Stranger}$ n=77 traces). **i,** Peak z-scored $\Delta F/F$ values. **j,** AUC values of z-scored $\Delta F/F$ traces. **k,** Probability of social touch behaviors following the onset of a social bout with a partner. Data are related to that shown in d (Partner). **l,** Probability of social touch behaviors following the onset of a social bout with a stranger. Data are related to that shown in d (Stranger). Detailed statistics are presented in Extended Data File 1. $^*p<0.05$, $^{**}p<0.01$, $^{***}p<0.001$, $^{****}p<0.0001$. AG, anogenital; WT, wild-type; AUC, area under the curve.

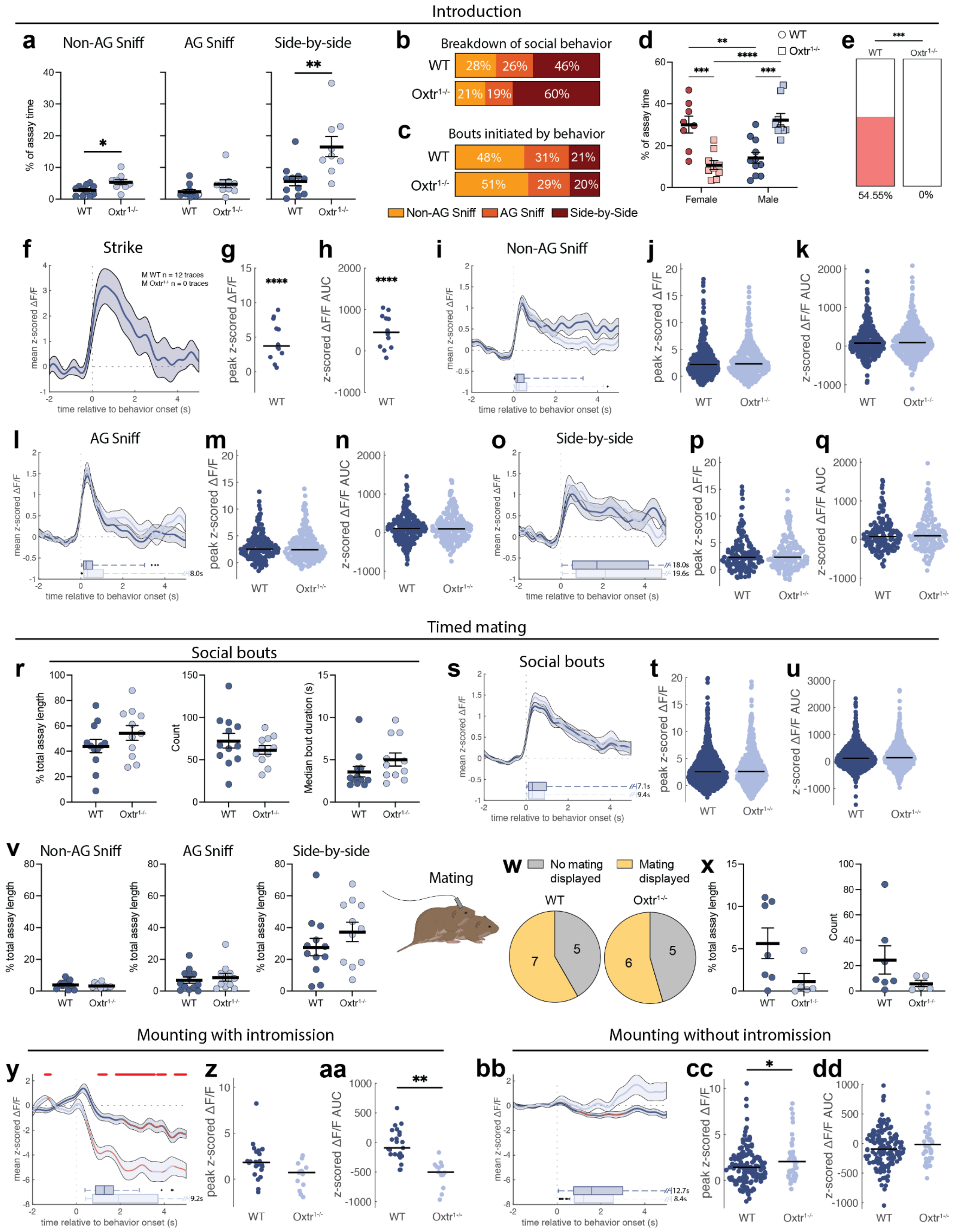


**Extended Data Figure 6: Additional data from male introduction and timed mating assays.**

**a,** Percent of introduction assay time spent engaged in non-anogenital (AG) sniffing, AG sniffing, or side-by-side contact (for plots h-q, WT n=11, $Oxtr^{1-/-}$ n=9 voles). **b,** Breakdown of social contact by the percentage of time engaged in anogenital (AG) sniffing, non-AG sniffing, and side-by-side contact. **c,** Percentages of social bouts initiated with non-AG sniff, AG sniff, or side-by-side contact. **d**, Percent of introduction assay time spent engaged in social bouts (female WT n=8, female $Oxtr^{1-/-}$ n=9, male WT n=11, male $Oxtr^{1-/-}$ n=9). **e,** Proportion of males displaying any offensive strikes during the introduction. **f,** Mean z-scored ΔF/F traces at the onset of strikes (WT n=12). **g**, Peak z-scored ΔF/F values. **h,** AUC values of z-scored ΔF/F traces. **i,** Mean z-scored ΔF/F traces at the onset of social bouts initiated by non-AG sniffing. At the base of the plot is an adjusted boxplot of the durations of the initiating behaviors (WT n=358 traces, $Oxtr^{1-/-}$ n=435 traces). **j,** Peak z-scored ΔF/F values. **k,** AUC values of z-scored ΔF/F traces. **l,** Mean z-scored ΔF/F traces at the onset of social bouts initiated by AG sniffing (WT n=219 traces, $Oxtr^{1-/-}$ n=255 traces). **m,** Peak z-scored ΔF/F values. **n,** AUC values of z-scored ΔF/F traces. **o,** Mean z-scored ΔF/F traces at the onset of social bouts initiated by side-by-side contact (WT n=171 traces, $Oxtr^{1-/-}$ n=171 traces). **p,** Peak z-scored ΔF/F values. **q,** AUC values of z-scored ΔF/F traces. **r**, Quantification of social bouts during the timed mating assay. Left, total amount of assay time spent engaged in social bouts. Middle, number of social bouts. Right, median duration of social bouts (for plots r-v, WT n=12, $Oxtr^{1-/-}$ n=11 voles). **s,** Mean z-scored ΔF/F traces at the onset of social bouts (WT n=782 traces, $Oxtr^{1-/-}$ n=640 traces). **t,** Peak z-scored ΔF/F values. **u,** AUC values of z-scored ΔF/F traces. **v,** Percent of timed mating assay time spent engaged in non-AG sniffing, AG sniffing, or side-by-side contact. **w,** Proportions of males that exhibited mating behavior. **x,** Left, percent of assay time spent engaged in mating behavior, plotting only the males that exhibited any mating. Right, number of mating bouts, plotting only males that exhibited any mating (WT n=7, $Oxtr^{1-/-}$ n=6 voles). **y,** Mean z-scored ΔF/F traces at the onset of mounts that transition to intromission. The red line indicates time points at which mean z-scored ΔF/F differs between WT and $Oxtr^{1-/-}$ males. The orange asterisks on the mean PETH lines indicate time points at which that mean z-scored ΔF/F is significantly different from 0 ΔF/F (WT n=23 traces/3 voles,

Oxtr$^{1-/-}$ n=13 traces/2 voles). **z,** Peak z-scored ΔF/F values. **aa,** AUC values of z-scored ΔF/F traces. **bb,** Mean z-scored ΔF/F traces at the onset of mounts that do not transition to intromission (WT n=157 traces/7 voles, Oxtr$^{1-/-}$ n=42 traces/6 voles). **cc,** Peak z-scored ΔF/F values. **dd,** AUC values of z-scored ΔF/F traces. Detailed statistics are presented in Extended Data File 1. *p<0.05, **p<0.01, ****p<0.0001. WT, wild-type; AUC, area under the curve; AG, anogenital.

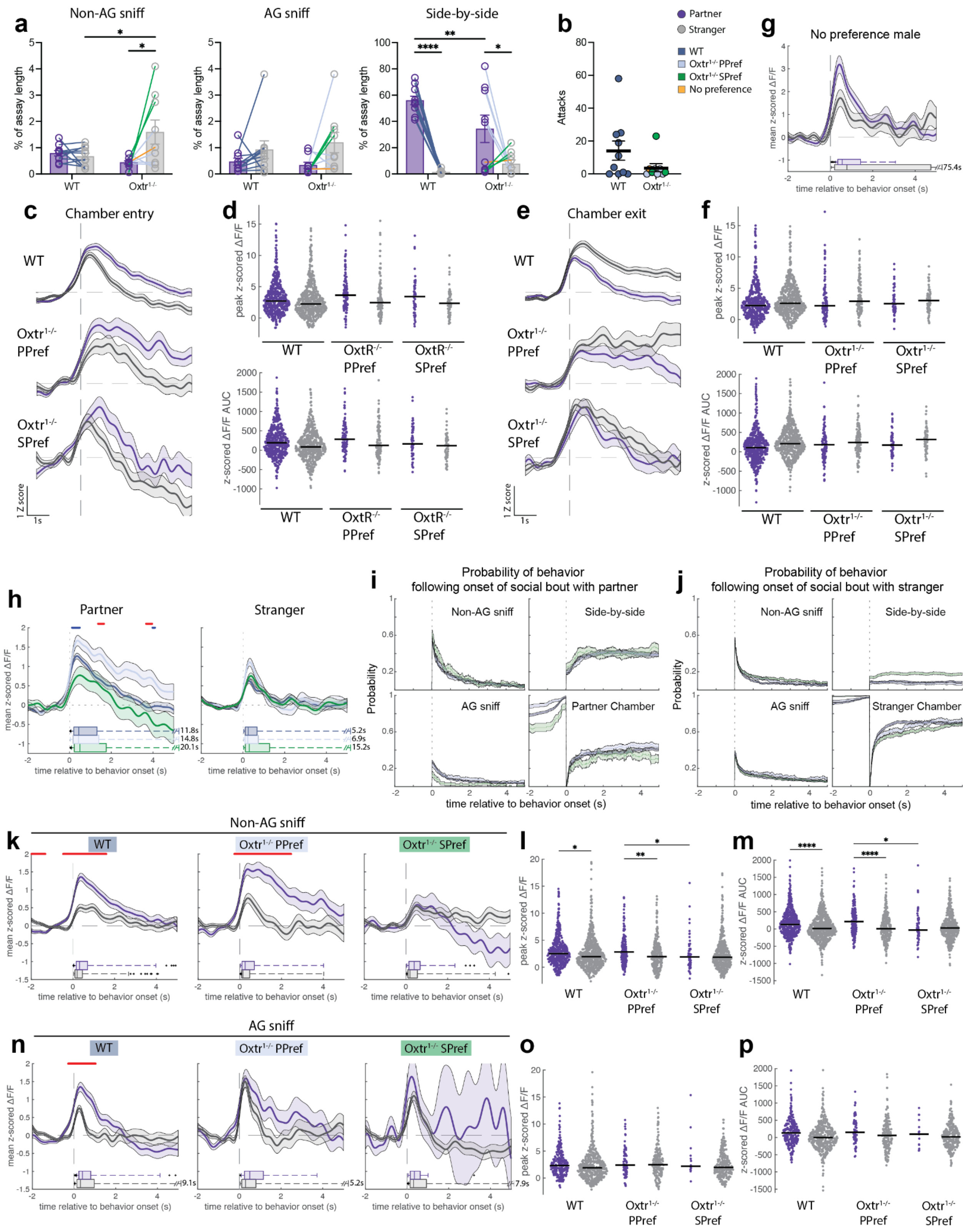


**Extended Data Figure 7: Additional data from male PPT. a,** Percent of PPT assay time spent engaged in non-anogenital (AG) sniffing, AG sniffing (significant main effect of stimulus animal), or side-by-side contact. Note the difference in y axis for side-by-side contact (WT n=10, $Oxtr^{1-/-}$ n=9 voles). **b,** Male attacks against the stranger female. **c,** Mean ΔF/F traces surrounding entries to either the partner chamber or stranger chamber (for this and subsequent plots: WT n=10 voles, $Oxtr^{1-/-}$ PPref n=5 voles, $Oxtr^{1-/-}$ SPref n=3 voles; $WT_{Partner}$ n=485, $WT_{Stranger}$ n=542; $Oxtr^{1-/-}$ $PPref_{Partner}$ n=107, $Oxtr^{1-/-}$ $PPref_{Stranger}$ n=108; $Oxtr^{1-/-}$ $SPref_{Partner}$ n=56, $Oxtr^{1-/-}$ $SPref_{Stranger}$ n=62 traces). **d,** Top, peak z-scored ΔF/F values with median value overlaid (significant main effect of stimulus animal chamber). Bottom, AUC values from z-scored ΔF/F traces (significant main effect of stimulus animal chamber). **e**, Mean ΔF/F traces surrounding exits from either the partner chamber or stranger chamber ($WT_{Partner}$ n=490, $WT_{Stranger}$ n=563; $Oxtr^{1-/-}$ $PPref_{Partner}$ n=105, $Oxtr^{1-/-}$ $PPref_{Stranger}$ n=116; $Oxtr^{1-/-}$ $SPref_{Partner}$ n=51, $Oxtr^{1-/-}$ $SPref_{Stranger}$ n=64 traces). **f,** Top, peak z-scored ΔF/F values (significant main effect of stimulus animal chamber). Bottom, AUC values from z-scored ΔF/F traces (significant main effect of stimulus animal chamber). **g**, Mean ΔF/F traces from the male exhibiting no partner or stranger preference surrounding social bouts with the partner (purple) or stranger (gray). At the base of the plot is an adjusted boxplot of durations of the initiating behaviors (partner n=41, stranger n=56 traces from 1 male). **h,** Mean z-scored ΔF/F traces at the onset of social bouts with a partner (left) or stranger (right). Data are the same as those plotted in Fig. 3k. Red bars above the plot indicate time points at which mean z-scored ΔF/F significantly differs between WT and partner-preferring $Oxtr^{1-/-}$ males. Blue bars indicate time points at which partner-preferring and stranger-preferring $Oxtr^{1-/-}$ males differ ($WT_{Partner}$ n=932 traces, $Oxtr^{1-/-}$ $PPref_{Partner}$ n=296 traces, $Oxtr^{1-/-}$ $SPref_{Partner}$ n=111 traces; $WT_{Stranger}$ n=1193 traces, $Oxtr^{1-/-}$ $PPref_{Stranger}$ n=500 traces, $Oxtr^{1-/-}$ $SPref_{Stranger}$ n=821 traces). **i,** Probability of social touch behaviors following the onset of a social bout with a partner. Data are related to those shown in h (Partner). **j,** Probability of social touch behaviors following the onset of a social bout with a stranger. Data are related to those shown in h (Stranger). **k,** Mean z-scored ΔF/F traces at the onset of non-AG sniffs of the partner or stranger. The red line indicates time points at which mean z-scored ΔF/F differs between partner and stranger-related activity ($WT_{Partner}$ n=508 traces,

$WT_{Stranger}$ n=647 traces; Oxtr$^{1-/-}$ $PPref_{Partner}$ n=166 traces, Oxtr$^{1-/-}$ $PPref_{Stranger}$ n=277 traces; Oxtr$^{1-/-}$ $SPref_{Partner}$ n=68 traces, Oxtr$^{1-/-}$ $SPref_{Stranger}$ n=434 traces). **l,** Peak z-scored ΔF/F values. **m,** AUC values of z-scored ΔF/F traces. **n,** Mean z-scored ΔF/F traces at the onset of social bouts initiated by AG sniffing ($WT_{Partner}$ n=252 traces, $WT_{Stranger}$ n=434 traces; Oxtr$^{1-/-}$ $PPref_{Partner}$ n=71 traces, Oxtr$^{1-/-}$ $PPref_{Stranger}$ n=185 traces; Oxtr$^{1-/-}$ $SPref_{Partner}$ n=15 traces, Oxtr$^{1-/-}$ $SPref_{Stranger}$ n=268 traces). **o,** Peak z-scored ΔF/F values. **p,** AUC values of z-scored ΔF/F traces (significant main effect of stimulus animal). Detailed statistics are presented in Extended Data File 1. *p<0.05, **p<0.01, ***p<0.001, ****p<0.0001. AG, anogenital; WT, wild-type; PPref, Oxtr$^{1-/-}$ partner-preferring; SPref, Oxtr$^{1-/-}$ stranger-preferring; AUC, area under the curve.

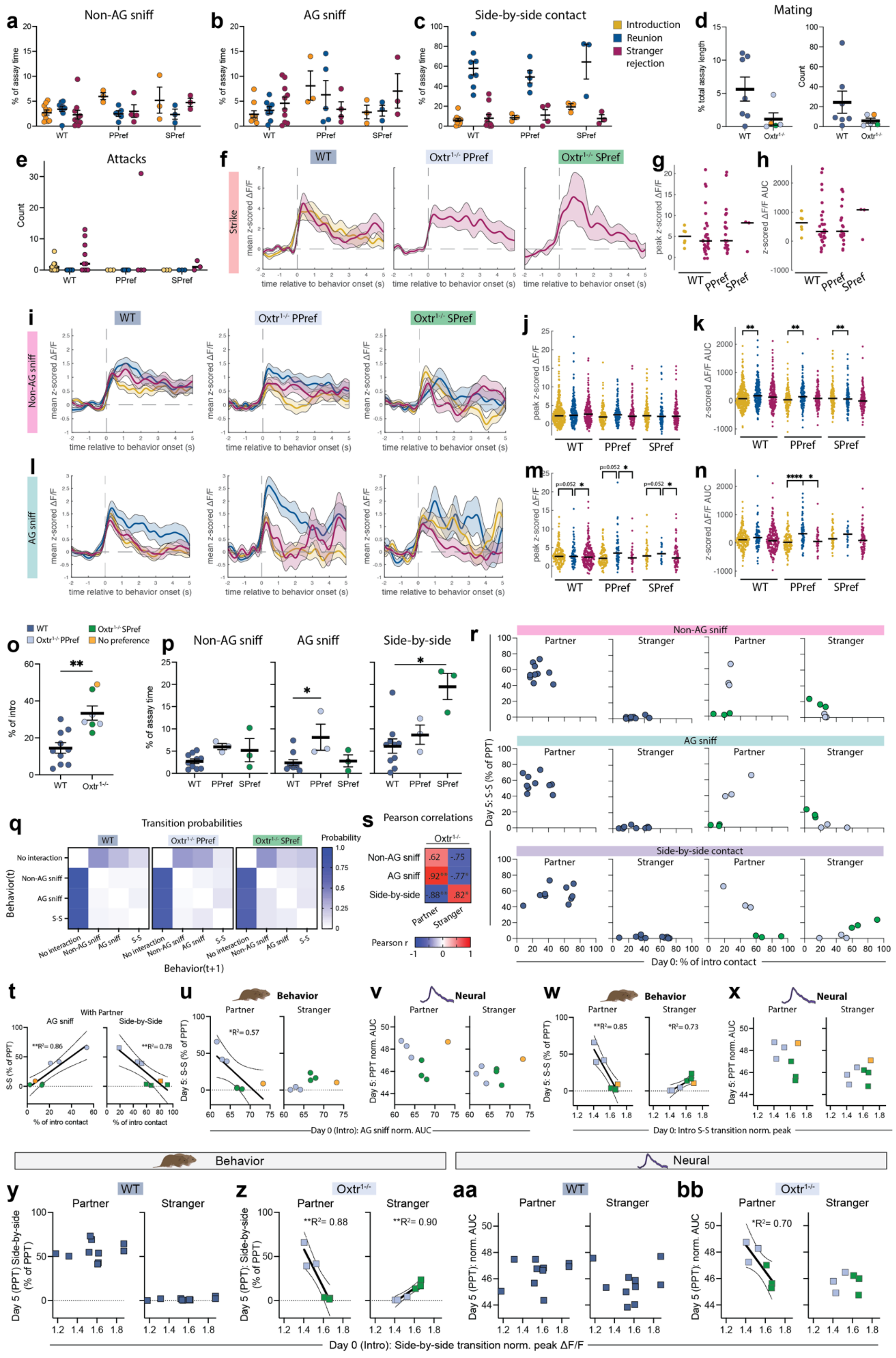

a Non-AG sniff
b AG sniff
c Side-by-side contact
Introduction
Reunion
Stranger rejection
d Mating
e Attacks
f WT Oxtr1-/- PPref Oxtr1-/- SPref
Strike
time relative to behavior onset (s)
g
h
i WT Oxtr1-/- PPref Oxtr1-/- SPref
Non-AG sniff
j
k
l AG sniff
m
n
o
p Non-AG sniff AG sniff Side-by-side
q Transition probabilities
No interaction
Non-AG sniff
AG sniff
S-S
Behavior(t)
Behavior(t+1)
Probability
s Pearson correlations
r Non-AG sniff
AG sniff
Side-by-side contact
Partner Stranger
Day 5: S-S (% of PPT)
Day 0: % of intro contact
t With Partner
u Behavior
v Neural
w Behavior
x Neural
Day 0 (Intro): AG sniff norm. AUC
Day 0: Intro S-S transition norm. peak
Behavior
Neural
y WT
z Oxtr1-/-
aa WT
bb Oxtr1-/-
Day 0 (Intro): Side-by-side transition norm. peak ΔF/F

**Extended Data Figure 8: Additional data related to Figure 4. a,** Percent of assay time spent non-AG sniffing the stimulus animal across assays (for subsequent plots unless otherwise noted, $WT_{Intro}$ n=10, $WT_{Reunion}$ n=8, $WT_{Stranger}$ n=10, $Oxtr^{1-/-}$ $PPref_{Intro}$ n=3, $Oxtr^{1-/-}$ $PPref_{Reunion}$ n=5, $Oxtr^{1-/-}$ $PPref_{Stranger}$ n=4, $Oxtr^{1-/-}$ $SPref_{Intro}$ n=3, $Oxtr^{1-/-}$ $SPref_{Reunion}$ n=3, $Oxtr^{1-/-}$ $SPref_{Stranger}$ n=3). **b,** Percent of time spent AG sniffing the stimulus animal across assays. **c,** Percent of time spent in side-by-side contact with the stimulus animal across assays (significant main effect of assay). **d,** Left, percent of assay time spent engaged in mating behavior, plotting only the males that exhibited any mating. Right, number of mating bouts, plotting only males that exhibited any mating (WT n=7, $Oxtr^{1-/-}$ n=5 voles). **e,** Number of strikes in the pairing, reunion, and stranger rejection assays. **f,** Mean ΔF/F traces surrounding strikes (WT Intro n=6 traces from 5 animals; WT Stranger rejection n=30 traces from 6 animals; $Oxtr^{1-/-}$ PPref Stranger rejection n=22 traces from 1 animal; $Oxtr^{1-/-}$ SPref Stranger rejection n=3 traces from 2 animals). **g,** Peak ΔF/F values. **h**, AUC values from z-scored ΔF/F traces. **i,** Mean z-scored ΔF/F traces surrounding non-AG sniffs during the introduction, reunion, and stranger rejection assays. Yellow, intro (WT n=338 traces; $Oxtr^{1-/-}$ PPref n=137 traces; $Oxtr^{1-/-}$ SPref n=123 traces); blue, reunion (WT n=193 traces; $Oxtr^{1-/-}$ PPref n=124 traces; $Oxtr^{1-/-}$ SPref n=87 traces); purple, stranger rejection (WT n=220 traces; $Oxtr^{1-/-}$ PPref n=115 traces; $Oxtr^{1-/-}$ SPref n=130 traces). **j,** Peak z-scored ΔF/F values with median value overlaid (significant main effect of group). **k,** AUC values from z-scored ΔF/F traces (significant main effects of group and assay). **l,** Mean ΔF/F traces surrounding AG sniffs during the intro, reunion, and stranger rejection assays. (Intro: WT n=187 traces; $Oxtr^{1-/-}$ PPref n=131 traces; $Oxtr^{1-/-}$ SPref n=42 traces; Reunion WT n=77 traces from 7 animals; $Oxtr^{1-/-}$ PPref n=65 traces from 4 animals; $Oxtr^{1-/-}$ SPref n=20 traces from 3 animals; Stranger rejection: WT n=215 traces; $Oxtr^{1-/-}$ PPref n=34 traces; $Oxtr^{1-/-}$ SPref n=81 traces). **m,** Peak ΔF/F values (significant main effect of assay). **n,** AUC values from z-scored ΔF/F traces. **o,** Percent of time engaged in social interaction with a newly partnered female during the introduction, plotting only animals from which we successfully collected both intro and PPT data (including the single $Oxtr^{1-/-}$ male with no preference). Individual animals with median overlaid and colored according to PPT behavior profile (WT n=10, $Oxtr^{1-/-}$ n=7). **p,** Percent of introduction assay time spent engaged in non-anogenital (AG) sniffing, AG sniffing, or side-by-side contact. **q,** Heat maps of transition probabilities from Markov modeling of behavior. **r,** Correlation plots between behavior exhibited during the intro and side-by-side contact with either the partner or stranger during the PPT (X axis, percent of social contact; Y axis, percent of PPT time). These plots are the raw data of the heat maps shown in Fig. 4d. **s,** Heat map of correlations between behavior exhibited during the intro and side-by-side contact with either the partner or stranger during the PPT. Data are related to Fig. 4d, but include the single animal with no huddling preference. **t-x,** Plots related to Fig. 4e,m,o,v,x but including the single $Oxtr^{1-/-}$ male with no huddle preference. **t,** Linear regression of intro behavior to PPT behavior. X-axis: Percent of social touch during the introduction spent AG sniffing (left, teal shading) or in side-by-side contact (right, purple shading). Y-axis: Percent of time spent in side-by-side contact (S-S) with the partner during the PPT. **u-v,** Linear regressions of mean normalized AUC at the onset of AG sniff bouts, averaged by animal, during the introduction and PPT side-by-side contact (g) or PPT mean normalized AUC surrounding social bouts by animal (h). **w-x,** Linear regressions of normalized peak $\Delta F/F$ surrounding sniff to side-by-side contact transitions during the introduction and PPT side-by-side contact (h) or normalized AUC surrounding social bouts (i). **y-bb,** Linear regressions comparing side-by-side-related neural activity and PPT behavior or neural data. X axis: normalized peak $\Delta F/F$ at the onset of sniff to side-by-side contact transitions during the introduction, averaged by animal. Y axis: PPT side-by-side contact (y-z) or PPT normalized AUC surrounding social bouts, averaged by animal (aa-bb). Detailed statistics are presented in Extended Data File 1. *p<0.05, **p<0.01, ***p<0.001, ****p<0.0001. WT, wild-type; AG, anogenital; PPref, partner-preferring $Oxtr^{1-/-}$ male; SPref, stranger-preferring $Oxtr^{1-/-}$ male; PPT, partner preference test; AUC, area under the curve.

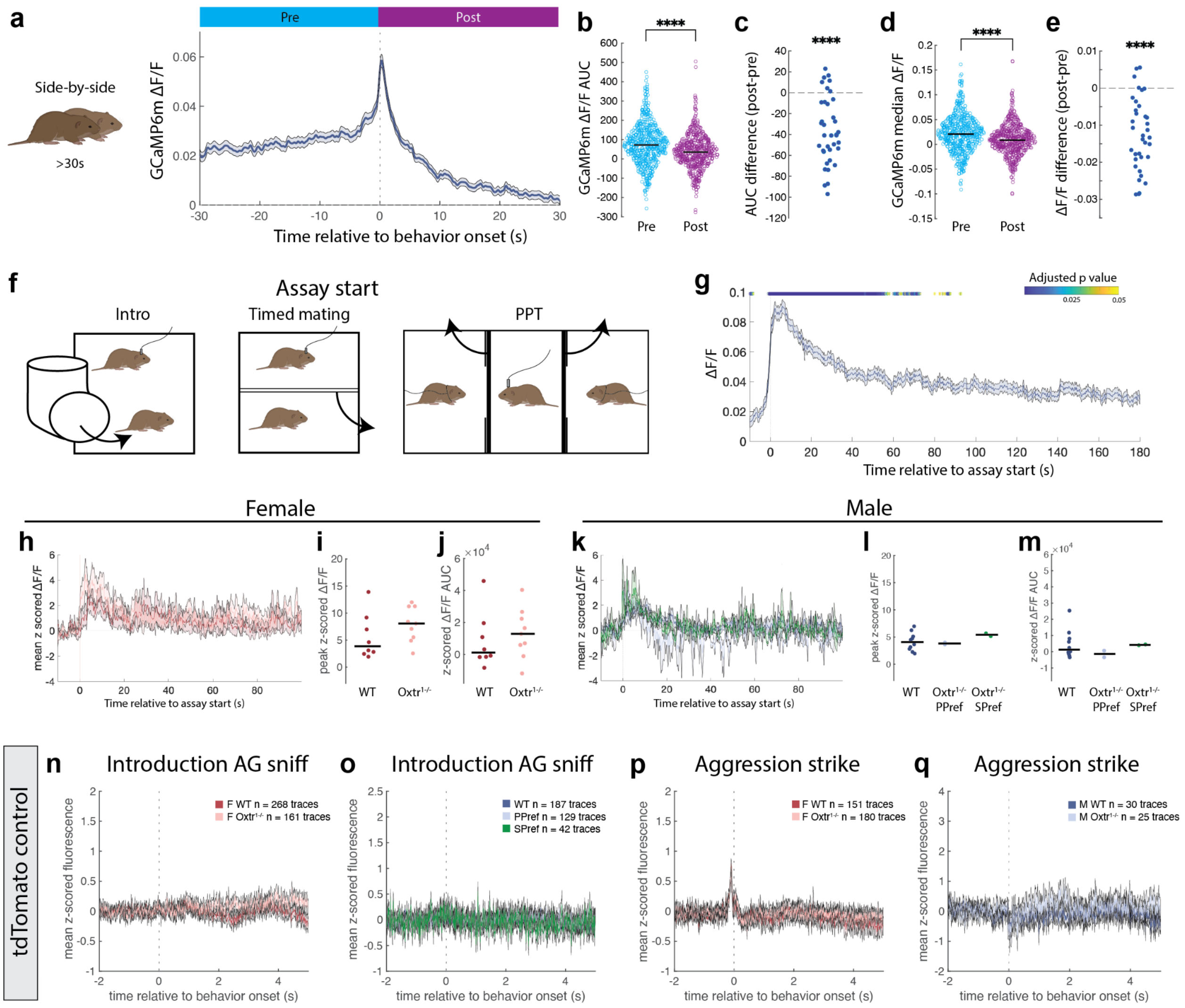

**Extended Data Figure 9: Dynamics of NAc calcium activity during periods of rest and at assay start. a,** GCaMP6m ΔF/F traces were extracted surrounding bouts of side-by-side contact lasting 30 seconds or more (prolonged huddling), during which time most animals rested or slept. Plotted is the mean of GCaMP6m ΔF/F traces (n=560 traces from 37 animals). We compared area under the curve (AUC) and median ΔF/F values from the pre-huddle period (blue, -30 to 0 seconds) to the post-huddle period (purple, 0 to 30 seconds). **b,** GCaMP6m ΔF/F AUC from the pre- vs. post-huddle period. **c,** Individual animal difference between median pre-huddle AUC and post-huddle AUC. **d,** Median GCaMP6m ΔF/F from the pre- vs. post-huddle period. **e,** Individual animal difference between median pre-huddle ΔF/F and post-huddle ΔF/F. **f,** For assays conducted in the home cage, the stimulus animal was transferred to the cage. “Assay start” was defined as the moment the stimulus was placed into the cage. Note: For timed matings and PPTs, assay start was defined as the moment the barriers between animals were removed. **g,** Mean of GCaMP6m $\Delta F/F$ traces from all animals and all assays at the time of assay start (n=192). Asterisks above the plot are time points where mean $\Delta F/F$ is significantly different from the pre-assay (-10s to 0s) mean $\Delta F/F$ value of 0.0237 (Benjamini-Yekutieli correction applied to the resulting p values). **h-m,** Assay start mean $\Delta F/F$ (h,k) and metrics from the introduction assay (h: WT n=8, $Oxtr^{1-/-}$ n=9; k: WT n=10, $Oxtr^{1-/-}$ PPref n=2; $Oxtr^{1-/-}$ SPref n=2). Traces were z scored to the -10 to 0 second period, and peak $\Delta F/F$ and AUC were calculated from the period between 0 to 100 seconds. (i) and (l): Peak z-scored $\Delta F/F$ values. (j) and (m): AUC of $\Delta F/F$ traces. **n-o,** Control tdTomato signal (plotted as mean +/- s.e.m.) at the onset of anogenital (AG) sniff bouts during the introduction in females (n: WT n=268 traces/8 voles, $Oxtr^{1-/-}$ n=161 traces/9 voles) and males (o: WT n=187 traces/11 voles, $Oxtr^{1-/-}$ PPref n=129 traces/5 voles; $Oxtr^{1-/-}$ SPref n=42 traces/3 voles). **p-q,** Control tdTomato signal at the onset of strikes during the stranger rejection assay in females (p: WT n=151 traces/8 voles, $Oxtr^{1-/-}$ n=180 traces/11 voles) and males (q: WT n=30 traces/6 voles, $Oxtr^{1-/-}$ n=25 traces/3 voles). Notably, there was a consistent increase in tdTomato signal immediately prior to strikes in females but no fluctuations following strike onset. Detailed statistics are presented in Extended Data File 1. ****p<0.0001. AUC, area under the curve; AG, anogenital; WT, wild-type.
